# Time-dependent Glucocorticoid-Induced Transcriptomic Changes in Human Trabecular Meshwork and Schlemm’s Canal

**DOI:** 10.64898/2025.12.31.696882

**Authors:** Sudeep Mehrotra, Haven Jeanneret, Kristin Perkumas, Renee Liu, Jyoti Lama, Katie Huynh, Ananya Mukundan, Hilary Scott, Atitaya Apivatthakakul, Janey L. Wiggs, Lucia Sobrin, W. Daniel Stamer, Ayellet V. Segrè

## Abstract

**Purpose:** To identify the transcriptomic changes induced by dexamethasone (DEX) in trabecular meshwork (TM) and Schlemm’s canal endothelial (SCE) cells with RNA-sequencing (RNA-seq).

**Methods:** Human TM (n=10) and SCE cell strains (n=5) were isolated from healthy donor eyes and exposed to DEX 100nM and vehicle (control). Three DEX exposure times were evaluated: 1-hour, 6-hours, and 2 days. RNA-seq was performed on Illumina’s TruSeq platform and gene expression was quantified using featureCount. DESeq2 paired (treated and untreated) sample test was applied to identify genes transcriptionally responsive to DEX (DEGs) at false discovery rate <0.05. Gene-set enrichment analyses were performed on DEGs. DEGs were tested for association with glaucoma (POAG) and intraocular pressure (IOP).

**Results:** Nine TM and 4 SCE strains passed quality control. After 2-day DEX exposure, there were 857 and 2,086 DEGs in TM and SCE, respectively. Of these, 411 genes were differentially expressed in both TM and SCE, including *FKBP5* (17.3-fold-change, p=6.9×10^-53^) and *FAM107A* (25.1-fold-change, p=4.0×10^-240^), the most significant DEG after 2-day DEX exposure in TM and SCE, respectively. The 2-day DEX DEGs in TM and SCE were enriched in cell adhesion, extracellular matrix, and response to stimulus in Gene Ontologies (p<3.7×10^-6^). Early response DEGs were enriched in immune-related processes. Thirteen DEGs in TM were significant at all three time points, including *PER1*. *LTBP2* is a TM-only DEG and *FAM105A* a SCE-only DEG associated with IOP and POAG risk.

**Conclusions:** This study identified candidate genes and pathways for glucocorticoid-induced ocular hypertension which can be further explored in human genetic analyses.

## Background

Glucocorticoids (GC) are frequently prescribed for a variety of reasons including ophthalmic indications.^1^ However, GC have been shown to increase intraocular pressure (IOP). Intravitreal GC injections elevate IOP by greater than 10 mmHg in about a third of patients.^2^ This GC-induced ocular hypertension (OHTN) can result in GC-induced glaucoma, a form of open-angle glaucoma.^3,4^

Several lines of evidence suggest that GC responses are heritable.^5,6^ First-degree relatives of high GC responders have a greater likelihood of being high GC responders themselves,^7–9^ and twin studies have identified a heritability index of 0.35, a value comparable to more commonly known inherited traits such as blood pressure and body mass index.^10,11^

Aqueous fluid flows out of the eye through the conventional outflow pathway that includes the trabecular meshwork (TM)^12^ and Schlemm’s canal endothelial (SCE)^13^ cells and increased resistance to aqueous outflow results in elevated IOP. To establish the specific mechanisms of GC-induced IOP elevation, several studies have examined changes in protein expression in TM cells after GC exposure. In one study, dexamethasone (DEX) treatment of TM cells alters the abundance of about 40 proteins, suggesting that extracellular matrix remodeling and mitochondrial dysfunction are disease mechanisms.^14^ A correlation between differential expression of GC receptor isoforms and GC responsiveness in TM cells has also been described.^15^ Moreover, GC-induced changes in expression of transendothelial junction-associated proteins in SCE cells alters their permeability.^16^ RNA-sequencing (RNA-seq) of TM cells from human-organ cultures anterior segment examined differentially expressed genes (DEG) after DEX exposure for 7 days; pathway analysis revealed that Wnt signaling, cell adhesion, transforming growth factor beta (TGFβ), and mitogen-activated protein kinase (MAPK) signaling were associated with GC responsiveness.^17,18^ More recently, an RNA-seq study of DEX-treated transformed human TM cells found significant enrichment and potential interplay between the TGFβ signaling and cellular senescence pathways.^19^

Despite these multiple studies, our understanding of the transcriptomic changes induced by GC in TM and SCE cells remains limited. Previous proteomic studies have been limited to TM cells, and few have used RNA-seq technology, which has the potential to be more comprehensive and more sensitive in its ability to detect changes in gene expression in response to GC exposure. In addition, many of the transcriptomic experiments have been done with immortalized cell lines, which may not reflect physiology or biological diversity as accurately as primary cell strains. Specifically, transformed cells often cause chromosomal aberrations and transcriptomic changes compared to primary cells.

The purpose of this study is to identify the transcriptomic changes induced in TM and SCE cells after exposure to GC for different exposure times using RNA-seq, and to investigate the biological pathways they cluster in and their potential association with IOP and glaucoma.

## Methods

The study adhered to the Tenets of the Declaration of Helsinki and was exempted by the institutional review board of Duke University (Pro00113746).

### Cell Culture and DEX treatment

Primary human TM and SCE cell strains were isolated from donor eyes without any known diseases and cultured as described in Supplementary Material. The TM and SCE culture passages were limited to under 5 passages to maintain the primary nature of our cell strains. Cells were authenticated upon isolation using established methods^20–22^ and additional details are provided in the Supplementary Material. The TM and SCE cell strains were treated using two different conditions: (1) Ethanol-treated (vehicle), which served as the untreated control cells (labeled as CTRL), and (2) DEX 100 nM-treated cells which served as the experimental group (labeled as DEX), generating pairs of treated and untreated samples from each donor. The treated cells were exposed to 100 nM DEX daily for 1 hour, 6 hours and 2 days prior to RNA isolation. The primary time point of interest was 2 days as the phenomenon of GC-induced ocular hypertension usually takes days to months to manifest in patients. Therefore, ten TM cell strains and five SCE cell strains were used for this time point. The 1-hour and 6-hour time points were secondary time points chosen to capture the immediate and short-term effects of GC exposure on TM and SCE cells. Three TM and three SCE cell strains were used for the 1-hour time point, and four TM and four SCE cell strains were used for the 6-hour time point. Cell pellets from the samples were then sent to the Broad Institute for RNA sequencing.^23^

### RNA-Sequencing

RNA extraction and RNA-sequencing was performed at the Broad Institute following their RNA-Seq 2.0 protocol and details are provided in the Supplementary Material. RNA-seq reads were aligned to the reference human genome (GRCh37) using the STAR^24^ v2.4.1a aligner in two-pass mode with the GENCODE (v19) gene model. QC of all RNA-seq samples from the three time points are described in Supplementary Material.

### Gene expression quantification and differential gene expression

Gene expression quantification was computed using featureCount from the SubRead package.^25^ Details of the settings are provided in the Supplementary Material. Differentially expressed genes in response to DEX treatment were detected using a paired-sample test in DESeq2^26^, which gains power by estimating differential gene expression between treated and untreated samples from the same donor. The Wald test was used for significance testing, and a false discovery rate (FDR) < 0.05 was used to correct for multiple hypothesis testing and identify significant differentially expressed genes (DEGs). To account for the impact of known (e.g., batch effects, sequencing depth) and hidden confounders on gene expression, we applied SmartSVA^27^ to infer the surrogate variables (SVs) needed to correct for confounding effects in each of the cell line and time point datasets separately. The optimal total number of SVs accounting for the majority of non-specific variance explained per dataset was added to the DESeq2 design model for the differential gene expression analysis.

### Cell type identity confirmation

To confirm the endogenous identity of TM and SCE cells strains used, an inclusive list of cell type marker genes taken from the van Zyl *et al.* 2020 single cell RNA-seq atlas^23^ were extracted. For TM cell strains, various marker genes for beam cells (Beam A, Beam B) and JCT cell types were used. For SCE cell strains, gene markers specific to or highly expressed in SC endothelium were used. Overall, 17 gene markers for TM and 13 gene markers for SCE were examined for expression in transcripts per million (TPM) units across all control sample replicates. We also examined these genes’ differential expression patterns in response to DEX.

### qPCR validation of RNA-seq

To validate the RNA-seq results, we performed quantitative qPCR on a subset of the significantly differentially expressed genes. Details of the qPCR methods are in the Supplementary Material.

### Gene set enrichment analysis (GSEA) of DEX-responsive genes

GSEA was applied to all sets of significant DEGs (FDR < 0.05) following different DEX exposure times in TM and SCE using g:Profiler^28^ (ve102_eg49_p15_e7ff1c9) and a custom background gene list that included all genes expressed in TM and SCE, respectively. g:Profiler was applied to the following gene set resources: Gene ontology (GO), Kyoto Encyclopedia of Genes and Genomes (KEGG), Reactome, human phenotype ontology (HPO), and transcription factor target genes identified based on the TRANSFAC transcription factor binding site (TFBS) database. Metascape^29^ [v3.5] was used for GSEA of Hallmark gene sets^30^ GO-Figure^31^ (v1.0.1) was additionally used to identify a non-redundant set of significantly enriched GO terms that was obtained from the initial GSEA analysis. Only gene sets with 10 to 2000 genes were considered. For each enrichment analysis, the Benjamini-Hochberg adjusted P-value (adjP) were recorded along with log2(fold-change). Gene names were added to each GSEA output using BioMart,^32^ except for the Hallmark gene set enrichment output that directly provided gene names. Gene sets with adjP<0.05 were considered significant.

### Mapping of DEX-responsive genes to IOP and glaucoma GWAS loci

The identified high confidence DEGs were further examined for association with primary open angle glaucoma (POAG) and IOP genetic loci taken from the largest available GWAS meta-analyses of these traits, including the UK Biobank GWAS. Details of the analysis methods can be found in the Supplementary Material.

### Statistical Analyses

Unless stated otherwise the statistical analyses were performed using R,^33^ and ggplot2^34^ was used to generate figures in the paper. We corrected for multiple hypothesis testing using an FDR below 0.05.

## Results

### Sample characteristics and study design

Demographic and experimental characteristics of all the donors of the TM and SCE samples are summarized in **Table 1**. Cell type identities were authenticated upon isolation using established criteria^20–22^ and confirmed in our passaged cells using previously reported TM and SCE gene markers taken from a single-cell RNA-seq study^23^ (Methods and Supplementary Material). The strongest and most specific markers for the three primary TM cell types - trabecular beam A and B cells and juxtacanalicular tissue (JCT) cells - were highly expressed in all our TM samples (e.g., *MGP, MYOC, ANGPTL7, CMEMIP, PDPN, RARRES*) with an average TPM of 827, which included JCT (*CHI3L1*), Beam A (*FABP4*) and Beam B (*PPPR1B1*) specific markers (Supplementary Table 82), supporting the presence of all three TM cell types in our TM cultures isolated from the entire intact TM. Also, the strongest cell type-specific markers for SCE (e.g., *FN1, PLAT, POSTN*) were highly expressed in all our SCE samples, with an average TPM of 1374 (Supplementary Table 83). While the most highly expressed and specific TM and SCE markers were found in our cell strains, 4 of the 17 markers we inspected in TM (primarily JCT markers) and 6 of 13 markers inspected in SCE (expressed also in collector channel cells that were not isolated in our SCE samples) were either not expressed or expressed at low levels in a fraction of our samples (see Discussion for potential explanations). In both cell strains only a few cell type markers responded to DEX treatment, including upregulation of *MYOC* in TM (Supplementary Table 82) and downregulation of *POSTN* and *PLAT* in SCE (Supplementary Table 83) at 2 days. Once confluent, all cell strains underwent a differentiation step for one week before DEX (case) or no treatment (control) for 1 hour, 6 hours, and 2 days in TM and 1 hour, 6 hours, and 2 days in SCE following by RNA isolation (Methods and Supplementary Material). Differential gene expression and pathway enrichment analyses were applied to all time points and cell strains and comparisons were performed.

**Table 1.** Demographic data of donors of eyes from which TM and SCE cells were isolated.

| **Sample ID** | **Age at Time of Death** | **Sex** | **Ancestry** | **Glucocorticoid Exposure Time Point Studied** |
| --- | --- | --- | --- | --- |
| **Trabecular Meshwork (TM) Cell Strain Donors** |  |  |  |  |
| TM92 | 38 | Female | White | 2 days, 6 hours |
| TM96 | 28 | Male | White | 2 days, 6 hours, 1 hour |
| TM123 | 39 | Male | White | 2 days, 1 hour |
| TM134 | 51 | Male | White | 2 days |
| TM135 | 77 | Female | White | 2 days, 6 hours |
| TM137 | 60 | Male | White | 2 days |
| TM140 | 60 | Male | Black | 2 days |
| TM151 | 40 | Female | Asian | 2 days, 1 hour |
| TM155 | 58 | Female | White | 2 days, 6 hours |
| **Schlemm's Canal Endothelial (SCE) Cell Strain Donors** |  |  |  |  |
| SC67 | 44 | Male | White | 2 days, 6 hours |
| SC77 | 23 | Female | Black | 6hrs |
| SC86 | 62 | Male | White | 2 days, 1 hour |
| SC89 | 68 | Male | White | 2 days, 6 hours, 1 hour |
| SC91 | 74 | Female | White | 2 days, 6 hours, 1 hour |

### Two-day GC Exposure Results

Nine TM and four SCE paired samples from 2-day GC exposure passed QC (**Supplementary Material, Supplementary Tables 1-2, Supplementary Figures 1-2**). DEX-treated and control samples from each donor clustered together based on normalized gene expression profiles for both the TM and SCE samples (**Supplementary Figure 2),** suggesting, as may be expected, that inter-individual expression differences are larger than intra-individual DEX-induced transcriptomic differences.

### 2-day DEX-induced Differential Gene Expression in TM and SCE cells

To identify genes whose expression levels significantly changed in response to 2-day DEX exposure in TM and SCE cell strains, we adjusted for 4 and 2 surrogate variables (SVs), respectively (Methods). A total of 857 genes were differentially expressed with high confidence (FDR<0.05) in response to 2-day DEX in TM cells with 453 (52.85%) genes up-regulated and 404 (47.25%) down-regulated (**Supplementary Table 3**). Full DEG results for all genes tested in TM are shown in **Supplementary Table 4**. The genes that were most strongly affected in TM are shown in **Figure 1A**. **Table 2** and **Supplementary Table 5** show the results for the top twenty statistically significant DEGs in TM. In SCE, about 2.4-fold more genes (2,086 genes) were DEX-induced with high confidence (FDR<0.05) after 2-day treatment, with 1,066 (51.10%) genes up-regulated and 1,020 (48.89%) down-regulated (**Supplementary Table 6**). Full DEG results for all genes tested in SCE cells are shown in **Supplementary Table 7**. The genes that were most strongly affected in SCE are highlighted in **Figure 1B**. **Table 3** and **Supplementary Table 8** show the results for the twenty most significant DEGs in SCE.

**Figure 1.**
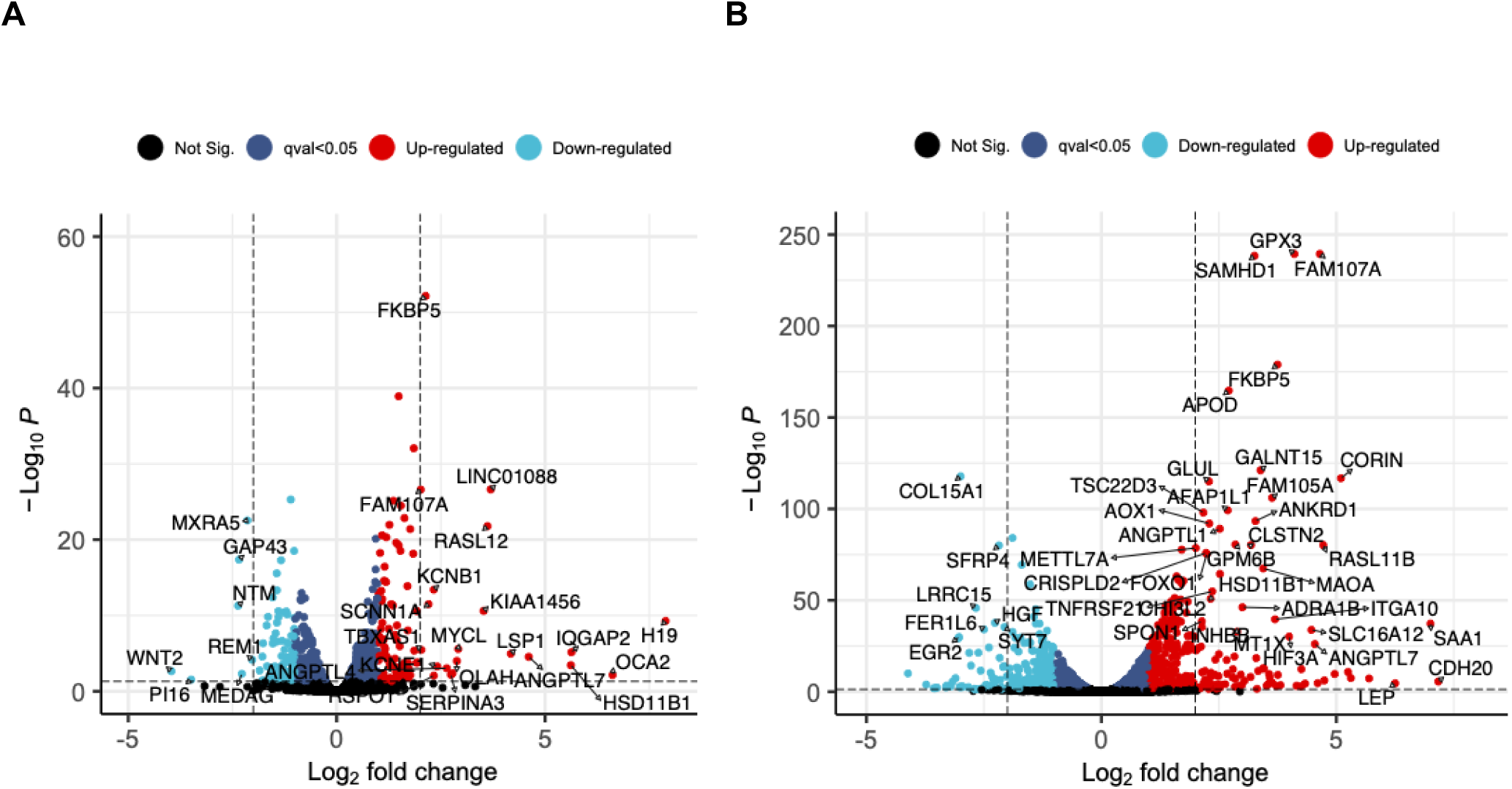
Differentially expressed genes (DEGs) after two days of dexamethasone exposure in trabecular meshwork (TM) and Schlemm’s canal endothelial (SCE) cells. Volcano plots display log_2_(fold-change) (LFC) on the *x*-axis and -log_10_(adjusted *P*-value) on the *y*-axis for (A) TM cells and (B) SCE cells. Red and blue points refer to significantly up- and down-regulated genes, respectively, at q-value <0.05. Highlighted (named) genes have >2 LFC and q-value <0.05.

**Table 2.**
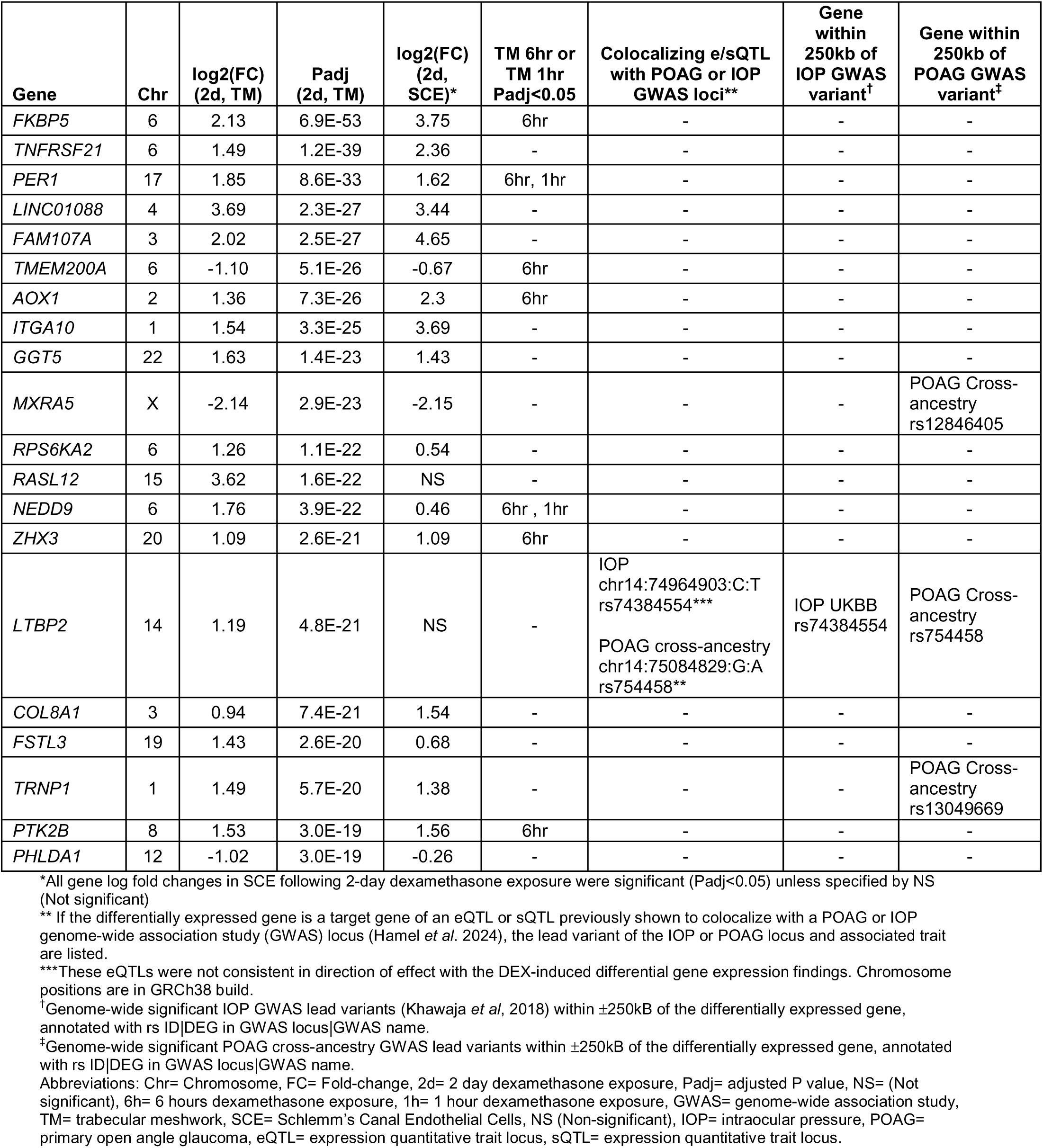
Twenty most statistically significant DEGs in TM cell strains following 2-day exposure to dexamethasone.

**Table 3.**
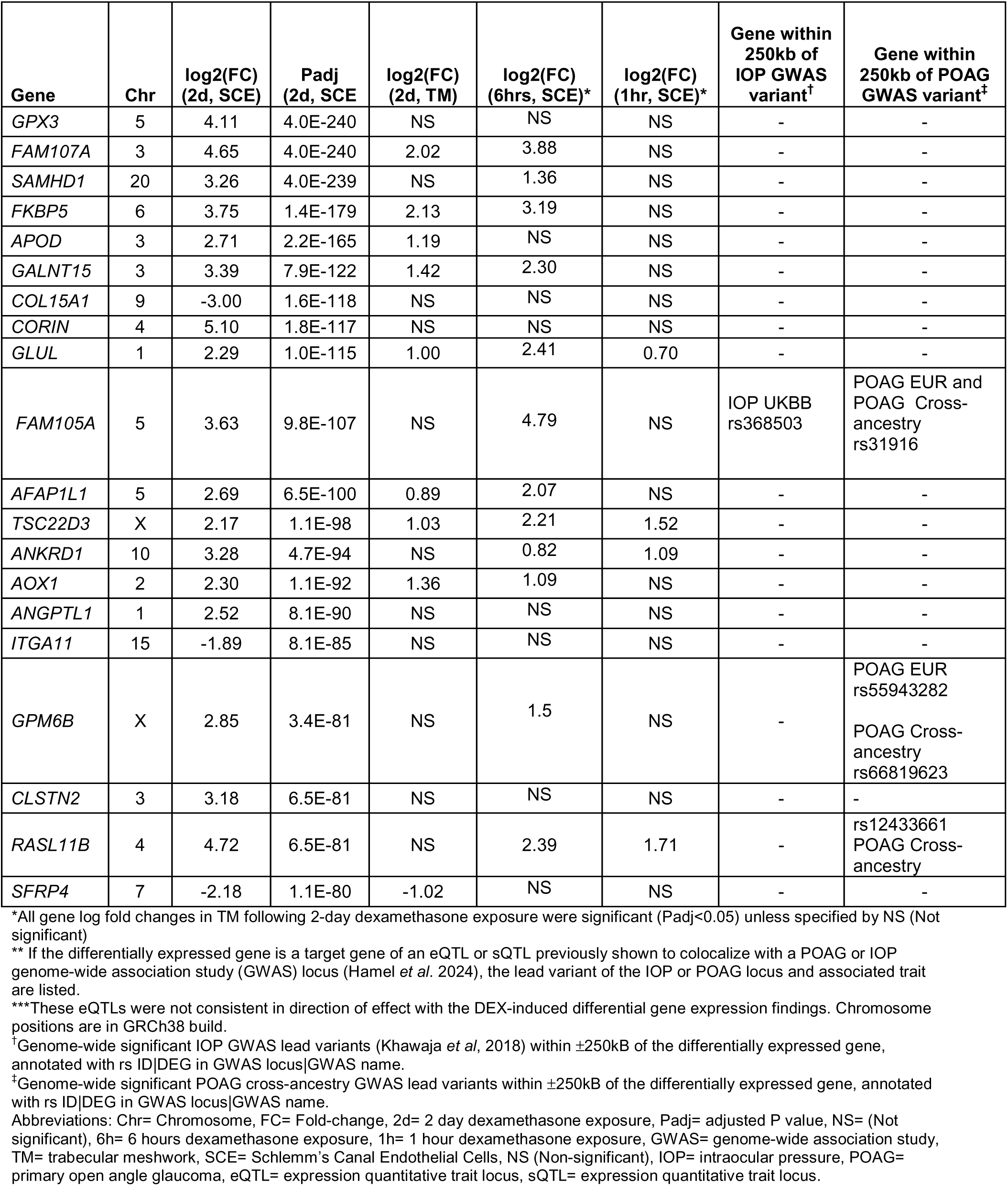
Twenty most statistically significant DEGs in Schlemm’s Canal Endothelium (SCE) cell strains following 2-day exposure to dexamethasone.

### Overlap between DEX-responsive genes in TM and SCE cells

A total of 411 genes (47% of TM DEGs and 20% of SCE DEGs) were differentially expressed in both TM and SCE cells after 2-day exposure to DEX (**Figure 2A, Supplementary Figure 3 and Supplementary Table 9**). The fold-change of the common DEGs were highly correlated between TM and SCE cells [Pearson correlation coefficient of log_2_(fold-change) r=0.62], with most (89.5%) of the DEGs showing the same direction of response to DEX exposure (**Figure 2B**). Of these, 187 DEGs were up-regulated and 181 were down-regulated in TM and SCE. Among the 43 genes whose fold-change were anti-correlated, 19 genes were upregulated in TM and downregulated in SCE, and 24 genes were downregulated in TM and upregulated in SCE. The fold-changes of the common DEGs were not statistically significantly different between TM and SCE (Wilcoxon rank sum test P=0.32).

**Figure 2.**
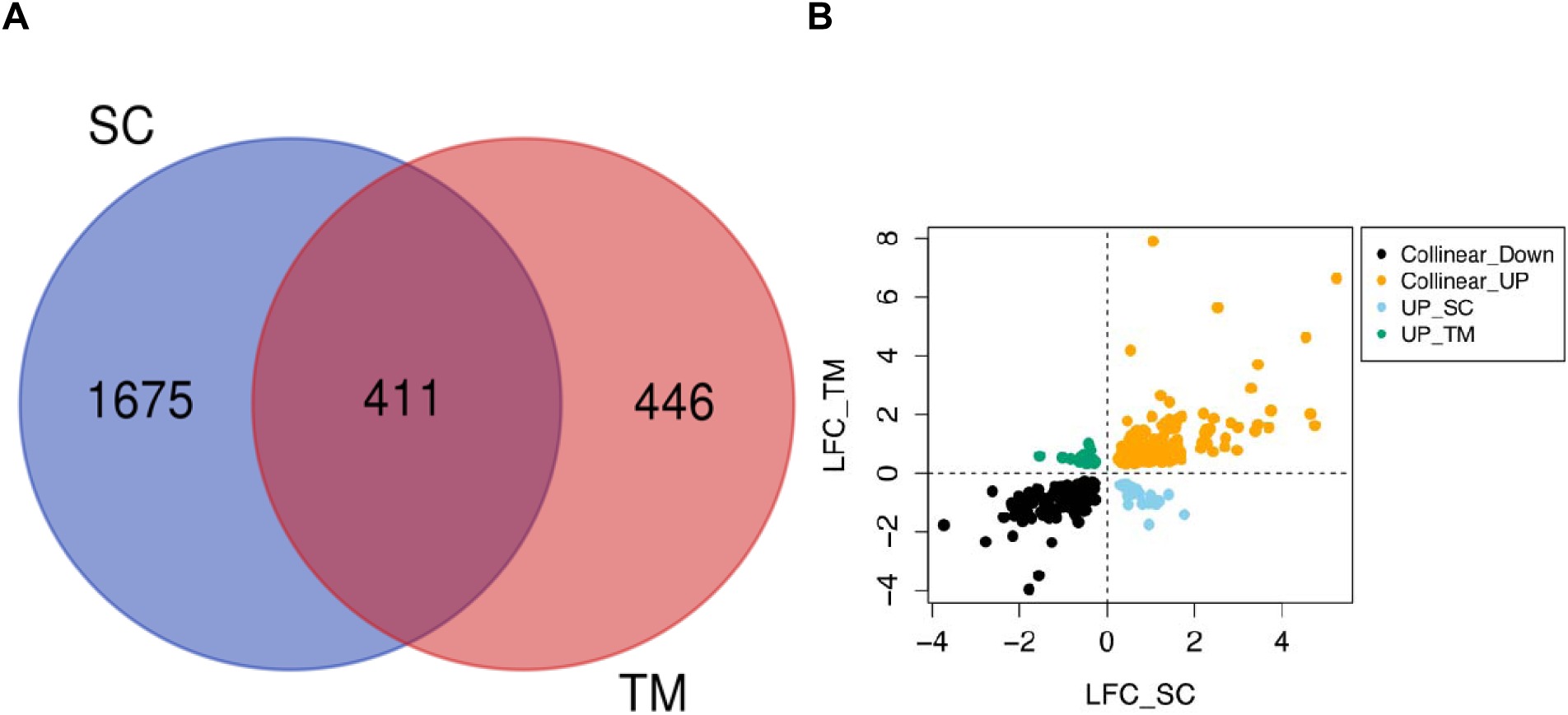
Comparison of differentially expressed genes (DEGs) between TM and SCE cells following 2-day dexamethasone exposure. (A) Venn diagram showing the overlap of DEGs between TM and SCE. (B) Comparison of the directionality of gene expression changes of the 411 common DEGs between TM and SCE cells following 2-day exposure to dexamethasone. Collinear Down: DEGs are positively correlated and down-regulated in both TM and SCE; Collinear_UP: DEGs are positively correlated and up-regulated in both cell types; UP_SC: DEGs are negatively correlated, up-regulated in SCE and down regulated in TM; UP_TM: DEGS are negatively correlated, up-regulated in TM and down regulated in SCE. TM: trabecular meshwork; SCE: Schlemm’s canal endothelial.

### Gene set enrichment analysis of 2-day DEX-responsive DEGs in TM and SCE

To gain biological insight into the transcriptomic effects of 2-day DEX exposure in TM and SCE, we tested whether the DEGs were enriched in specific biological pathways or gene ontologies by applying GSEA to different gene set databases. The results are summarized in the Supplementary Material, **Supplementary Figure 4** and **Supplementary Tables 10 - 21** for TM cells, and **Supplementary Figure 5** and **Supplementary Tables 22 - 33** for SCE cells. DEGs in TM were enriched in multiple pathways, including extracellular matrix organization, the top finding from Reactome (adjusted P-value [adjP]=1.59E-09) and Gene Ontology (GO) (BP, adjP=1.48E-11). DEGs in SCE cells were enriched most strongly in pathways for signal receptor binding and extracellular matrix-receptor interaction; the top findings from the GO (MF, adjP=5.75E-15), Reactome (adjP=6.37E - 07) and KEGG (adjP=9.24 E-05) databases **(Figure 3)**. From the ECM-enriched pathways identified from the 2-day time point data for TM cells, a total of 56 genes with 20 (35%) up-regulated and 36 (64%) down-regulated were significantly (q<0.05) differentially expressed (**Supplementary Figure 6A**). Similarly, for the SCE cells, 100 genes with 39 (39%) up-regulated and 61 (61%) down-regulated were significantly differentially expressed (**Supplementary Figure 6B**).

**Figure 3.**
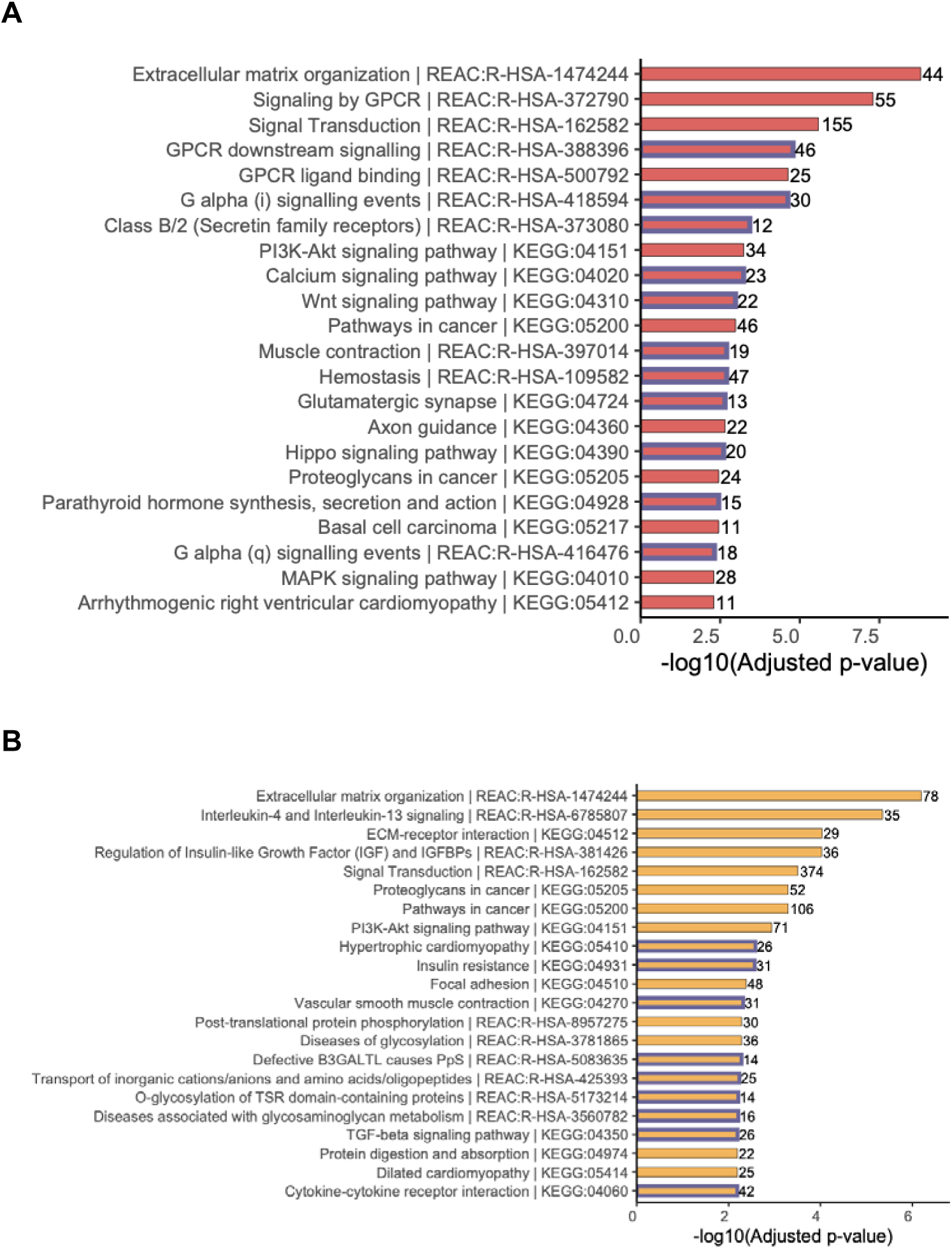
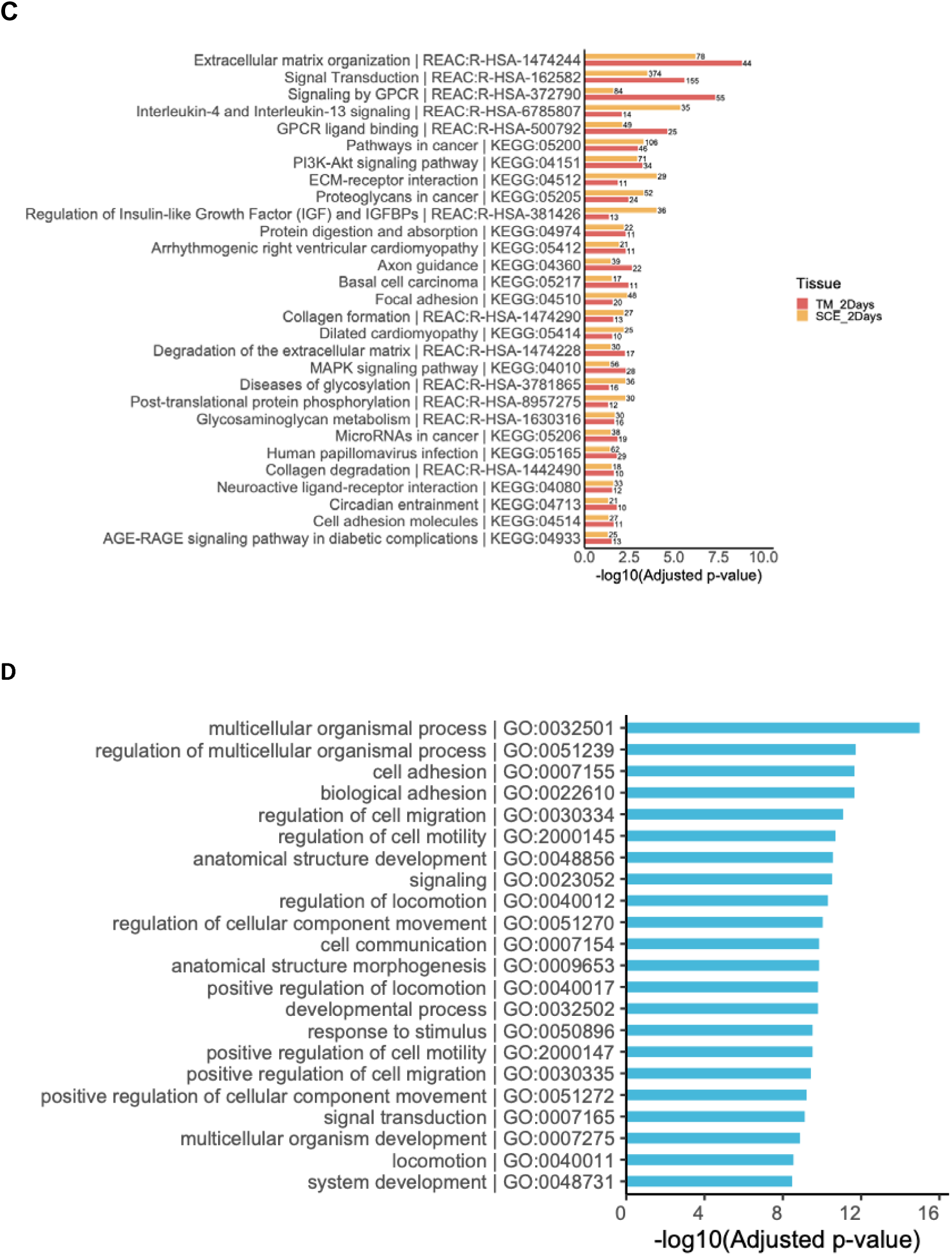
Top gene sets enriched for differentially expressed genes following 2-day dexamethasone (DEX) exposure in TM and SCE cells. Bar plots of top gene sets from Reactome and KEGG databases are shown for (A) TM 2-day DEX exposure, (B) SCE 2-day DEX exposure, (C) common gene sets between TM and SCE 2-day DEX exposure, and (D) from gene ontology (GO) biological processes for the common 411 DEGs between TM and SCE 2-day exposure samples. Blue outlines of the bars specify gene sets that are only significant at Padj<0.05 for the given cell type.

We further applied GSEA to the common TM/SCE set of 411 DEGs. Enrichment for biological pathways identified extracellular matrix organization (adjP=0.02), and Interleukin-4 and Interleukin-13 signaling (adjP=0.02) from Reactome databases (**Supplementary Table 34**). Additional results for other gene sets are shown in **Supplementary Tables 35-38**. Results for non-redundant GO terms are displayed in **Supplementary Figure 7**. From the non-redundant enriched GO terms, regulation of multicellular organismal process (GO BP adjP=1.90E-12), cell adhesion (BP adjP=2.2E-12), regulation of cell migration (BP adjP=8.3E-08), response to stimulus (BP adjP=3.0E-10), extracellular matrix (CC, adjP=1.01E-16), extracellular matrix structural constituent (MF, adjP=2.72E-05) and receptor ligand activity (MF, adjP=2.718E-05) (**Figure 3**) were identified. Extracellular matrix was a common pathway observed across many GSEA.

### Mapping DEX-responsive genes to POAG and IOP GWAS loci

To test whether any of the DEX-responsive genes may be associated with IOP or POAG risk, we first examined whether any of the DEGs fell within 250 kb of over a hundred known genome-wide significant variants associated with IOP and/or POAG based on large cross-ancestry and European GWAS meta-analyses^35–37^ and Multi-trait Analysis of GWAS (MTAG)^36^ (see Methods). 83 (9.7%) of the DEGs in TM and 207 (9.9%) of the DEGs in SCE fell within 250 kb of IOP and/or POAG GWAS variants, respectively (**Supplementary Tables 3** and **6**). Among the 20 most significant DEGs in TM, three genes fell within 250 kb of a POAG variant: *MXRA5* (near rs12846405, POAG variant),^35^ *LTBP2* (near rs754458, IOP, POAG and MTAG variant)^35,36^, and *TRNP1* (near 13049669, POAG variant)^35^ (**Table 2**). Among the 20 most significantly DEGs in SCE cells, three other genes also fell within 250 kb of a POAG or IOP GWAS variant (none were found near MTAG variants)^36^: *FAM105A* (near rs368503, IOP variant, rs31916 POAG variant),^37^ *GPM6B* (near rs55943282, POAG variant),^38^ and *RASL11B* (near rs12433661, POAG variant)^35^ (**Table 3**).

### Differential Gene Expression Across All Three Time Points

The results for the 6-hour and 1-hour experiments are described separately in the Supplementary Material. We also examined DEGs which were identified across all three time points.

### TM Cells

When examining the DEGs across the three time points (1 hour, 6 hours and 2 days) of DEX exposure in TM cells, 741 (73.65%) DEGs were unique to 2 days, 134 DEGs (13.20%) were unique to 6 hours, and 12 DEGs (1.19%) were unique to 1 hour (**Figure 4**). Thirteen DEGs (1.29%) were common to all three time points, 98 DEGs (9.74%) were common to 6-hour and 2-day experiments, 5 DEGs (0.49%) were common to 1-hour and 2-day experiments, and 3 DEGs (0.29%) were common to the 1-hour and 6-hour experiments. Thirteen common genes were identified showing differences in expression patterns (Kruskal-Wallis chi-squared p-value = 0.018; Wilcoxon rank sum exact test p-value = 0.04). For the 13 DEGs common to all three time points, the LFC expression over time is shown in **Figure 5**. *PDK4* shows the widest variation in expression over the time points*. PER1, NEDD9, TSC22D3* and *LIF* show intermediate levels of variation and cluster together. The remaining genes show less variation in expression over time. For most genes, the highest expression was seen at the 6-hour time point.

**Figure 4.**
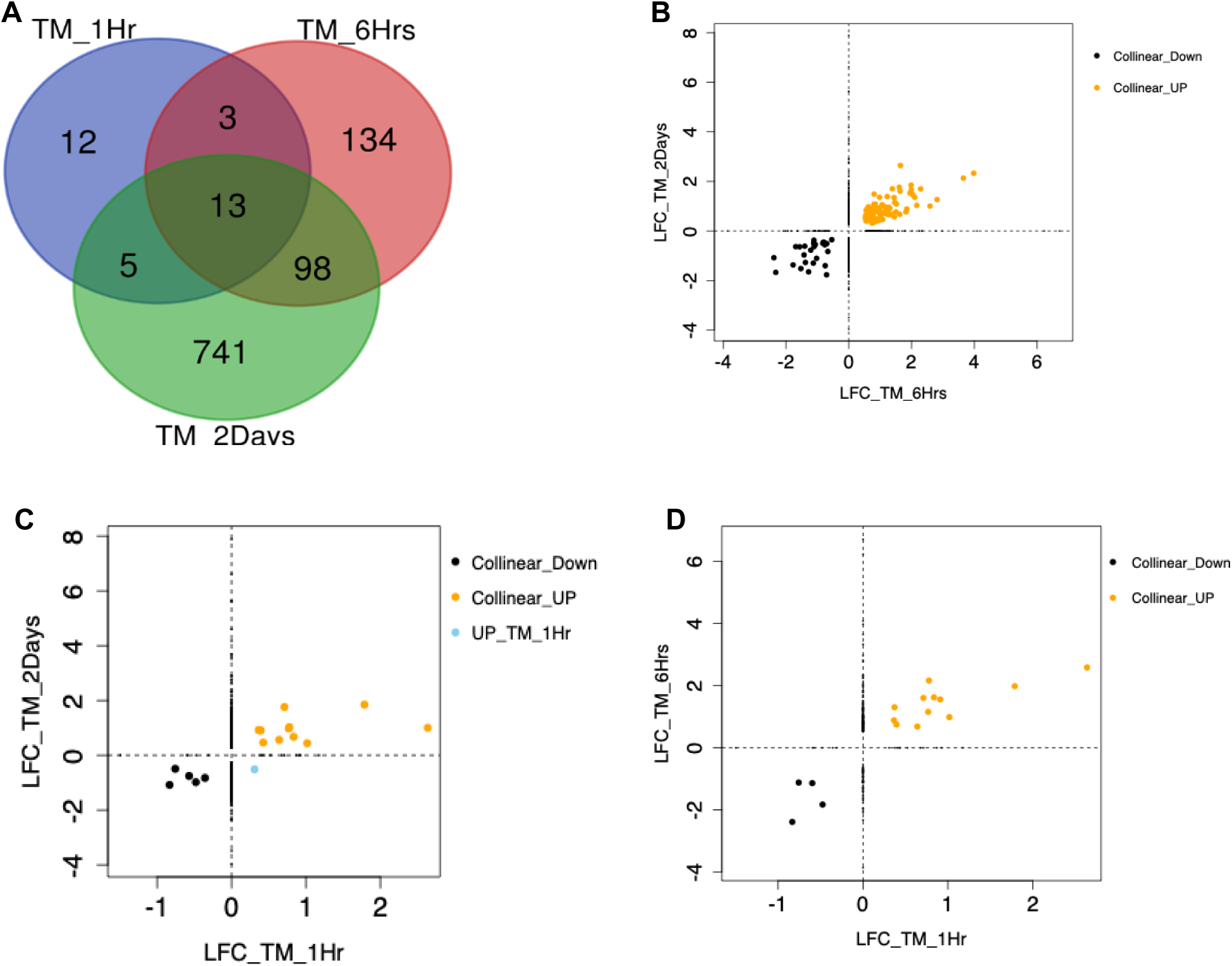
Comparison of DEGs in TM from 1-hour, 6-hours and 2-days experiments of dexamethasone (DEX) exposure. (A) Venn Diagram showing common DEGs between the different DEX exposure time points. (B) Scatter plot of log2(fold-change) (LFC) of the significant differentially expressed genes (DEGs) (adjP<0.05) for TM 2-day versus 6-hour experiments. Collinear_Down=26 DEGs, Collinear_UP=85 DEGs. (C) Scatter plot of LFC of the significant DEGs (adjP<0.05) for TM 2-day versus 1-hour experiments: Collinear_Down=5 DEGs, Collinear_UP=12 DEGs, UP_TM_1Hr=1 DEG. (D) Scatter plot of DEG LFC for the TM 6-hour versus 1-hour experiments: Collinear_Down=4 DEGs, Collinear_UP=12 DEGs. Collinear_Down: positive correlation of LFC and DEGs down-regulated in both time points; Collinear_UP: positive correlation of LFC and DEGs up-regulated in both time points; UP_TM_1Hr: upregulated in TM 1-hour and downregulated in TM 2-days exposure.

**Figure 5.**
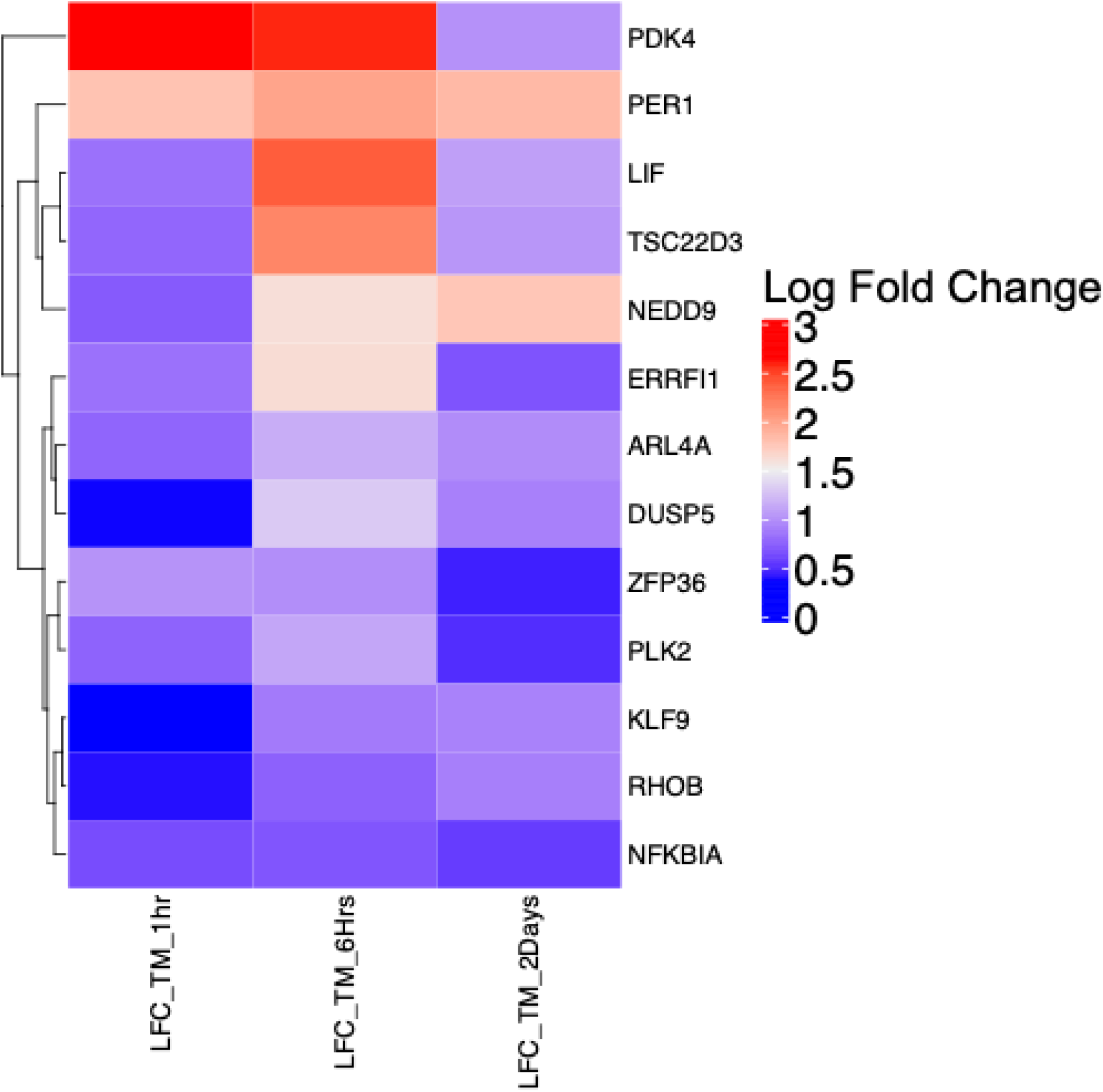
Heatmap clustering of differentially expressed genes across all three time points of dexamethasone (DEX) exposure in trabecular meshwork cells. The color bar represents gene expression fold-change in response to DEX treatment at the different time points.

### SCE Cells

When examining the DEGs across the three time points (1 hour, 6 hours and 2 days) of DEX exposure in SCE cells, 1,164 (26.53%) DEGs were unique to 2 days, 2,282 DEGs (52.02%) were unique to 6 hours, and 8 DEGs (0.18%) were unique to 1 hour (**Figure 6**). Thirty-two (32) DEGs (0.72%) were common to all three time points, 885 DEGs (20.1%) were common to 6-hour and 2-day experiments, 5 DEGs (0.1%) were common to 1-hour and 2-day experiments, and 10 DEGs (0.22%) were common to the 1-hour and 6-hour experiments. In total 32 genes were identified as common to all the three time points and 21 of these showed a similar upregulated expression pattern (Kruskal-Wallis chi-squared p-value =0.45). For the 32 DEGs common to all three time points, the LFC expression over time is shown in **Figure 7**. *RASL11B* shows highest expression at 2 days among all the common SCE genes with decreasing expression across the time points. Non-monotonic high expression changes were observed in other genes such as *ANKRD1*, *MT1X*, *TSC22D3*, *GLUL* and *MT1E*. Intermediate changes were observed in *ANGPTL4*, *ARL4A*, *DUSP5*, and *ERRFI1*. At the 1-hour time point, *KLF9*, *TXNIP* and *NNMT* showed least expression with rise in expression at 6 hours and 2 days. Two genes, *C10orf10* and *KCNE4* show high expression at 6 hours with low expression at 1 hour and 2 days. Interestingly, *DDIT4* showed high expression at the early 1- and 6-hour time points but then was downregulated at the 2-day time point. *TNFAIP3* also showed variation in change with moderate expression at 1 hour, downregulation at 6 hours and upregulation at 2 days. Nine genes including *LIF*, *IER3*, *HES1*, *GDF15*, *ID4*, *JUN*, *IRF1*, *IER2* and *BHLHE40* were downregulated across all three time points. *LIF* showed the largest downregulation at 6 hours in comparison to 1 hour and 2 days.

**Figure 6.**
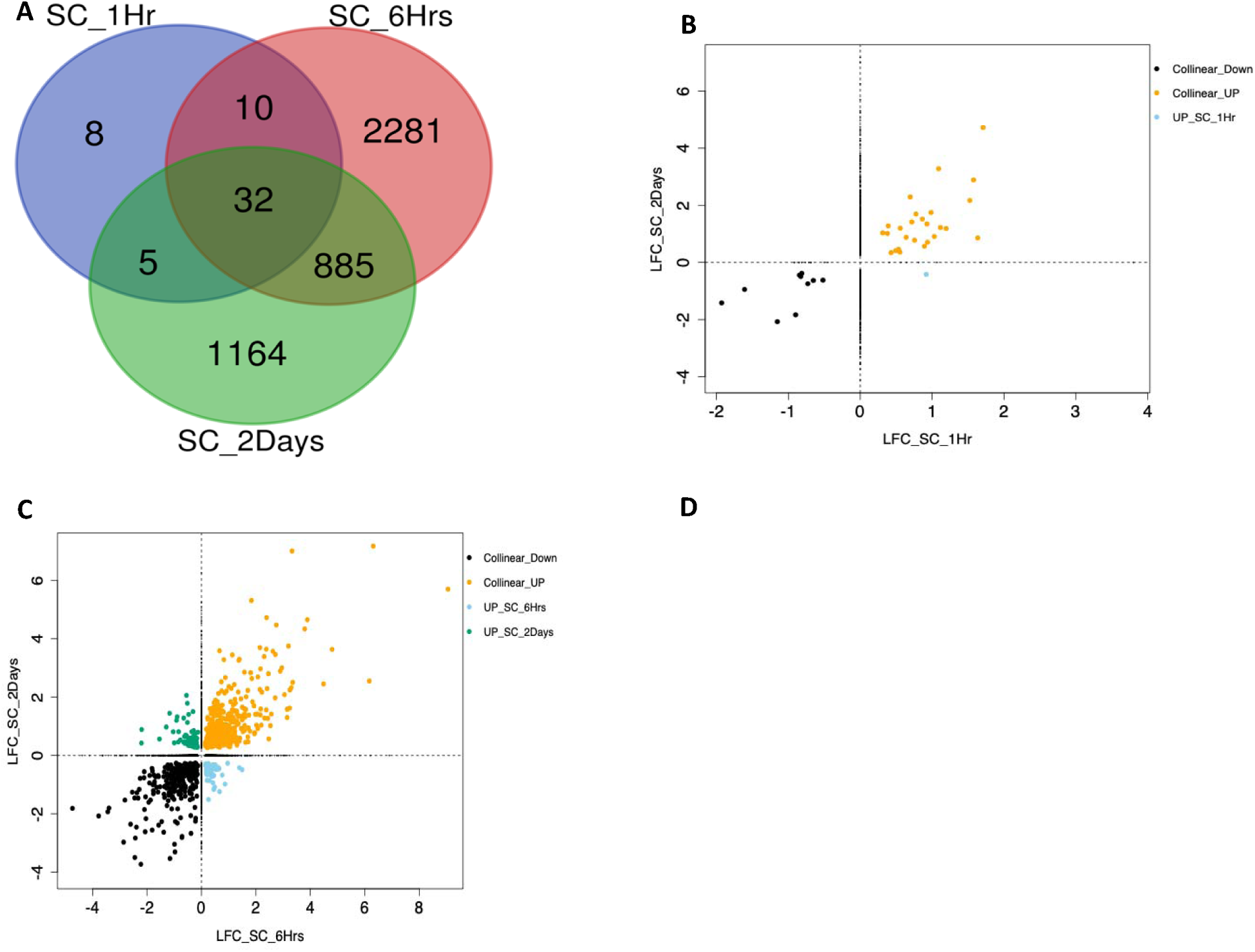
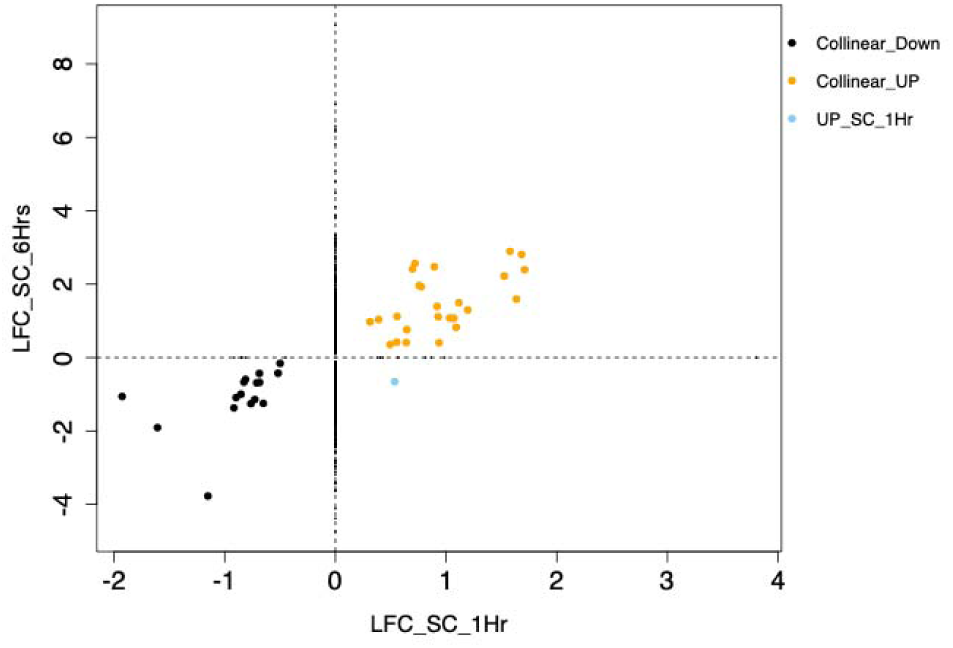
Comparison of differentially expressed genes (DEGs) in Schlemm’s Canal Endothelial (SCE) cells from 1-hour, 6-hours and 2-day experiments of dexamethasone (DEX) exposure. (A) Venn Diagram showing common DEGs between the different DEX exposure time points. (B) Scatter plot of LFC of the significant DEGs for SCE 2-day versus 1-hour experiments: Collinear_Down=10 DEGs, Collinear_UP=26 DEGs, UP_SC_1Hr=1 DEG. (C) Scatter plot of log2(fold-change) (LFC) of the significant DEGs (adjP<0.05) for SCE 2-day versus 6-hour experiments. Collinear_Down=341 DEGs, Collinear_UP=429 DEGs, UP_SC_2Days=67 and UP_SC_6Hrs=80. (D) Scatter plot of DEG LFC for the SC 6-hour versus 1-hour experiments: Collinear_Down=16 DEGs, Collinear_UP=25 DEGs, UP_SC_1Hr=1 DEG. Collinear_Down: positive correlation of LFC and DEGs down-regulated in both time points; Collinear_UP: positive correlation of LFC and DEGs up-regulated in both time points; UP_SC_1/6Hr: upregulated in SC 1/6-hour and down-regulated in comparative time point; UP_SC_2Days: upregulated in SC 2-days and down-regulated in comparative time point

**Figure 7.**
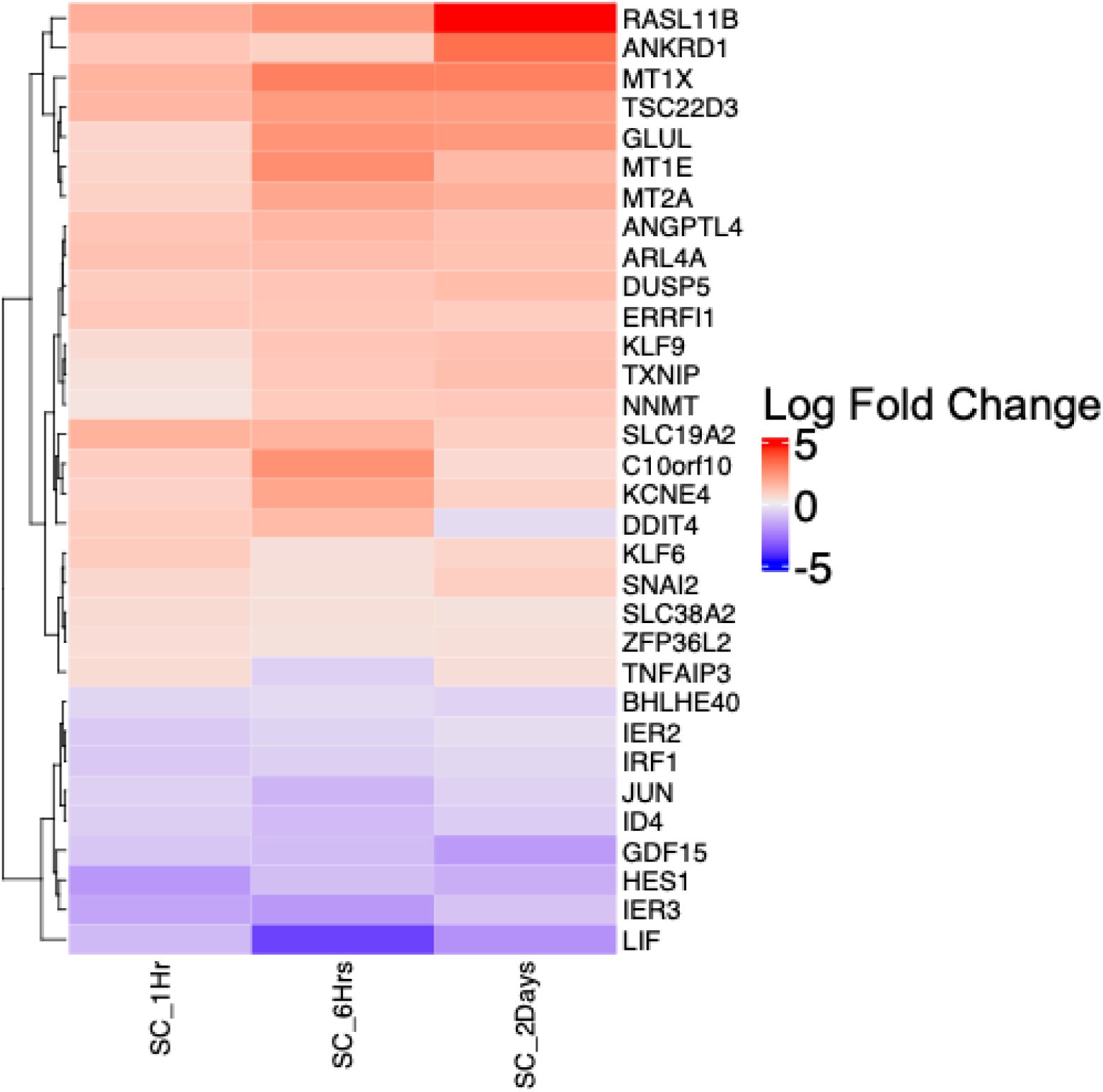
Heatmap clustering of differentially expressed genes across all three time points of dexamethasone (DEX) exposure in Schlemm’s canal endothelial (SCE) cells. The color bar represents gene expression fold-change in response to DEX treatment at the three different time points.

### Validation of differential gene expression with quantitative PCR

We performed quantitative polymerase chain reaction (qPCR) validation on 11 DEX responsive genes in 9 TM samples and 10 DEX responsive genes in 4 SCE samples from the 2 day exposure experiments. High correlation was found between the average qPCR relative quantification (RQ) and RNA-seq fold-change in both TM (Pearson correlation coefficient, r=0.94, P=1.6E-05; **Figure 8A**) and SCE (Pearson r=0.98, P<4.9E-07; **Figure 8C**) samples. When examining the correlation for each TM donor separately, <u>high correlation was observed</u> <u>for 8 of the 9 TM donors</u> (r=0.85-0.96, P<9.2E-04; **Figure 8B**), with primarily TM137 exhibiting higher noise levels (r=0.61, P=0.048; Figure **8B**). For SCE, <u>high correlation for each of the 4</u> <u>donors</u> was observed (r=0.96-0.99, P<1.4E-05; **Figure 8D**). See Supplementary Material for more details.

**Figure 8.**
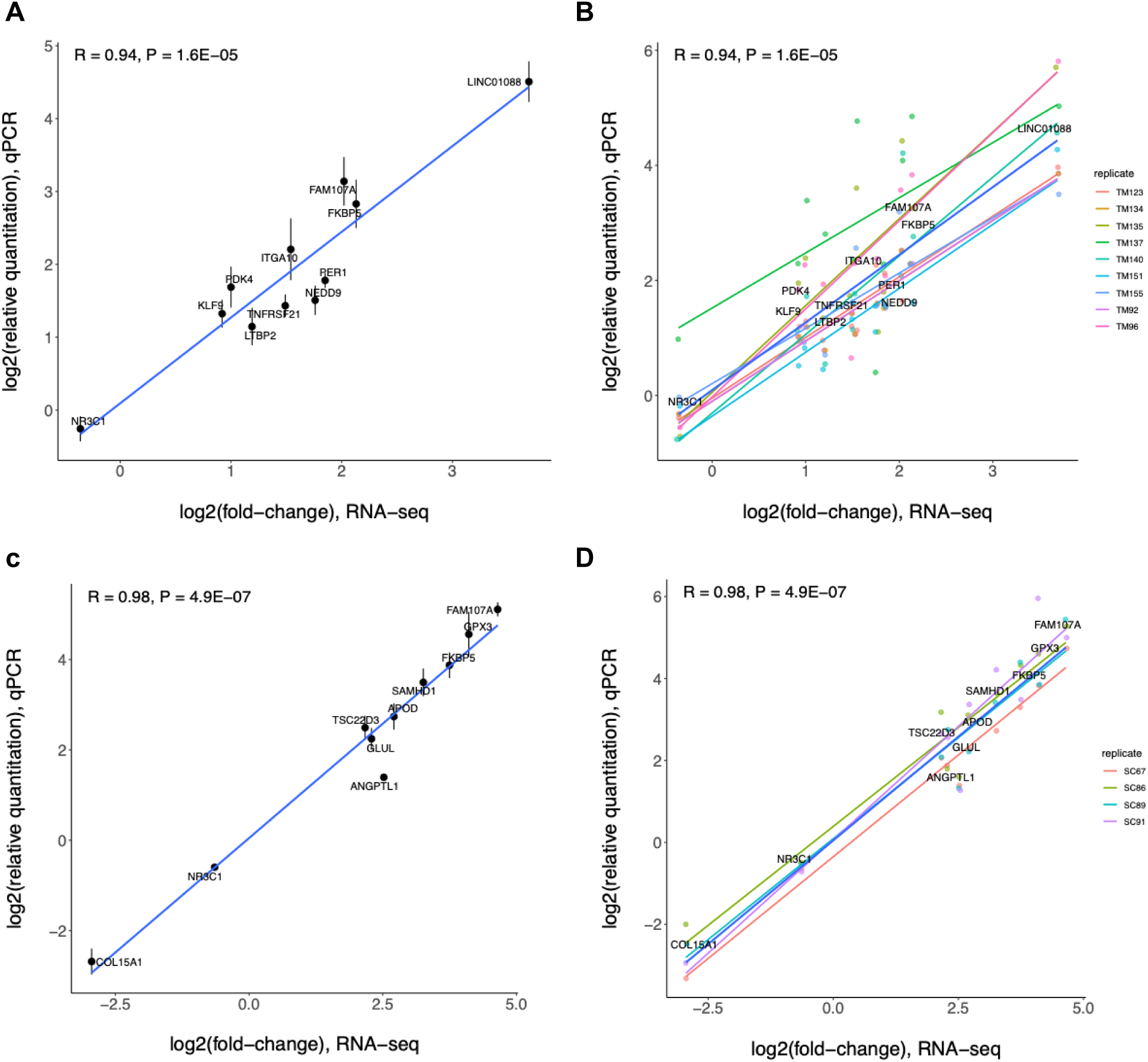
Correlation of qPCR relative quantification (RQ) values and RNA-seq fold-change in (A,B) trabecular meshwork (TM) and (C,D) Schlemm’s canal endothelial (SCE) cell strains in response to dexamethasone treatment for 2 days. Values are shown in log2 scale, and the error bars represent standard error (s.e.) of the average RQ values per gene across 11 TM donor pairs (A) and 4 SCE donor pairs (C). A linear line is fit using all donors per cell strain (A,C) or per donor (B,D). R, Pearson correlation coefficient.

## Discussion

This investigation represents the largest and most comprehensive analysis of transcriptomic changes in TM and SCE cells in response to GC exposure. Importantly, this study examines the changes over multiple time points of exposure, including immediate, short-term and long-term exposure. Because they recapitulate gene expression patterns in humans much better than immortalized human cell lines, experiments were carried out on human primary cell strains from outflow tissues dissected from non-diseased eyes. Our TM and SCE cell strains indeed highly express well established marker genes of JCT, Beam A and Beam B, and SC endothelial cells. We note that some of the previously reported marker genes (24-46%) were lowly expressed or absent from our TM or SCE strains. This may be due to differences in cell culture compared to endogenous tissue, heterogeneity of cell type abundance or expression heterogeneity between bulk tissue and single cell studies.

We have identified multiple genes with significant and consistent expression changes in response to GC in both TM and SCE. A minority of these genes are known to impact IOP and POAG risk from previous GWAS studies, but several are also novel genes that have not been previously identified as important GC-altered genes. *FKBP5*, the most significant DEG in TM cells (17.3-fold expression change) after 2-day GC exposure, is also a significant DEG in SCE cells after 2-day GC exposure. *PER1* was identified as a DEG across all three time points in TM cells, which also has a transcription binding site for NRC31. GSEA of the DEGs consistently finds extracellular matrix as a pathway that is affected by GC-exposure in TM and SCE cells, as previously reported.^39^ Upregulation of genes linked to extracellular matrix regulation has been observed in bulk RNA sequencing of TM cells.^39^ GSEA in this study has also identified other novel pathways, including immune-related pathways, such as interleukin-4 and interleukin-13 signaling. These insights from RNA-seq can be leveraged in human genetic studies of GC-induced ocular hypertension to identify novel genes involved in disease pathophysiology.

*FKBP5*, FK506-Binding Protein 5, encodes a protein that is a member of the immunophilin protein family, plays a role in immunoregulation and basic cellular processes involving protein folding and trafficking, and is thought to mediate calcineurin inhibition.^40^ It was the most significant DEG in TM cells after 2-day DEX exposure and the fourth top DEG in SCE cells after 2-day DEX exposure. *FKBP5* (also known as *FKBP51*) has been shown to be involved in the nuclear transport of the human glucocorticoid receptor in TM cells,^41^ however a small human study with 107 steroid responders and 400 controls failed to show any association between GC-induced OHTN and ten SNPs tested in *FKBP5*.^42^ Given the degree of differential expression in response to GC exposure in both TM and SCE cells, more comprehensive examination of genetic variation in this gene in a larger human genetic association study is warranted.

*PER1*, period circadian regulator 1, encodes components of the circadian rhythms of locomotor activity, metabolism and behavior. In this study, its expression was consistently increased across all three time points in TM cells. To our best knowledge, it has not been previously reported to be associated with GC-induced OHTN. *PER1* has been identified in the nonpigmented epithelium of the ciliary body, but not in TM, of mice.^43^ It has also been shown to have a circadian pattern of expression in the iris-ciliary body complex of mice and its expression has been correlated with diurnal variations in IOP.^44^

*FAM107A*, family with sequence similarity 107 member A, encodes a protein that enables actin binding activity. It was a significant DEG for both the SCE (rank #2, 21.5-fold expression change) and TM (rank #5) following 2-day GC exposure. *FAM107A* has been previously found to be upregulated in RNA-seq analyses in human organ-cultured anterior segment model of GC-response for 7 days.^17^ *TNFRSF21*, TNF receptor superfamily member 21, is another highly DEG in TM and SCE. It encodes a tumor necrosis factor receptor which activates nuclear factor kappa B and induces apoptosis.^45^ In a functional pathway cluster analysis using human microarray datasets that investigated the effect of DEX versus control medium on TM tissue, *TNFRSF21* was found in two (apoptosis, adipogenesis) of the nine functional pathway clusters identified by the authors to synthesize the molecular response to DEX exposure in TM cells.^46^ It has also been found to be downregulated in TM from patients with POAG compared to controls.^47^

*GPX3* was the most significant DEG in SCE cells after 2 days of GC exposure but was not a significant DEG in TM cells at any time point tested, nor at the 1-hour SCE time point. Interestingly, it was also identified in two (inflammation, oxidative stress) of the nine functional pathway clusters identified by the aforementioned study of the molecular response to DEX exposure in TM cells.^46^ *GPX3* encodes glutathione peroxidase which catalyzes the reduction of organic hydroperoxides by glutathione, thereby protecting cells against oxidative damage.

In comparing our results with previously published studies of gene expression changes in TM cells, there was some overlap observed. In both the current study and the study of RNA-seq of TM cells from human-organ cultures of the anterior segment with 7-day DEX exposure, the gene *APOD* (apolipoprotein D) had significantly increased expression; apolipoprotein D is an enzyme involved in lipoprotein metabolism.^17^ The TGFβ pathway was associated with GC responsiveness in two studies.^17,19^ We found that the TGFβ induced gene (*TGF*β*I*) was a high confidence DEG for both TM and SCE cells after 2-day DEX exposure.

In comparing our results with a previously published study of proteomic changes in TM cells exposed to DEX for 10 days,^14^ there were seven DEGs from the current study that overlapped with the 40 proteins significantly altered in abundance in that paper. Those genes are *ALCAM, FLNB, SERPINH1, PLS3, P4HA2, TAGLN, and NES*. In Table 2 of the paper the *ALCAM* gene coding for the CD166 antigen and *FLNB* coding for Filamin-B were elevated in DEX-treated TM cells and were involved in cytoskeletal/cell-cell/matrix interactions. Table 3 of the paper shows that the genes *SERPINH1, PLS3, P4HA2, TAGLN,* and *NES* coding for proteins Serpin H1, Plastin-3, Prolyl 4-hydroxylase subunit alpha-2, Transgelin, and Nestin, respectively, were all found to be decreased in DEX-treated TM cells. *SERPINH1* is involved in stress response, cellular defense, and protein processing. *PLS3, P4HA2, TAGLN*, and *NES* are indicated to be involved in cytoskeletal and extracellular matrix interactions. Additionally, Serpin H1, Plastin-3, and Transgelin are calcium-binding proteins.

There were some DEGs which have been previously identified to contain or lie near variants associated with IOP or POAG by GWAS. Their existence among the DEG findings in our study indicates that DEGs identified have clear potential to impact human disease risk. Variants in *LTBP2* have been associated with both IOP and POAG.^35,37^ *TRNP1* and *MXRA5* were DEGs for both TM and SCE at the 2-day exposure time point and both are near POAG associated variants in a large GWAS study.^35^ In addition, *MXRA5* was a DEG at the 6-hour time point in TM cells. *FAM105A* was a DEG in SCE at the 2-day time point only, and this gene is near POAG and IOP associated-variants.^35,37,38^ *GPM6B* and *RASL11B* were DEGs for the 2-day SCE time point, and *RASL11B* was also a DEG for the 1-hour SCE time point; both genes are in known POAG loci.^38,48^ Myocilin (*MYOC*) is a gene that has been implicated in early onset glaucoma and POAG,^35–37^ and it was differentially expressed in TM cells (P adjusted = 4.35 × 10^-6^) in our study in response to GC. Finally, *ANGPTL7* has been associated with IOP in GWAS^37^ was differentially expressed in both TM and SCE cells after 2-day DEX exposure (P adjusted = 2.87 × 10^-5^ and 8.54 × 10^-27^, respectively).

GSEA of the DEGs implicated several pathways in GC-induced ocular hypertension. The most consistent pathway identified across the different DEX-exposure time points and in both SCE and TM cells, with various gene set databases, was the extracellular matrix pathway, including degradation of the matrix and interaction of the matrix with receptors. This is consistent with what has been found by other pathway or GSEA of the GC response.^6,46^ Extracellular matrix organization in the TM is different in GC-induced-glaucoma patients in comparison to that in POAG patients.^49^ GC treatment initiates biophysical alterations associated with increased resistance to aqueous humor outflow and a resultant increase in IOP.^50^ DEX treatment increases TM cell stiffness. Concurrent activation of the mitogen-activated protein kinase (MAPK) pathway stiffens the extracellular matrix *in vitro* along with upregulation of Wnt antagonists and fibrotic markers embedded in a more organized matrix and increases the stiffness of TM tissues *in vivo.*^50^ *In vitro* extracellular matrix derived from DEX-treated cells is stiffer than extracellular matrix deposited by control TM cells and is associated with compositional changes, with fibronectin morphology being the most significantly altered, and elevated mRNA expression of *MYOC*.^51^ One difference noted between the GSEA results is that shear stress was a pathway enriched more strongly for in SCE, than TM; this is an interesting finding particularly given that human GWAS studies for glaucoma traits have shown shear stress as an important pathway.^52^ DEX-induced DEGs in SCE following 2-day treatment were enriched for vasculature development processes, but not DEGs in TM. Of not, genes associated with IOP in a large GWAS meta-analysis are enriched in vasculature development.^52^

We found many more DEGs in SCE versus TM. The reason for this difference is unclear, especially given that the sample size was larger for TM but may be due to a potentially higher transcriptional activity in the SCE due to its unique blood and lymphatic vascular characteristics.^53^ In this way, SCE may be more sensitive to GC due to its role as part of the blood-aqueous barrier.

The current study expands our knowledge of the transcriptomic changes in the anterior segment of the eye in response to GC. Using both TM and SCE we document the change in gene expression over three time points and have validated a subset of these results with qPCR, including the downregulation of *NR3C1* that encodes the glucocorticoid receptor, and upregulation of genes previously shown to response to steroid, such as *KLF9*, *PER1* and *FAM107A*. One area for future exploration is to evaluate the extent to which the GC-induced gene expression changes are observed at the protein level and test whether there are other genes that only respond to GC post-transcriptionally. Differential gene expression related to GC exposure may contribute to GC induced IOP elevation and POAG. This study has identified differential expression of genes known to be located in IOP and POAG GWAS loci, as well as several novel genes not previously implicated in glaucoma. Future human genetic studies of GC-induced OHTN will be necessary to show that these genes contribute to GC related IOP elevation.

## Acknowledgements

Gene expression assays and analysis were provided by the Ocular Genomics Institute (OGI) Genomics Core which is funded in part by the NIH National Eye Institute Core Grant P30EY014104.

