## Supplementary Figures for "Time-dependent Glucocorticoid-Induced Transcriptomic Changes in Human Trabecular Meshwork and Schlemm’s Canal"

**Supplementary Figure 1.** Distribution of Transcript Per Million (TPM) from 2-day dexamethasone exposure experiment in (A) trabecular meshwork (TM) and (B) Schlemm’s canal endothelial (SCE) cell strains. For each sample mean TPM value is indicated by the purple color dot and median by the line in the box plot.

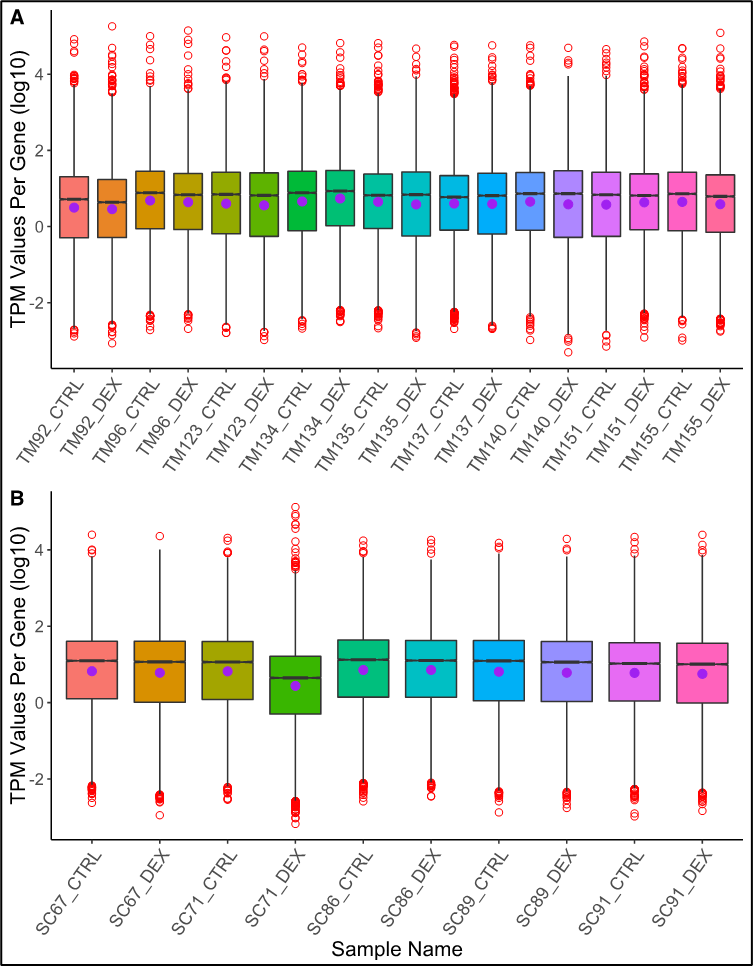

TPM=transcript per million, TM=trabecular meshwork, SC=Schlemm’s canal, CTRL=vehicle-treated cells (controls), DEX=dexamethasone-treated cell.

**Supplementary Figure 2.** Hierarchical clustering of samples two-day experiment post-DE DESeq2 normalization for (A) trabecular meshwork (TM) cell strains and (B) Schlemm’s canal endothelial (SCE) cell strains. Samples show expected paired clustering as per donor. Sub-clustering within paired samples is observed. TM samples: TM134, TM135 and TM137 formed a different sub-clade. Similarly, SC paired samples SC86 formed a distinct clade.

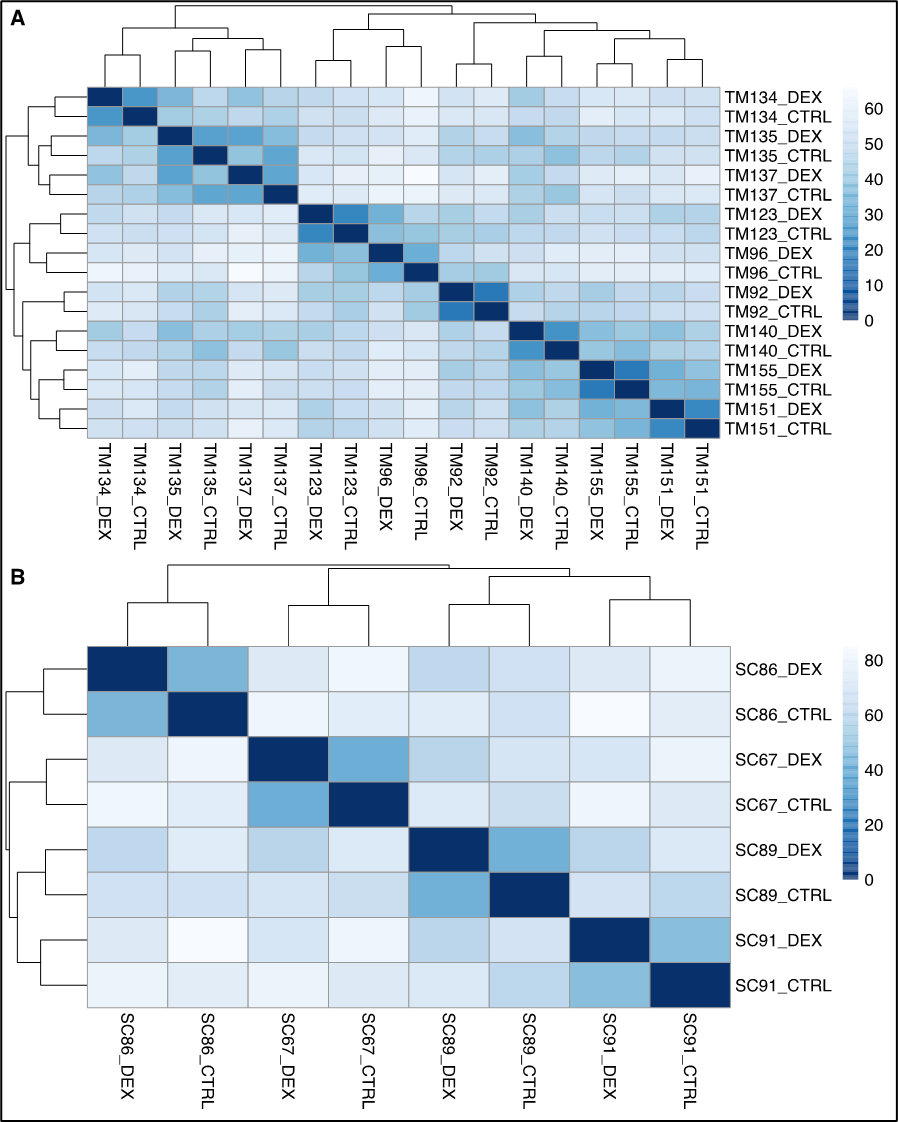

TM=trabecular meshwork, SC=Schlemm’s canal, CTRL=vehicle-treated cells (controls), DEX=dexamethasone-treated cells

**Supplementary Figure 3.** Differentially expressed genes (DEGs) of common to TM and SCE cells after two days of dexamethasone exposure. Volcano plots display log_2_(fold change) (LFC) on the *x*-axis and the -log_10_(adjusted *P*-value) on the *y*-axis for (A) trabecular meshwork (TM) cells and (B) Schlemm’s canal endothelial (SCE) cells. A total of 411 DEGs passed FDR<0.05 (206 up- and 205 down-regulated) in TM, and 11 up and 200 down-regulated in SCE. Highlighted (named) are genes with >1 LFC and q-value <0.05. For the top DEGs highlighted in each cell strain irrespective of the other cell strain see also Figure 2.

A.

.
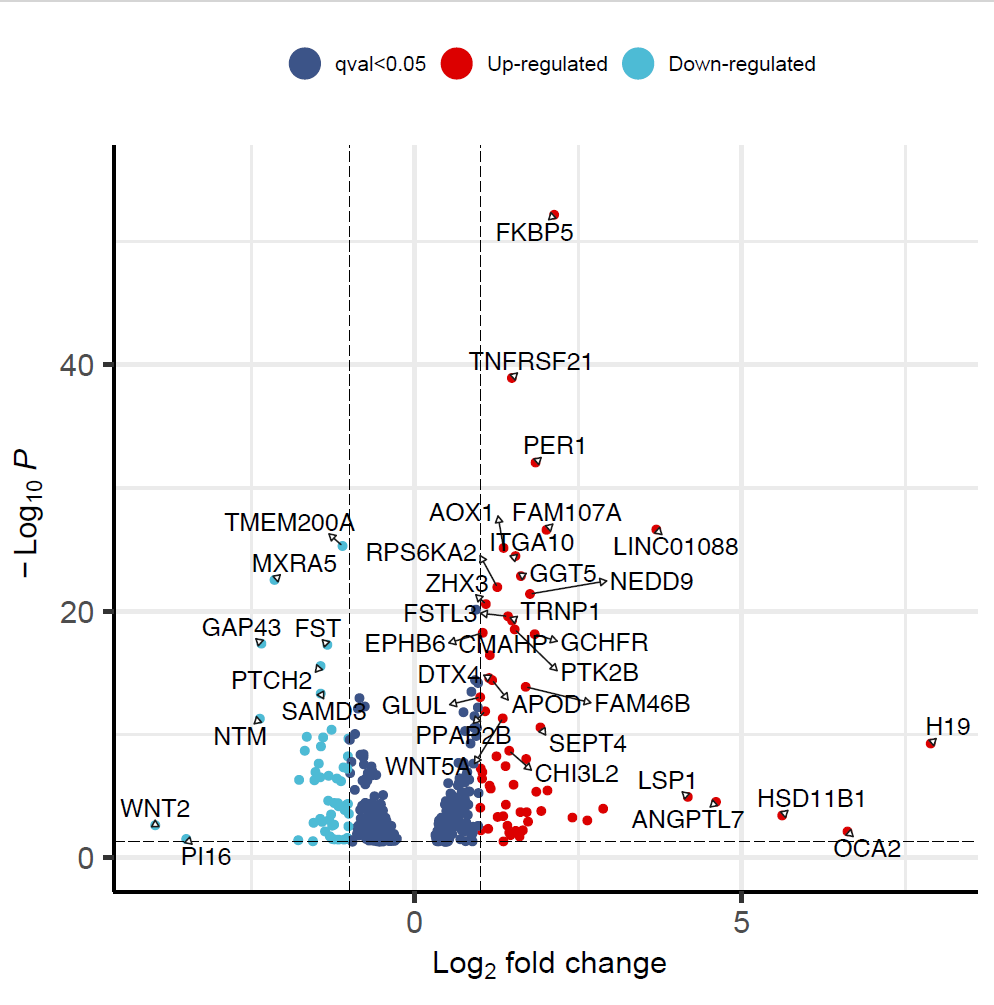

B.

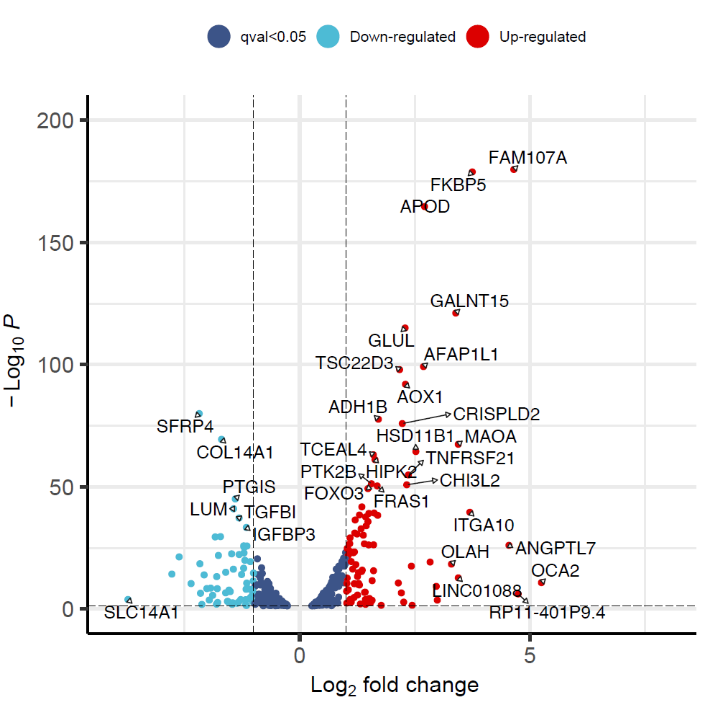

**Supplementary Figure 4.** Non-redundant gene ontology (GO) enrichment results for the differentially expressed genes in trabecular meshwork cells after 2 days of exposure to dexamethasone (DEX). Shown are (A) biological processes, (B) cellular components and (C) molecular functions. Top results with maximum 30 non-redundant GO terms are shown. (D) Bar plot of top biological processes GO terms for TM 2-day DEX exposure that pass Padj<0.05. Blue outlines of the bars specify gene sets that are only significant at Padj<0.05 for TM 2-day DEX exposure and not SCE 2-day exposure.

**A**

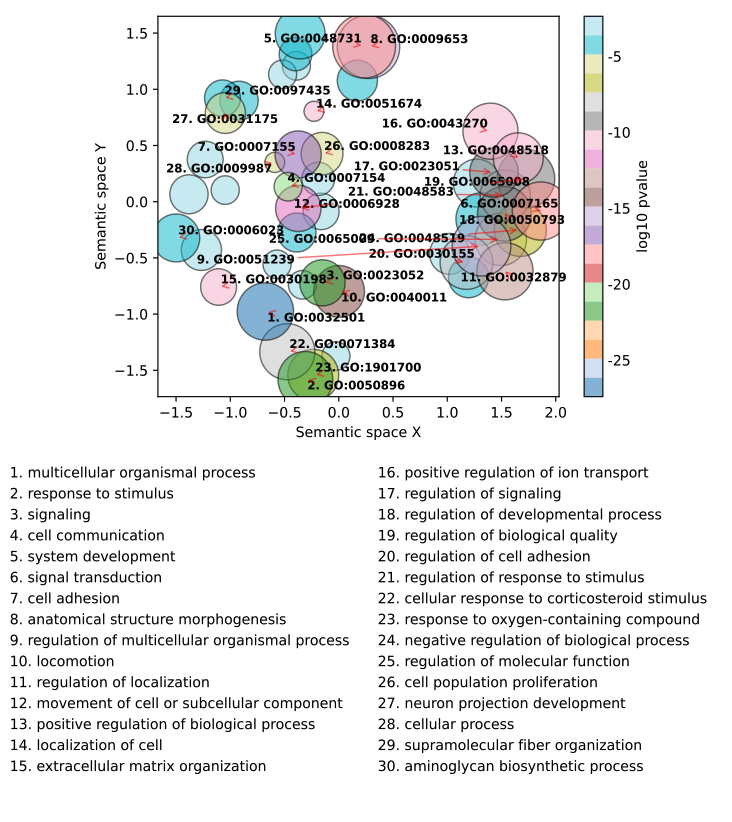

**B**

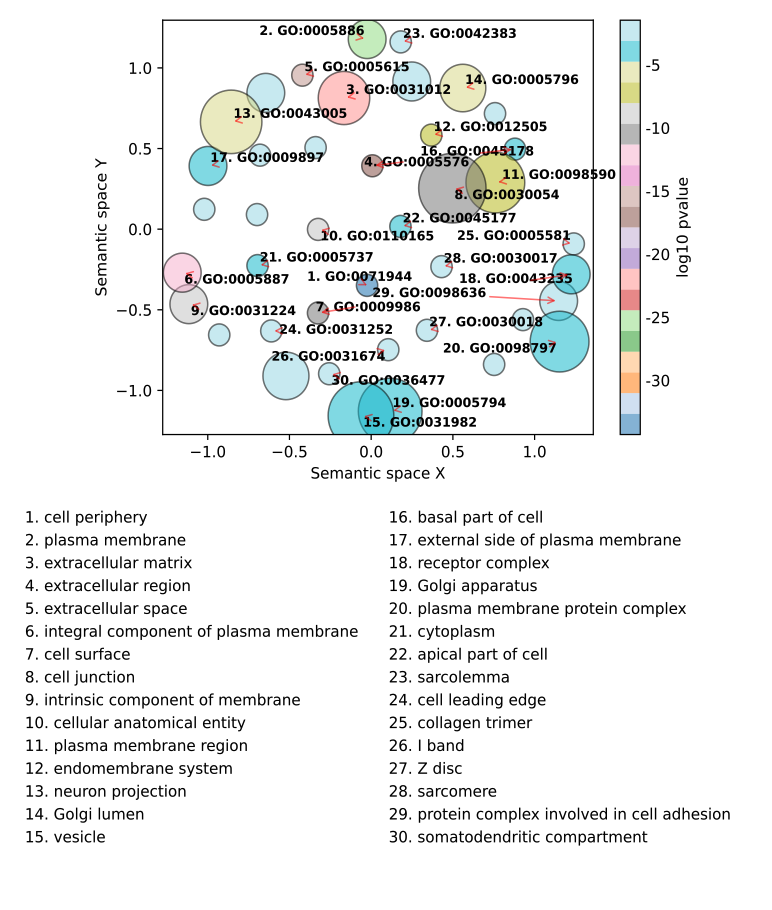

**C**

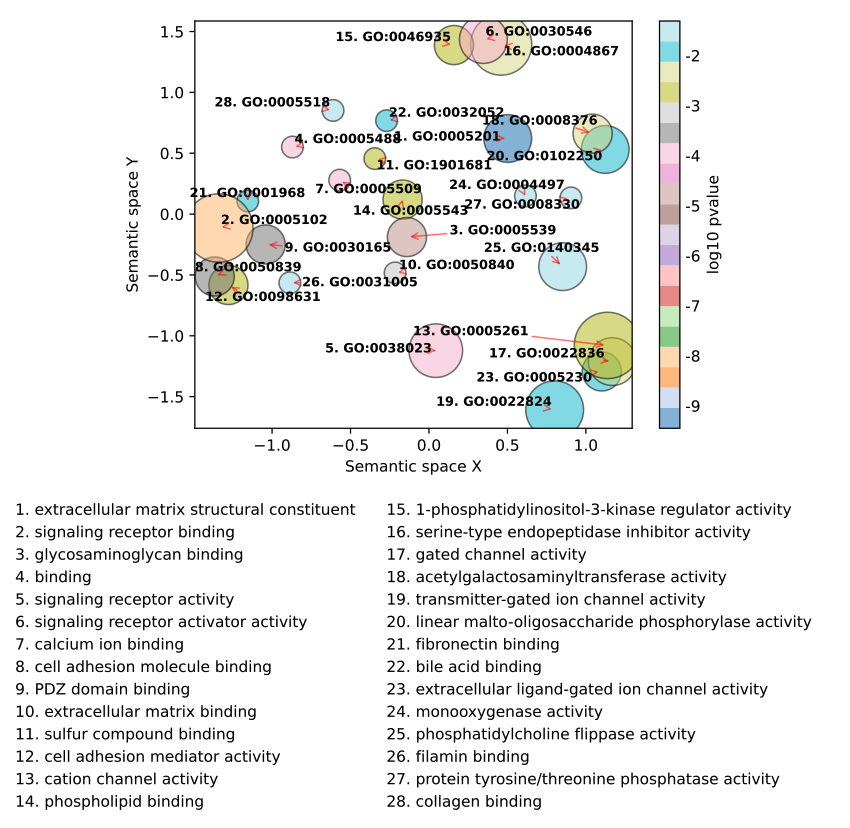

**D**

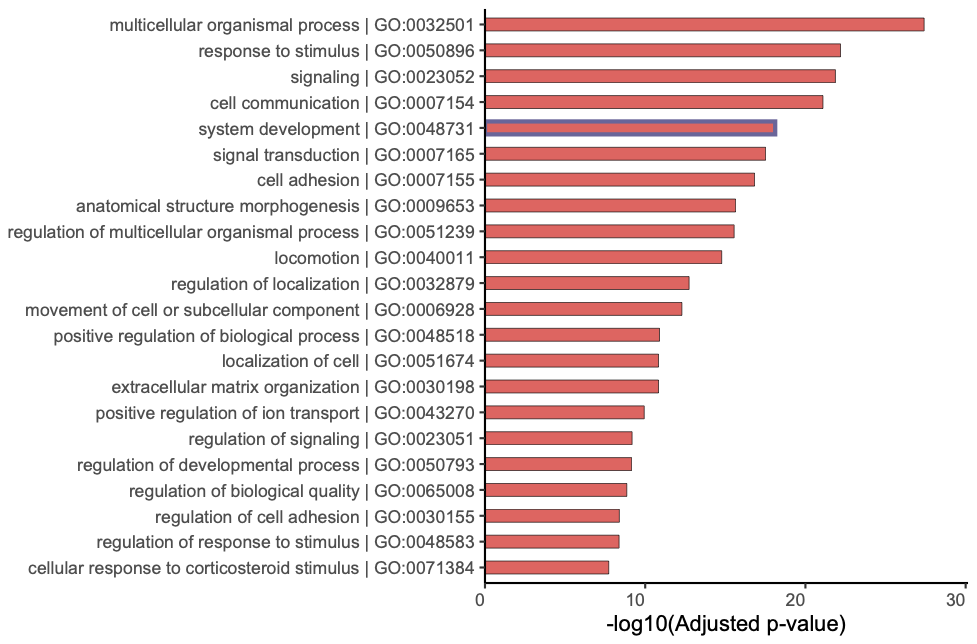

**Supplementary Figure 5.** Non-redundant gene ontology (GO) enrichment results for the differentially expressed genes in Schlemm’s canal endothelial (SCE) cells after 2 days of exposure to dexamethasone. Shown are (A) biological processes, (B) cellular components and (C) molecular functions. Top results with maximum 30 non-redundant GO terms are shown. (D) Bar plot of top biological processes GO terms for SCE 2-day DEX exposure that pass Padj<0.05. Blue outlines of the bars specify gene sets that are only significant at Padj<0.05 for SCE 2-day DEX exposure and not TM 2-day exposure.

**A**

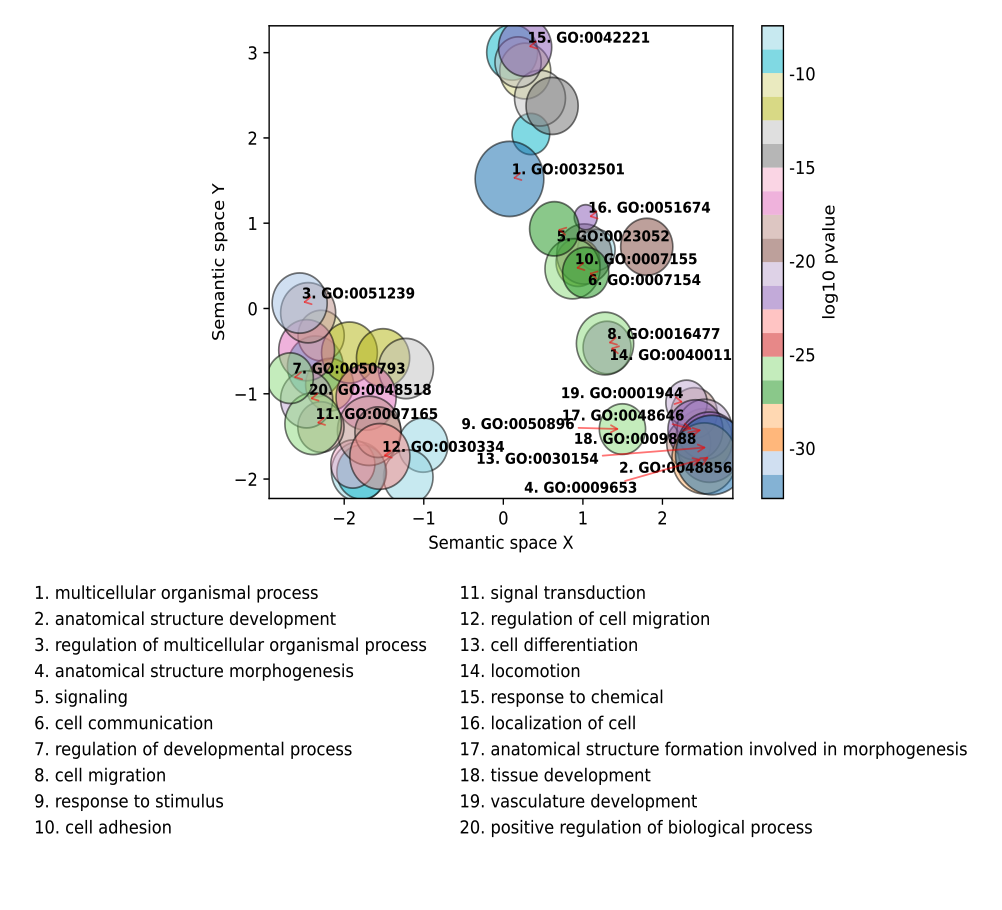

**B**

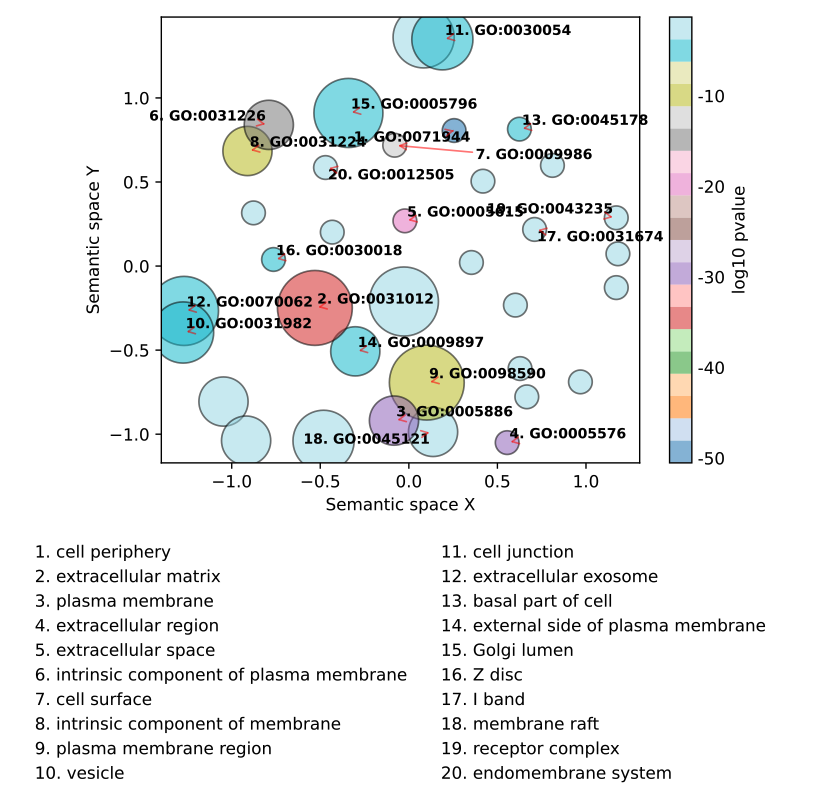

**C**

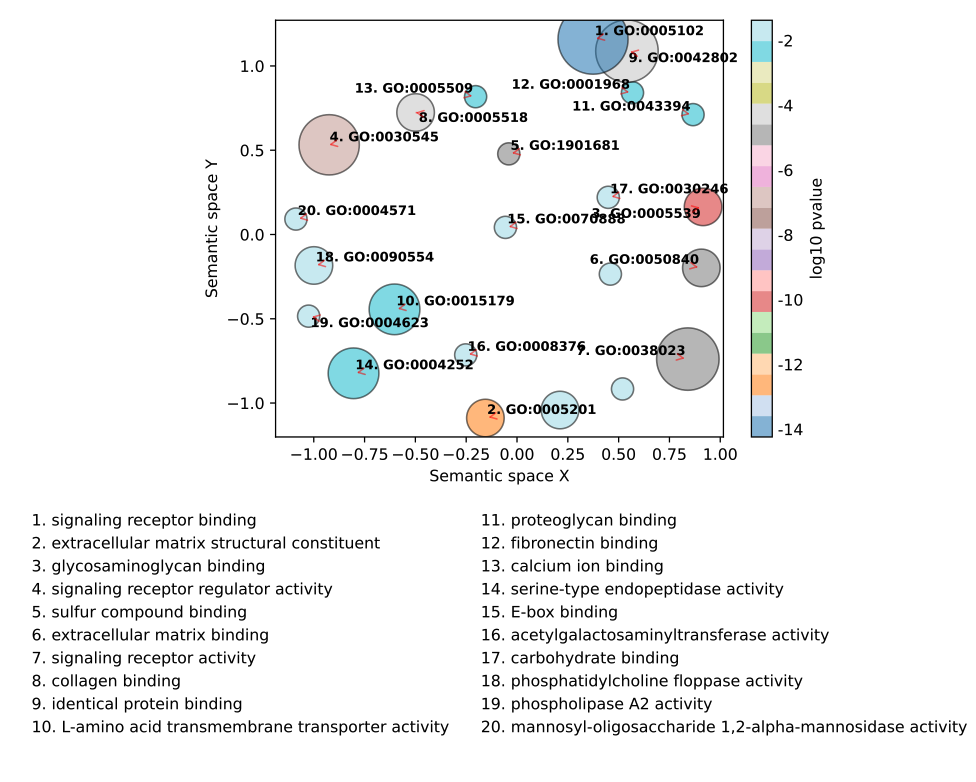

**D**

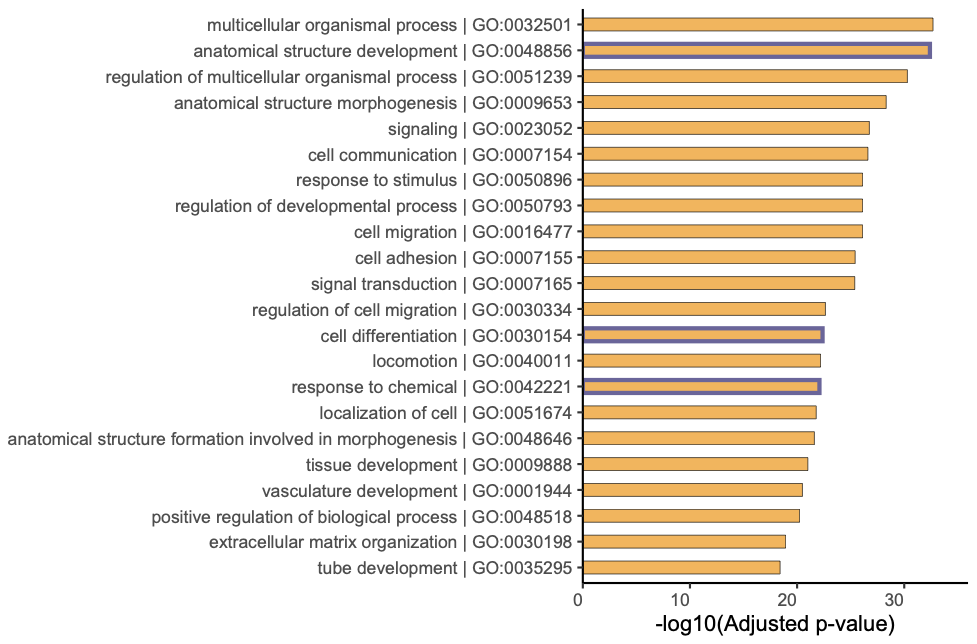

**Supplementary Figure 6.** Gene sets identified for extracellular matrix organization (gene ontology) in trabecular meshwork (TM) and Schlemm’s canal endothelial (SCE) cell strains after 2 days of exposure to dexamethasone (DEX). Heatmap shows expression patterns of 56 significant genes in TM (A) and 100 significant genes in SCE (B). Genes passing nominal (p value) and multiple hypothesis correction (qval) are shown for all three time points.

**A**

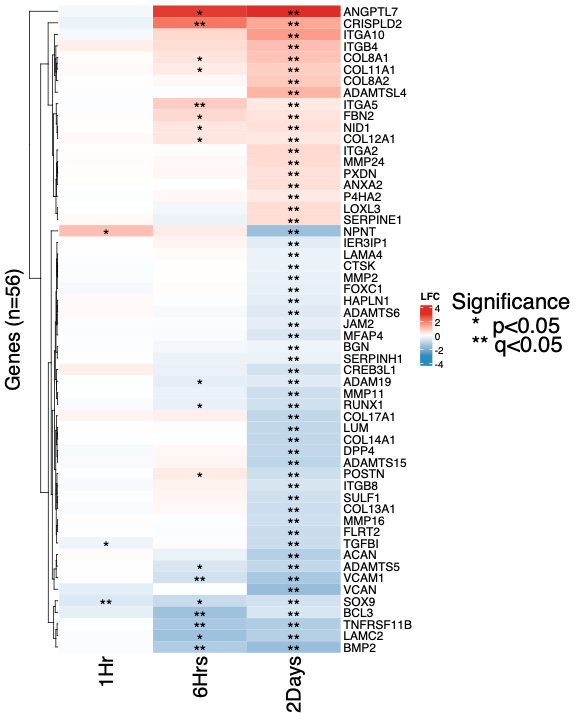

**B**

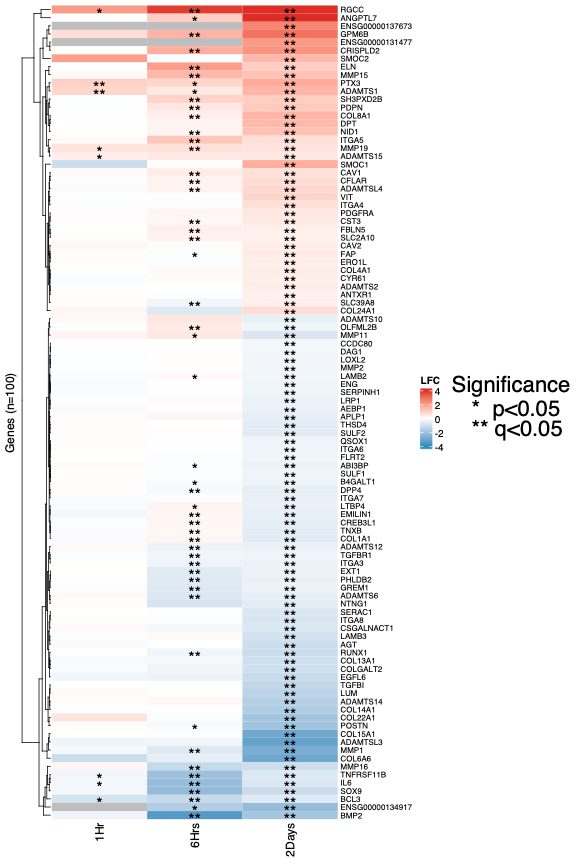

**Supplementary Figure 7.** Non-redundant gene ontology enrichment results for the common 411 differentially expressed genes in trabecular meshwork and Schlemm’s canal endothelial cells after 2 days of exposure to dexamethasone. Shown are (A) biological processes, (B) cellular components and (C) molecular functions. Top results with maximum 20 non-redundant GO terms are shown.

**A**

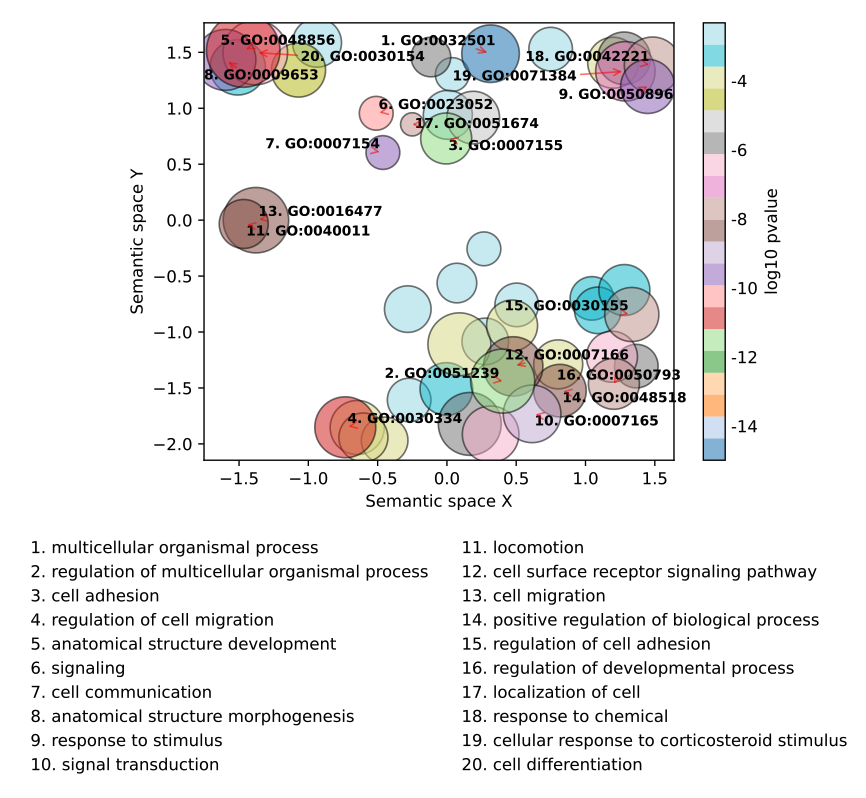

**B**

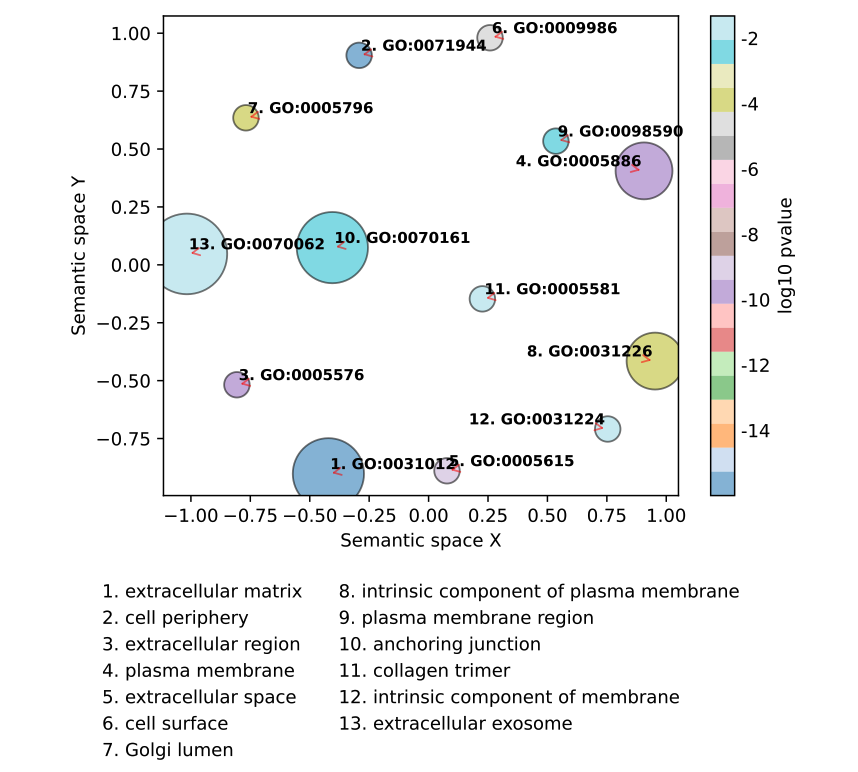

**C**

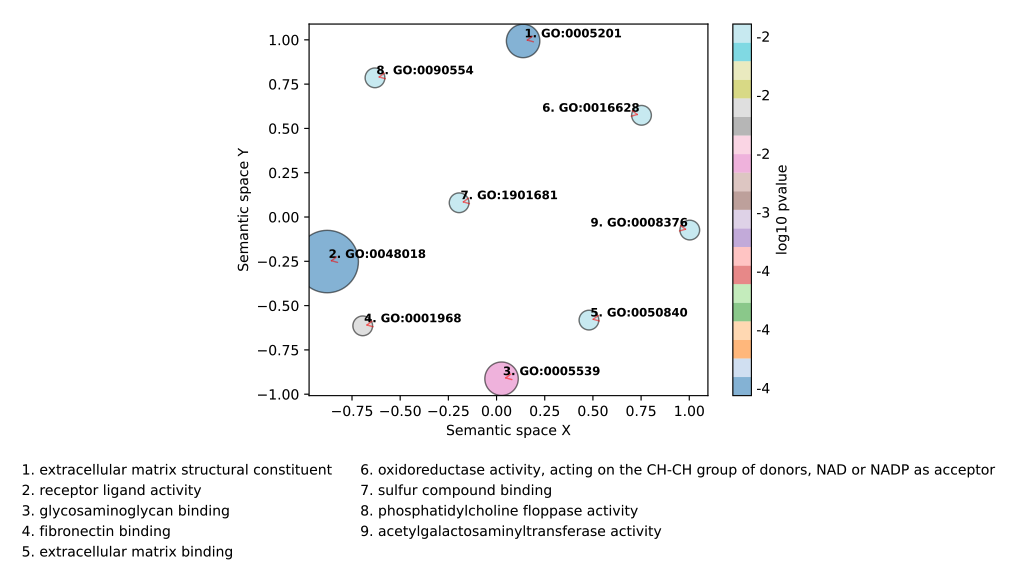

**Supplementary Figure 8.** Distribution of Transcript Per Million (TPM) from 6 hours dexamethasone exposure experiment with the (A) trabecular meshwork (TM) and (B) Schlemm’s canal endothelial (SCE) cell strains. For each sample, mean TPM value is indicated by the purple dot.

A

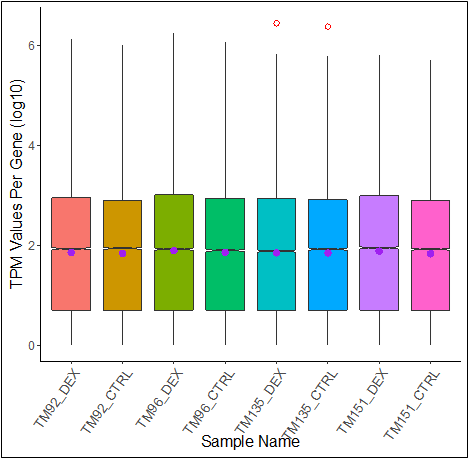

B

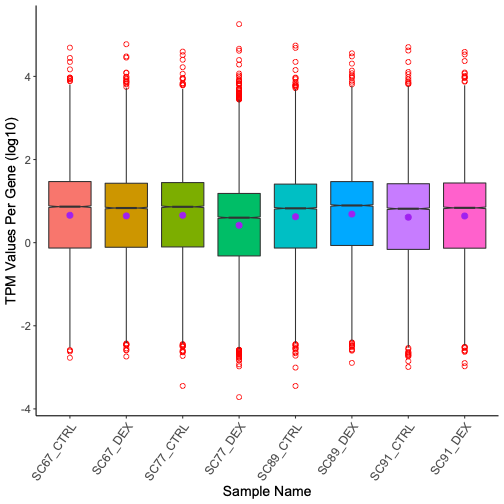

**Supplementary Figure 9.** Hierarchical clustering of samples from 6-hour dexamethasone exposure experiment post DESeq2 normalization for (A) trabecular meshwork (TM) and (B) Schlemm’s canal endothelial (SCE) cell strains. Samples show expected paired clustering per donor. Sub-clustering within paired samples is observed.

A

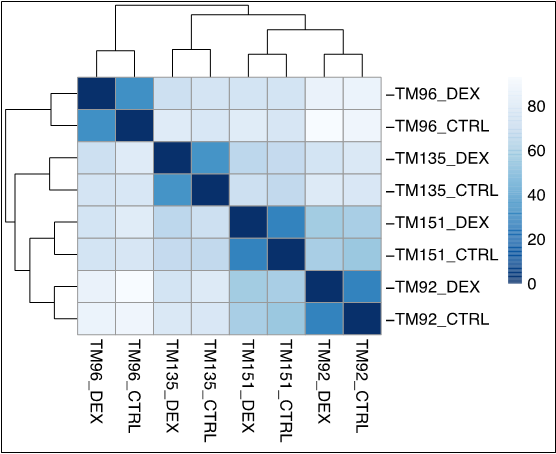

B

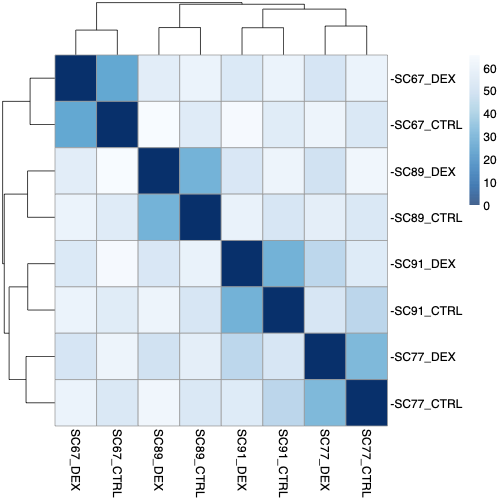

**Supplementary Figure 10.** Distribution of Transcript Per Million (TPM) from 1-hour dexamethasone exposure experiment for (A) trabecular meshwork (TM) and (B) Schlemm’s canal (SC) endothelial cell strains. For each sample mean TPM value is indicated by the purple color dot.

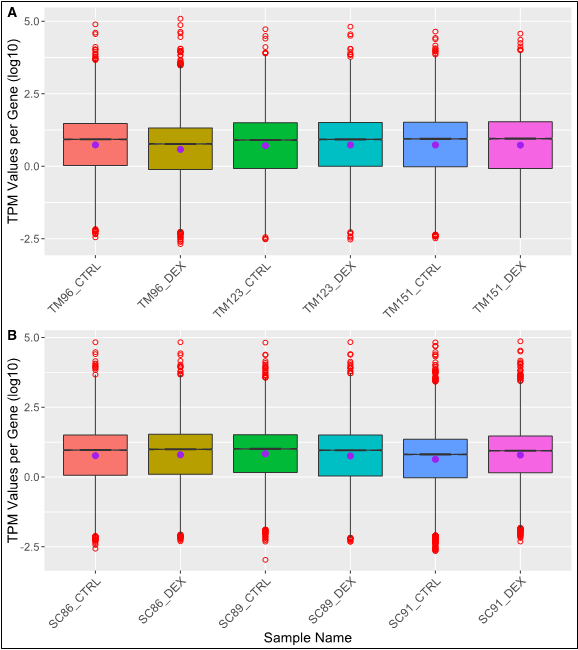

TPM=transcript per million, TM=trabecular meshwork, CTRL=vehicle-treated cells (controls), DEX=dexamethasone-treated cells, SC=Schlemm’s canal

**Supplementary Figure 11.** Hierarchical clustering of samples 1-hour experiment post normalization for (A) trabecular meshwork (TM) cell strains (B) Schlemm’s canal (SC) endothelial cell strains. Samples show expected paired clustering as per treatment. Sub-clustering within paired samples is observed. Paired TM samples 151 forming a distinct clade. Similarly paired SC samples 86 forming a subclade.

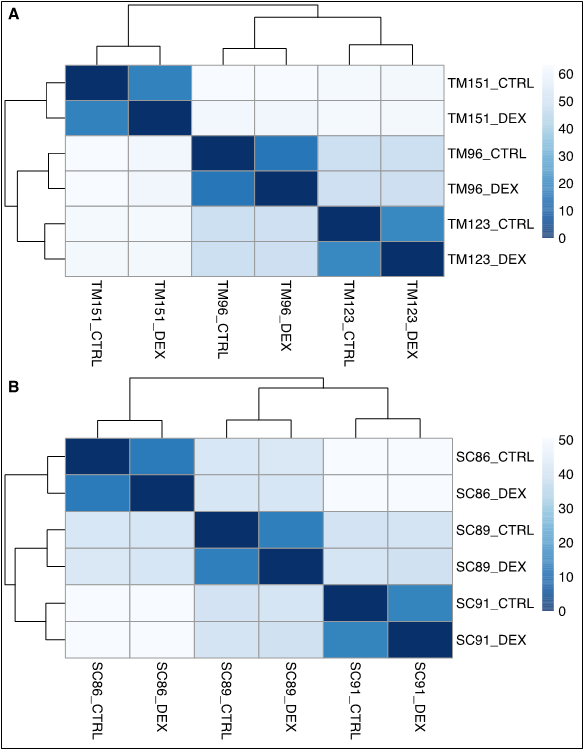

TM=trabecular meshwork, SC=Schlemm’s canal, CTRL=vehicle-treated cells (controls), DEX=dexamethasone-treated cells

**Supplementary Figure 12.** Differentially expressed genes (DEGs) after 6-hour of dexamethasone exposure on trabecular meshwork (TM) cells and Schlemm’s Canal Endothelial (SCE) cell strains. Volcano plots display log_2_(fold change) (LFC) on the x-axis and the -log_10_(adjusted P-value) on the y-axis. (A) A total 248 DEGs in TM cells were reported with 186 (75%) upregulated and 62 (24.89%) downregulated. Highlighted (named) are genes with >1.0 LFC and q-value of <0.05. (B) A total 3,208 DEGs were reported in SCE cells with 1726 (53.8%) up regulated and 1482 (46.19%) down-regulated. Highlighted (named) are genes with >2.0 LFC and q-value of <0.05.

A

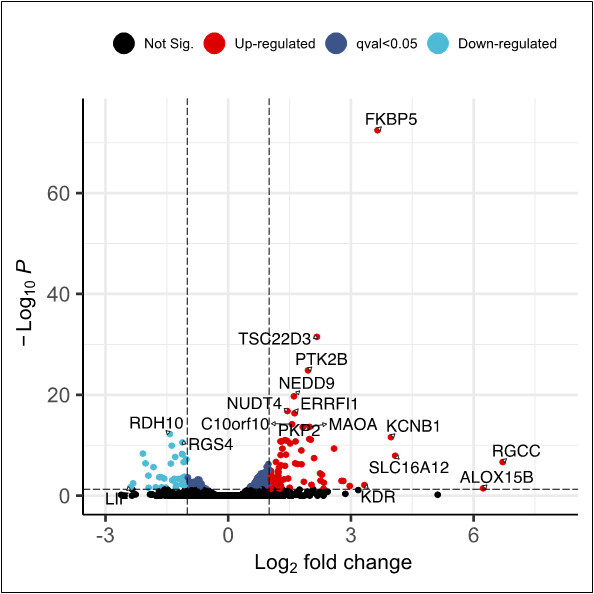

B

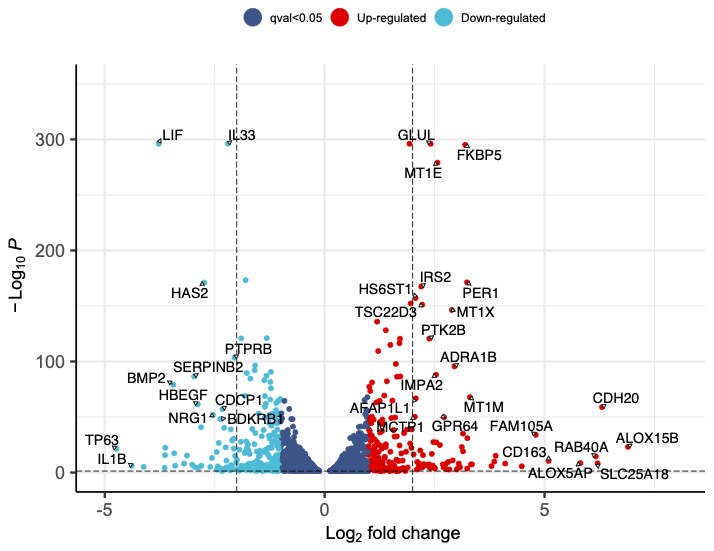

**Supplementary Figure 13.** Comparative of trabecular meshwork (TM) and Schlemm’s canal endothelial (SCE) cells differentially expressed genes (DEGs) after 6 hours of exposure to dexamethasone. (A) Venn diagram showing 202 common DEGs between TM and SCE cells exposed to dexamethasone for 6 hours. (B) Scatter plot of common DEGs to both cells lines. One hundred and forty-two (142) were collinear and up regulated while 55 were collinear and down-regulated. LFC= Log(2) fold change.

**A**

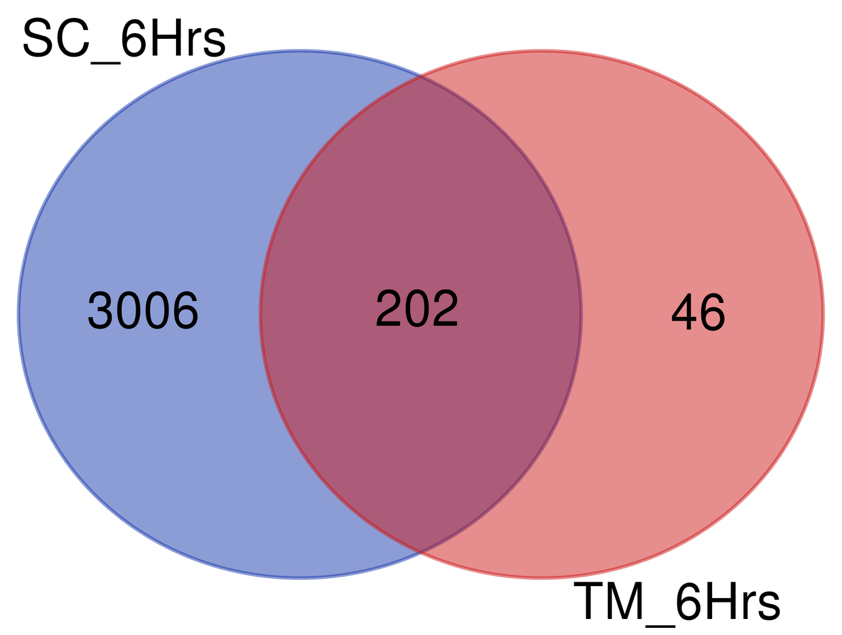
 **B**

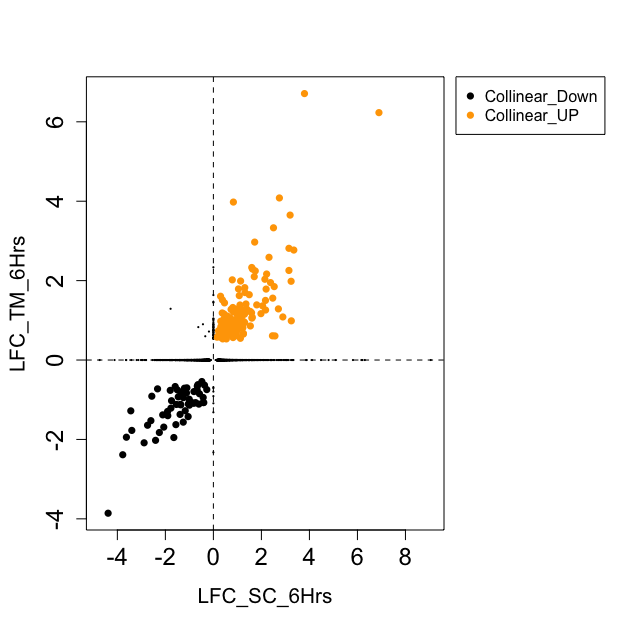

**Supplementary Figure 14.** Non-redundant gene ontology enrichment results for the differentially expressed genes in trabecular meshwork endothelial cells after 6-hours of exposure to dexamethasone. Shown are (A) biological processes, (B) cellular components and (C) molecular functions. Top results with maximum 20 non-redundant GO terms are shown.

**A**

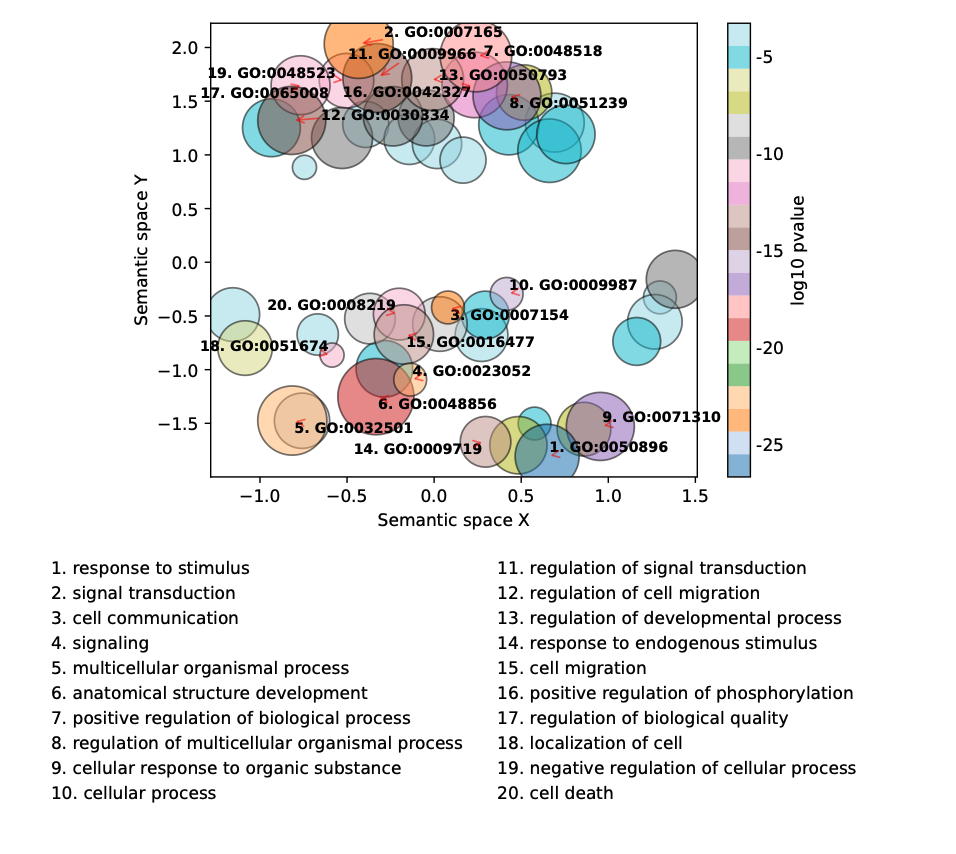

**B**

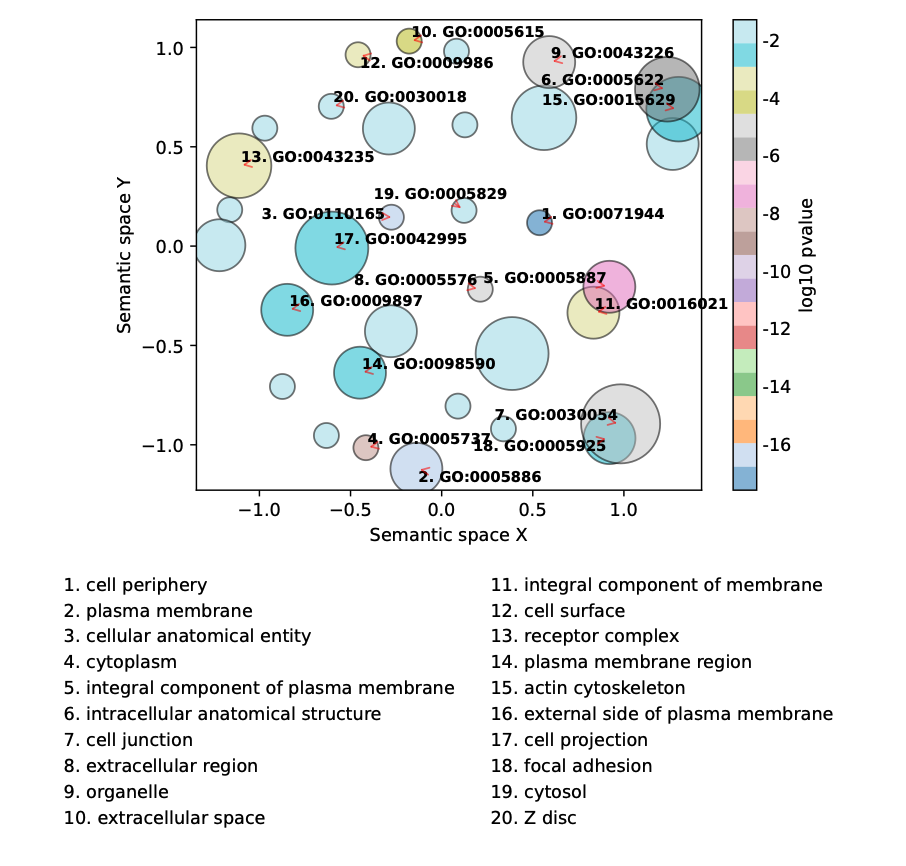

**C**

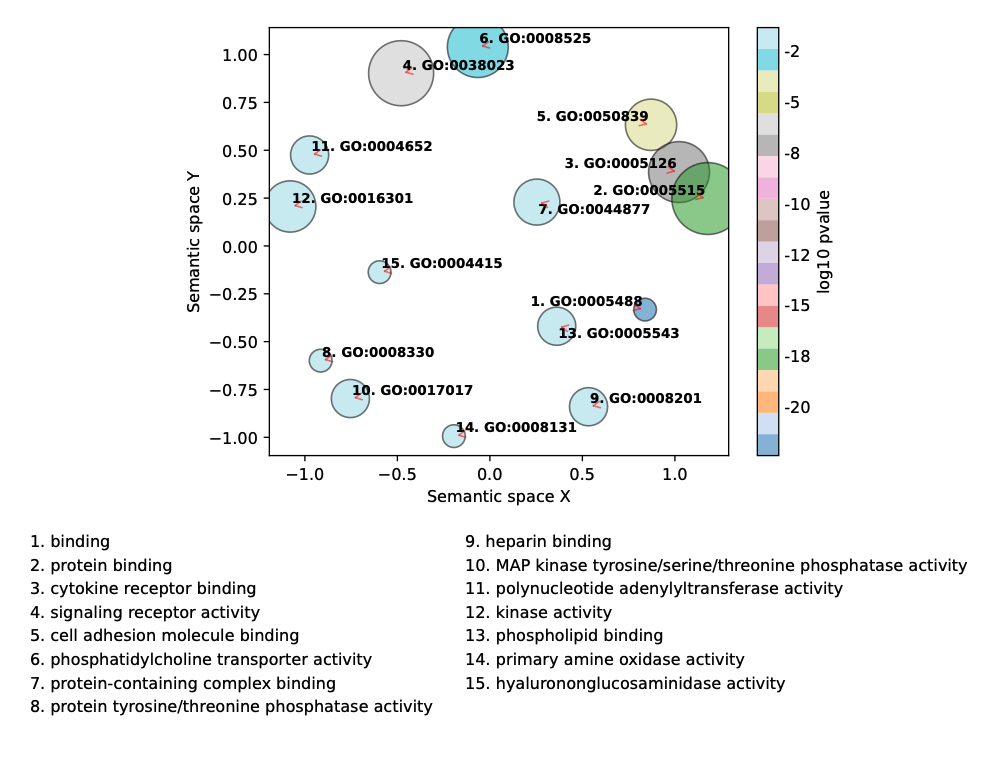

**Supplementary Figure 15.** Non-redundant gene ontology enrichment results for the differentially expressed genes in Schlemm’s canal endothelial (SCE) after 6 hours of exposure to dexamethasone. Shown are (A) biological processes, (B) cellular components and (C) molecular functions. Top results with maximum 20 non-redundant GO terms are shown.

**A**

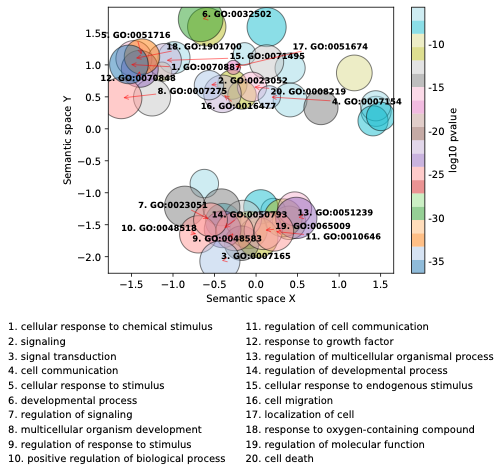

**B**

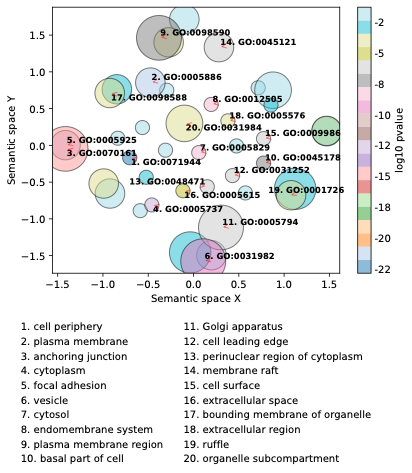

C

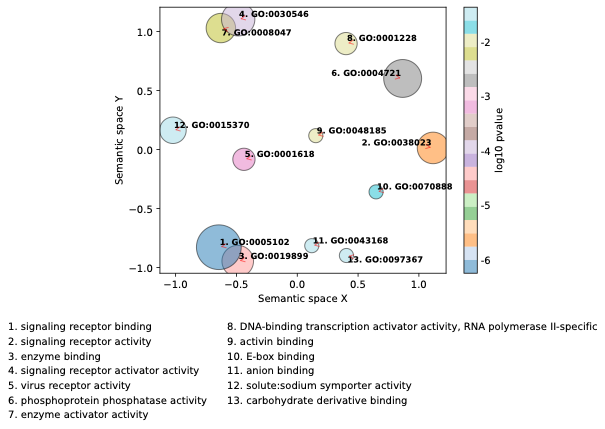

**Supplementary Figure 16.** Differentially expressed genes (DEGs) after 1-hour of dexamethasone exposure in TM and SCE cells. Volcano plots display log_2_(fold change) (LFC) on the x-axis and the -log_10_(adjusted P-value) on the y-axis (A) trabecular meshwork cells (TM) and (B) Schlemm’s canal (SC) endothelial cells. A total 33 differentially expressed genes (DEGs) in TM and 55 in SC were reported. Highlighted (named) are genes with >0.5 LFC and q-value of <0.05.

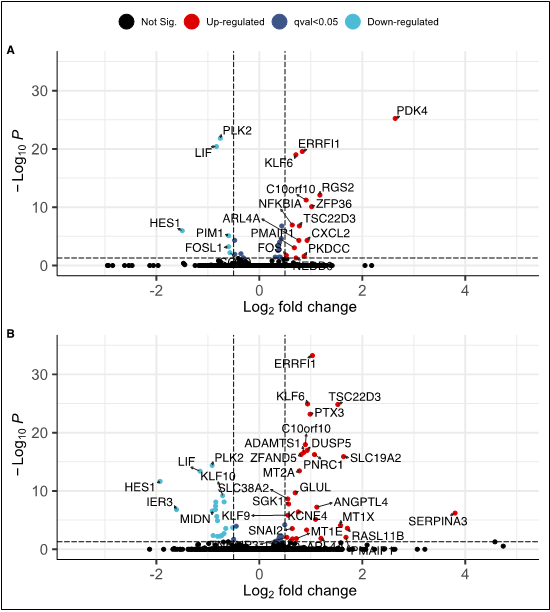

**Supplementary Figure 17.** Comparative of trabecular meshwork (TM) and Schlemm’s canal endothelial (SCE) cells differentially expressed genes (DEGs) after 1 hour of exposure to dexamethasone. (A) Venn diagram showing eighteen common DEGs between TM and SC cells exposed to dexamethasone for 1 hour. (B) Scatter plot of common DEGs to both cells lines. Twelve genes were collinear and up regulated while 6 collinear and down regulated. TM=trabecular meshwork cells, SC=Schlemm’s canal endothelial cells, DEG=differentially expressed gene, LFC= Log(2) fold change

**A**

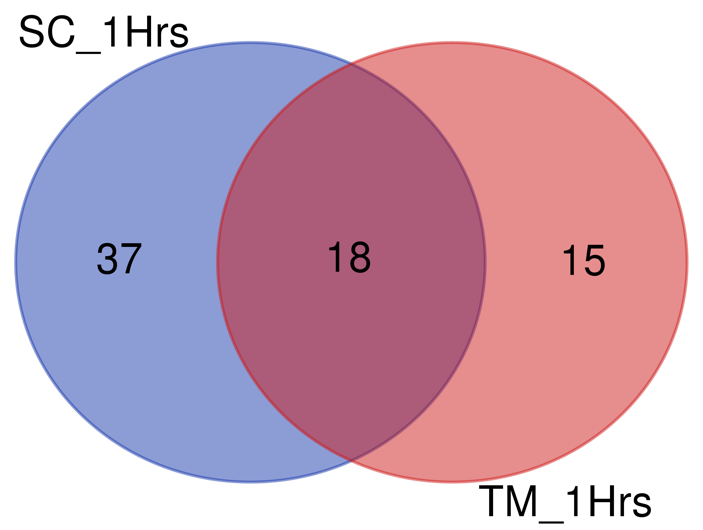

**B**

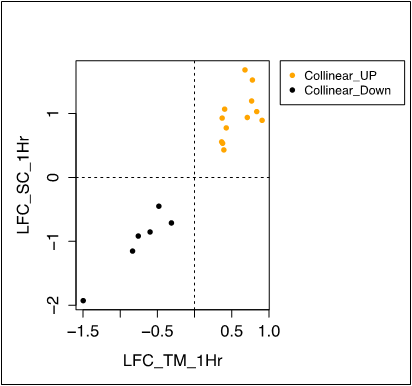

**Supplementary Figure 18.** Non-redundant gene ontology enrichment results for the differentially expressed genes in trabecular meshwork cells after 1-hour of exposure to dexamethasone. Shown are (A) biological processes, (B) cellular components and (C) molecular functions. Top results with maximum 20 non-redundant GO terms are shown.

**A**

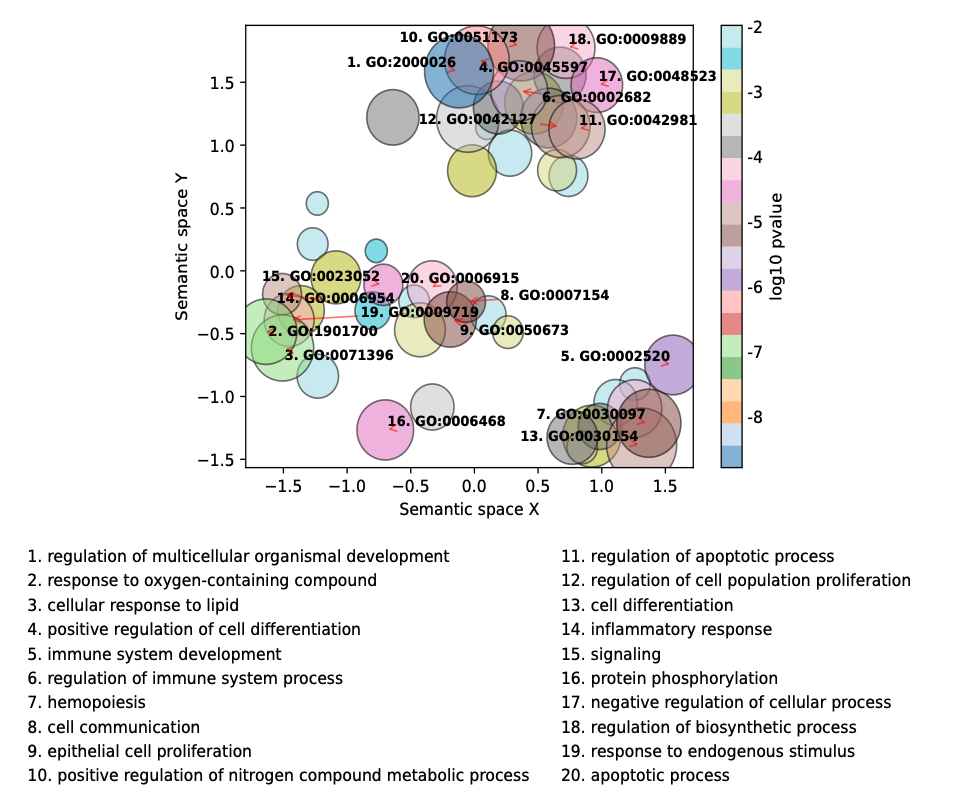

**B**

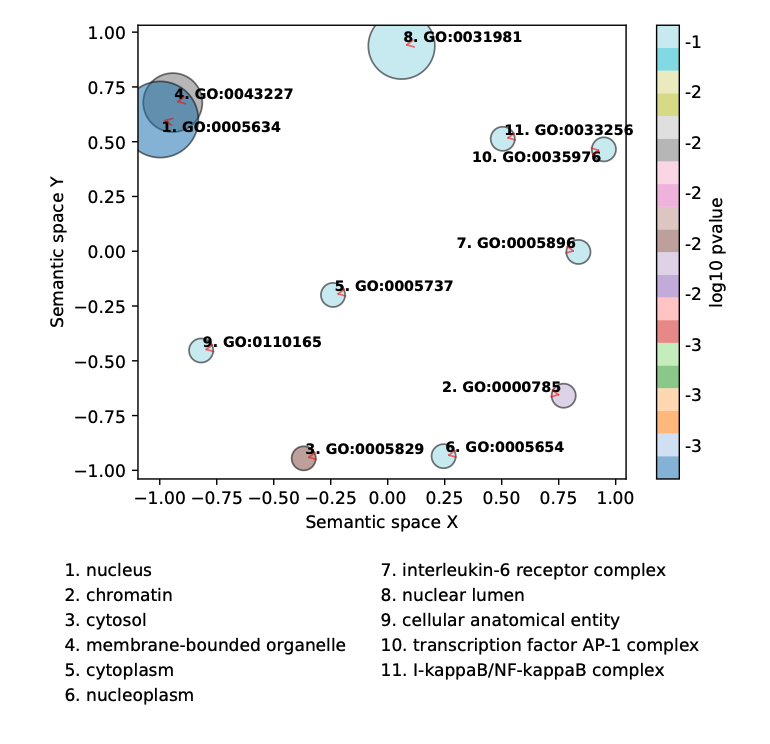

**C**

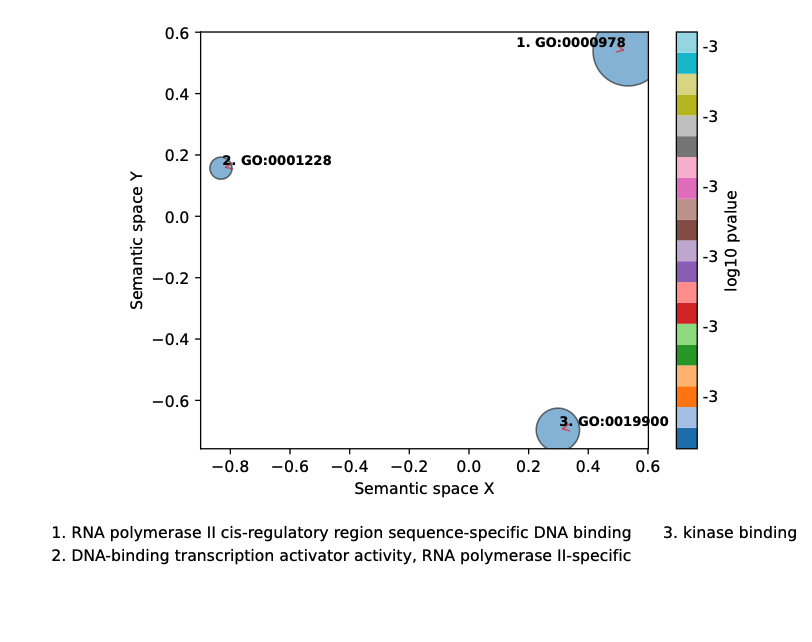

**Supplementary Figure 19**. Non-redundant gene ontology (GO) enrichment results for the differentially expressed genes in Schlemm’s canal endothelial cells after 1-hour of exposure to dexamethasone. Shown are (A) biological processes, (B) cellular components and (C) molecular functions. Top results with maximum 20 non-redundant GO terms are shown.

**A**

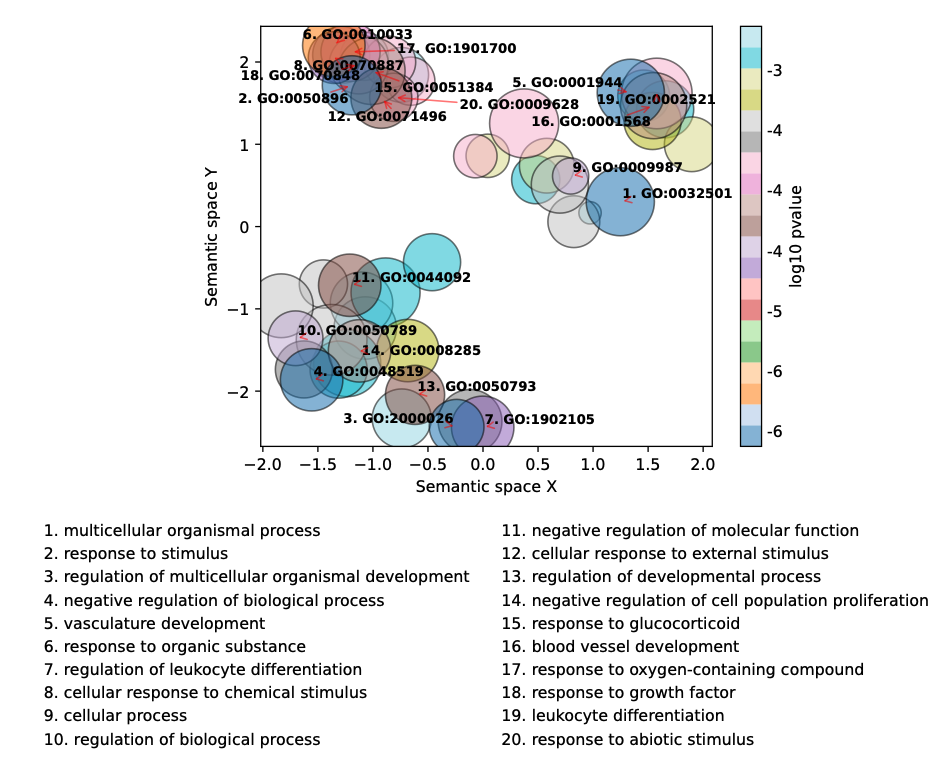

**B**

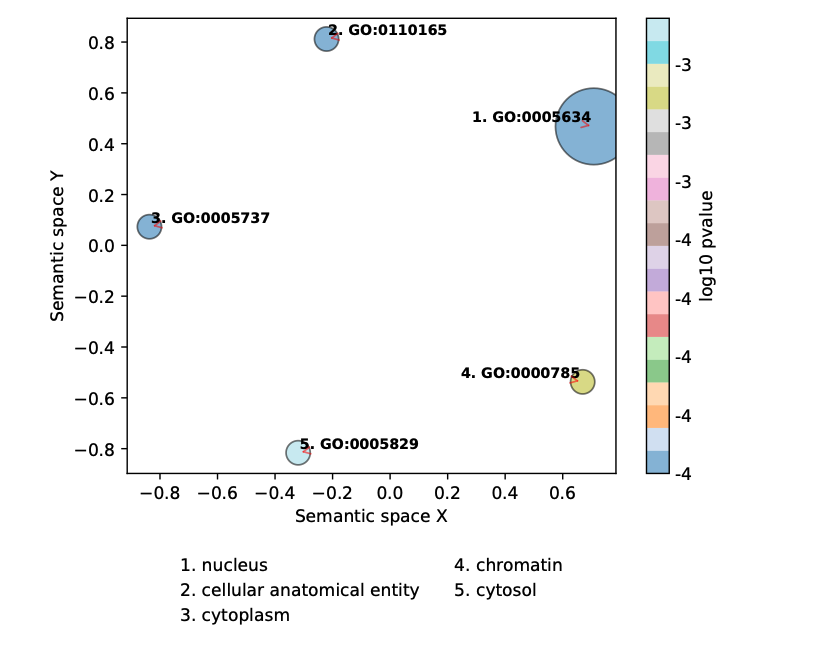

**C**

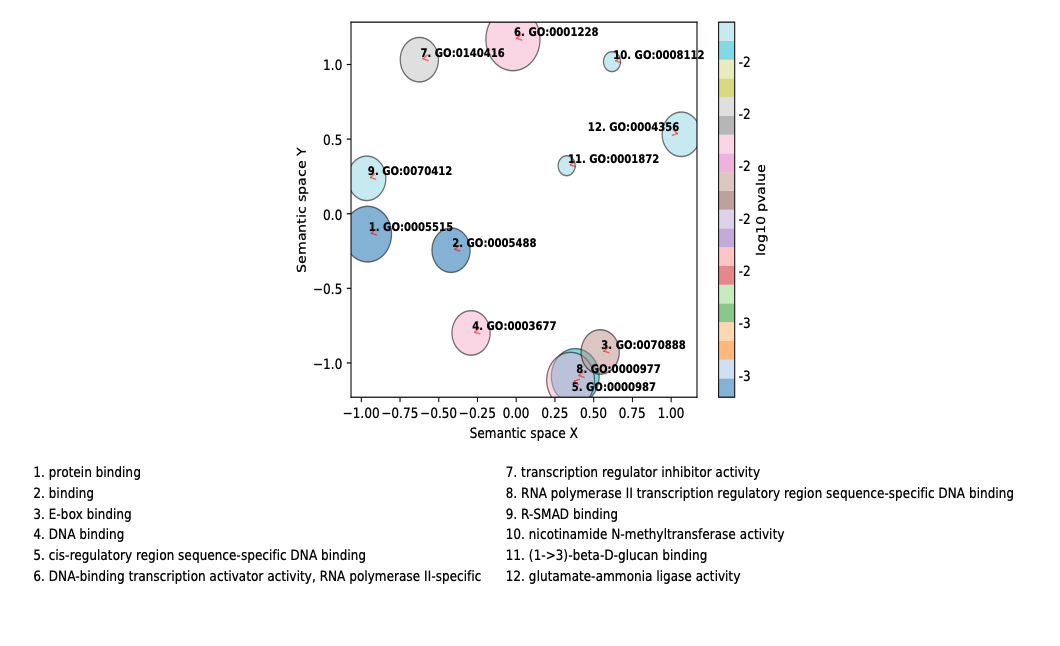

**Supplementary Figure 20.** Pearson correlation coefficient of 10 genes across 9 TM replicates using counts per million values. No outlier is observed across the samples with the selected qPCR genes.

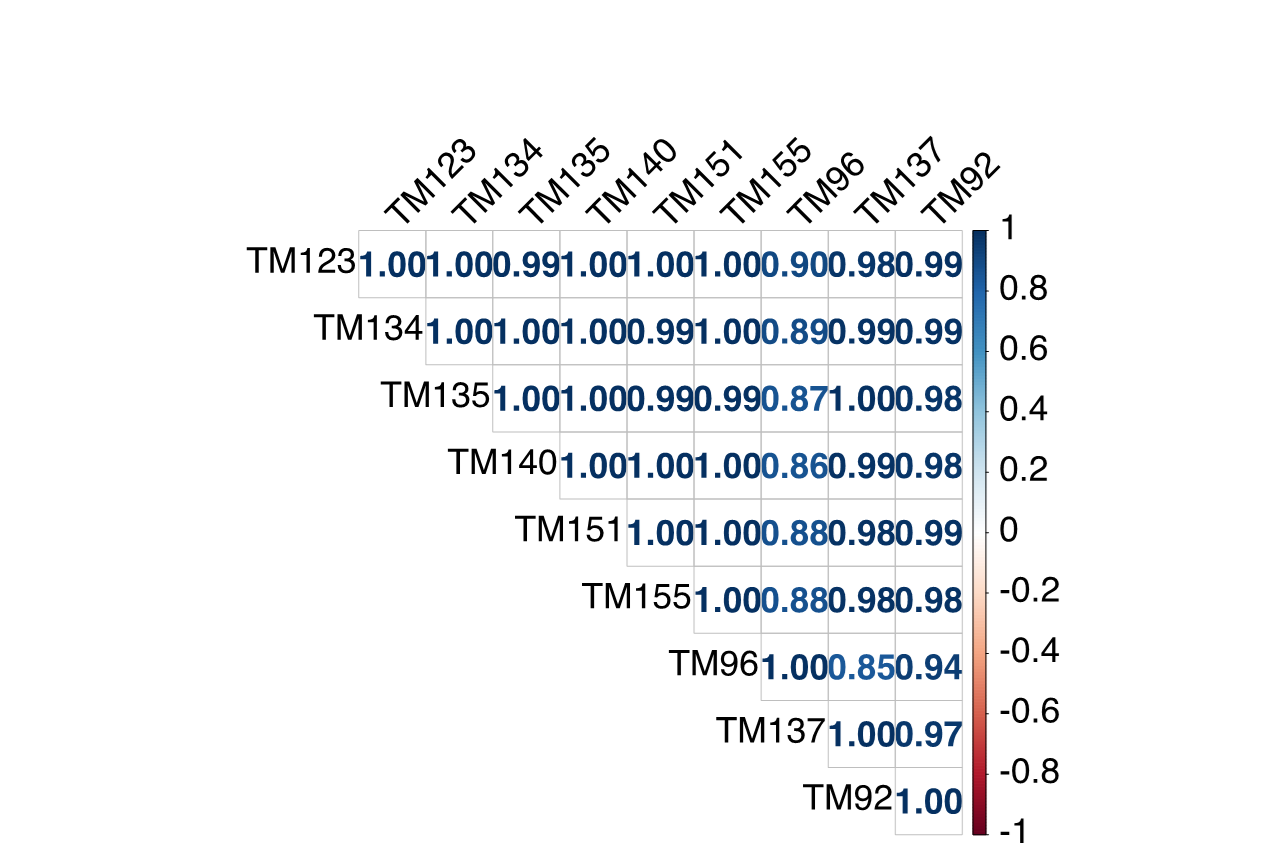
