## Supplementary Tables for "Time-dependent Glucocorticoid-Induced Transcriptomic Changes in Human Trabecular Meshwork and Schlemm’s Canal"

**Supplementary Table 1**. RNA relative quality score (RQS) across three time points. Shown are minimum, mean and 95^th^ percentile values of RQS. High quality scores of >8 was achieved across all samples for all time points.

| **Time** | **Cell Type** | **Treatment** | **Minimum RQS** | **Mean RQS** | **RQS 95^th^ Percentile** |
| --- | --- | --- | --- | --- | --- |
| 2 Days | TM | CTRL | 8.75 | 9.08 | 9.33 |
|  |  | DEX | 8.07 | 8.98 | 9.49 |
|  | SCE | CTRL | 9.0 | 9.42 | 9.7 |
|  |  | DEX | 9.1 | 9.42 | 9.7 |
| 6 Hours | TM | CTRL | 8.27 | 8.86 | 9.28 |
|  | SCE | DEX | 8.41 | 8.83 | 9.21 |
| 1 Hour | TM | CTRL | 8.4 | 8.49 | 8.53 |
|  |  | DEX | 7.8 | 8.10 | 8.37 |
|  | SCE | CTRL | 8.4 | 8.48 | 8.56 |
|  |  | DEX | 8.5 | 8.58 | 8.70 |

RNA=ribonucleic acid, TM=trabecular meshwork, SCE= Schlemm’s canal endothelium, DEX=dexamethasone-treated cells, CTRL=vehicle-treated cells (controls)

**Supplementary Table 2.** Selective quality control metrics for two-day exposure experiment. All the quality metrics obtained from ribonucleic sequencing for all the samples from trabecular meshwork (TM) and Schlemm’s canal (SC) endothelial cell strains. The sequencing was done for 101 cycles.

| **Sample ID** | **Mapping Rate** | **High Quality Rate** | **High Quality Reads** | **Mapped Reads** |
| --- | --- | --- | --- | --- |
| TM92_CTRL | 0.99462 | 0.956169 | 104,948,000 | 109,759,000 |
| TM92_DEX | 0.993804 | 0.954907 | 72,866,200 | 76,307,100 |
| TM96_CTRL | 0.995054 | 0.955187 | 74,957,200 | 78,473,900 |
| TM96_DEX | 0.992259 | 0.94336 | 67,308,700 | 71,349,900 |
| TM123_CTRL | 0.993578 | 0.950293 | 111,878,000 | 117,730,000 |
| TM123_DEX | 0.992311 | 0.943487 | 144,884,000 | 153,562,000 |
| TM134_CTRL | 0.992691 | 0.945367 | 80,232,400 | 84,869,000 |
| TM134_DEX | 0.993506 | 0.948443 | 59,266,600 | 62,488,300 |
| TM135_CTRL | 0.992655 | 0.950015 | 52,954,200 | 55,740,400 |
| TM135_CTRL | 0.992947 | 0.947352 | 150,741,000 | 159,118,000 |
| TM137_CTRL | 0.993337 | 0.951348 | 62,734,400 | 65,942,600 |
| TM137_DEX | 0.993465 | 0.9513 | 89,905,300 | 94,507,800 |
| TM140_CTRL | 0.989985 | 0.953264 | 70,930,300 | 75,565,700 |
| TM140_DEX | 0.991005 | 0.940093 | 164,780,000 | 175,281,000 |
| TM151_CTRL | 0.992839 | 0.945119 | 131,316,000 | 138,941,000 |
| TM151_DEX | 0.992071 | 0.949467 | 55,823,800 | 58,794,800 |
| TM155_CTRL | 0.993205 | 0.912776 | 70,118,300 | 76,818,700 |
| TM155_DEX | 0.993717 | 0.934635 | 71,666,200 | 76,678,200 |
| SC67_CTRL | 0.993832 | 0.950624 | 69,169,600 | 72,762,300 |
| SC67_DEX | 0.9928 | 0.95069 | 93,184,400 | 98,017,600 |
| SC71_CTRL | 0.9951 | 0.954107 | 72,262,700 | 75,738,600 |
| SC71_DEX | 0.993963 | 0.951867 | 76,167,100 | 80,018,700 |
| SC86_CTRL | 0.994193 | 0.90438 | 75,997,900 | 84,033,200 |
| SC86_DEX | 0.991596 | 0.949333 | 55,106,000 | 58,047,100 |
| SC89_CTRL | 0.991703 | 0.950389 | 89,163,500 | 93,817,900 |
| SC89_DEX | 0.990514 | 0.94854 | 87,373,600 | 92,113,800 |
| SC91_CTRL | 0.993075 | 0.949446 | 80,907,000 | 85,215,000 |
| SC91_DEX | 0.992981 | 0.951653 | 93,740,700 | 98,503,000 |

Mapping Rate: The proportion of all reads in the Bam which were mapped, and not secondary alignments or platform/vendor quality control (QC) failing reads ("Mapped Reads").

High Quality Rate: The proportion of properly paired reads with less than 6 mismatched bases and a perfect mapping quality out of all "Mapped Reads"

High Quality Reads: Mapped Reads that passed the following criteria: aligned as proper pairs, mismatches (’NM’ tag) at or below threshold, passed mapping quality (MAPQ) threshold

Mapped Reads: Unique mapping, vendor QC passed reads that were mapped

CTRL=vehicle-treated cells (controls), DEX=dexamethasone-treated cells

**Supplementary Table 3.** All statistically significant differentially expressed genes (DEGs) in trabecular meshwork (TM) cell strains after exposure to dexamethasone for 2 days. The table has various annotations including annotations from three different time points, 2 days, 6 hrs and 1 hr including Schlemm’s canal cell strains, POAG/IOP/multi-trait analysis GWAS, GSEA, biological pathways, GO, TFBSS, HP, hallmark and other genomic annotations. Please see the tab titled “SuppTable3_TM2Days_HighConfDEG” in the Excel file titled: Supplementary Excel Files.

**Supplementary Table 4.** All differentially expressed genes (DEGs) in trabecular meshwork (TM) cell strains after exposure to dexamethasone for 2 days. The table has various annotations including annotations from three different time points, 2 days, 6 hrs and 1hr including Schlemm’s canal cell strains, POAG/IOP/multi-trait analysis GWAS, GSEA, biological pathways, GO, TFBSS, HP, hallmark and other genomic annotations. Complete data on all DEGs are listed in the tab titled “SuppTable4_TM2Days_All_DEGs” in the Excel file titled: Supplementary Excel Files.

**Supplementary Table 5. Top twenty differentially expressed genes in TM cell strains following 2-day exposure to dexamethasone.** Genes are annotated for changes in TM and SCE strains at all three time points (2 days, 6 hrs and 1hr) tested, as well as for mapping to IOP and POAG associated loci based on distance to lead GWAS variants and colocalization with eQTLs and/or sQTLs in 49 GTEx tissues and retina.

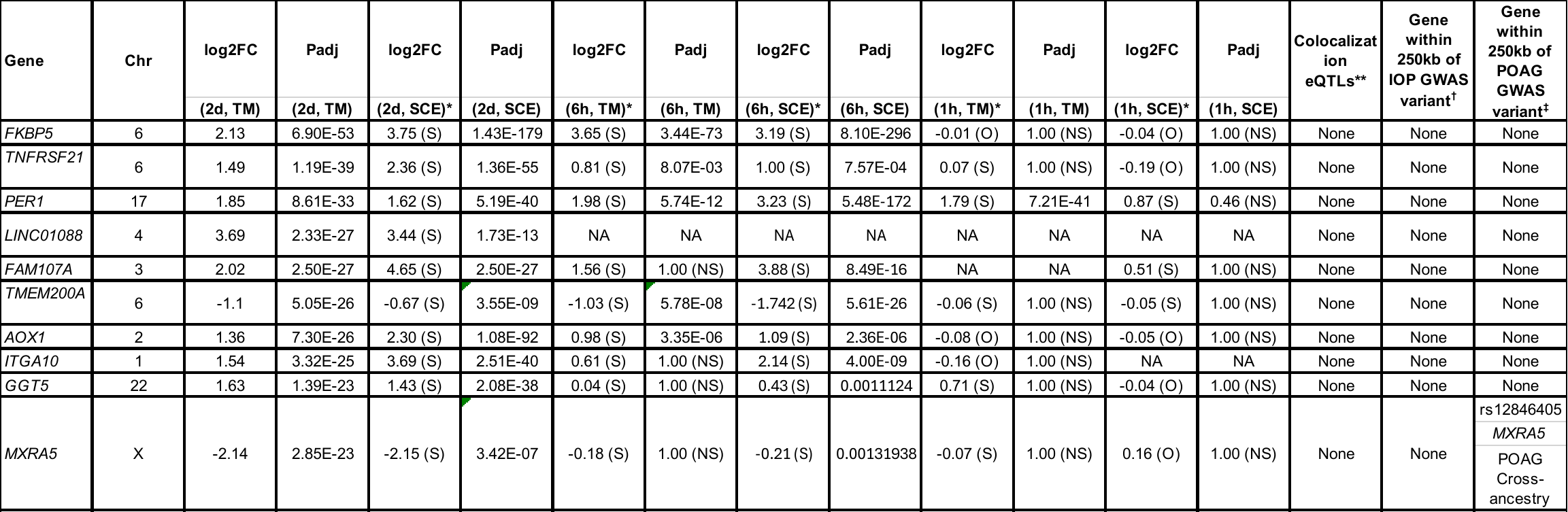

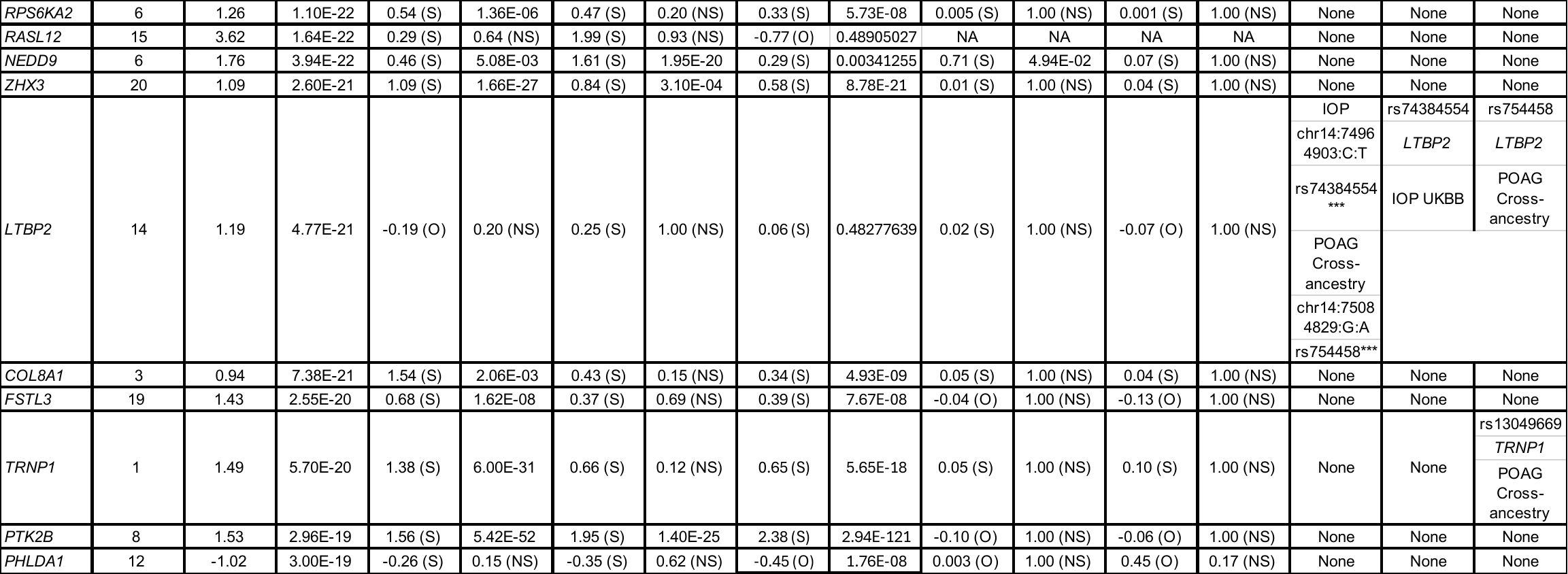

* Notation in parentheses indicates if the direction of log2FC is the same (S) or opposite (O) compared with the direction in the other TM or SCE time point experiments.

** If the differentially expressed gene is a target gene of an eQTL or sQTL previously shown to colocalize with a POAG or IOP genome-wide association study (GWAS) locus (Hamel *et al*. 2024), the lead variant of the IOP or POAG locus and the associated triait are listed.

***These eQTLs were not consistent in direction with the DEX-induced differential gene expression findings.

^†^ Genome-wide significant IOP GWAS lead variants (Khawaja *et al*, 2018) within ±250kB of the differentially expressed gene, annotated with rs ID|DEG in GWAS locus|GWAS name.

^‡^ Genome-wide significant POAG cross-ancestry GWAS lead variants within ±250kB of the differentially expressed gene, annotated with rs ID|DEG in GWAS locus|GWAS name.

Chromosome positions are in GRCh38 build.

Abbreviations: Chr= Chromosome, log2FC= log(base2) Fold-Change, 2d= 2 days dexamethasone exposure, TM= trabecular meshwork, Padj=adjusted P value, SCE= Schlemm’s Canal Endothelial Cells, NS (Non-significant), 6h= 6 hours dexamethasone exposure, 1h= 1 hour dexamethasone exposure, IOP = intraocular pressure, POAG= primary open angle glaucoma, eQTL = expression quantitative trait locus, sQTL = expression quantitative trait locus.

**Supplementary Table 6**. All statistically significant differentially expressed genes (DEGs) in Schlemm’s canal endothelial (SCE) cell strains after exposure to dexamethasone for 2 days. Annotations from all three time points; 2 days, 6 hrs and 1hr for TM and SCE strains is shown. The table is annotated various annotations including genome-wide association studies (GWAS) (intraocular pressure [IOP] and primary open-angle glaucoma [POAG]), colocalization e/sGenes from ENLOC and eCAVIAR including direction of effect from this analysis, POAG/IOP/multi-trait analysis GWAS, GSEA; biological pathways, GO, TFBSS, HP, hallmark and various genomic annotations are also included. Please see the tab titled “SuppTable6_SC2Days_HighConfDEGs” in the Excel file titled: Supplementary Excel Files.

**Supplementary Table 7**. All differentially expressed genes (DEGs) in Schlemm’s canal endothelial (SCE) cell strains after exposure to dexamethasone for 2 days. Annotations from all three time points; 2 days, 6 hrs and 1hr for TM and SCE strains is shown. The table is annotated various annotations including genome-wide association studies (GWAS) (intraocular pressure [IOP] and primary open-angle glaucoma [POAG]), colocalization e/sGenes from ENLOC and eCAVIAR including direction of effect from this analysis, POAG/IOP/multi-trait analysis GWAS, GSEA; biological pathways, GO, TFBSS, HP, hallmark and various genomic annotations are also included. Please see the tab titled “SuppTable7_SC2Days_All_DEGs” in the Excel file titled: Supplementary Excel Files.

**Supplementary Table 8. Top twenty differentially expressed genes in SCE cell strains following 2-day exposure to dexamethasone.** Genes are annotated for changes in TM and SCE strains at three the time points(2 days, 6 hrs and 1hr) tested, as well as for mapping to IOP and POAG associated loci based on distance to lead GWAS variants and colocalization with eQTLs and/or sQTLs in 49 GTEx tissues and retina.

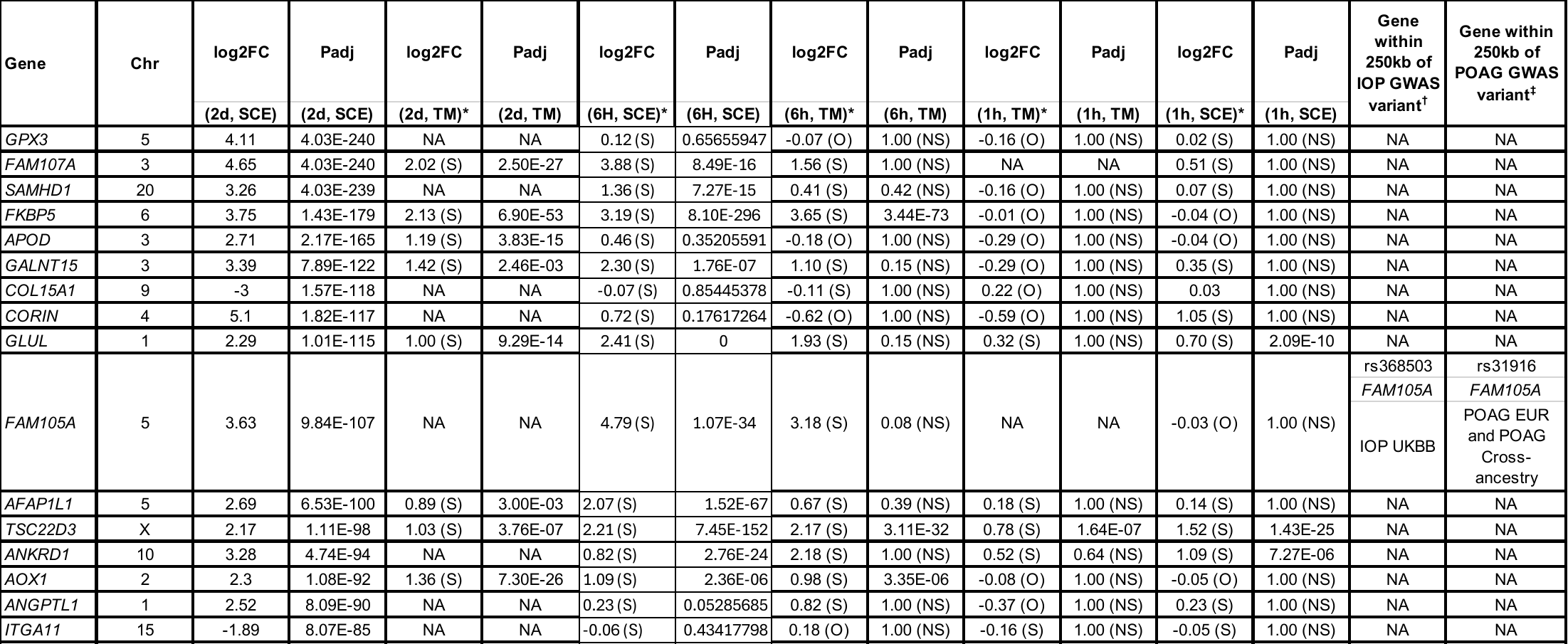

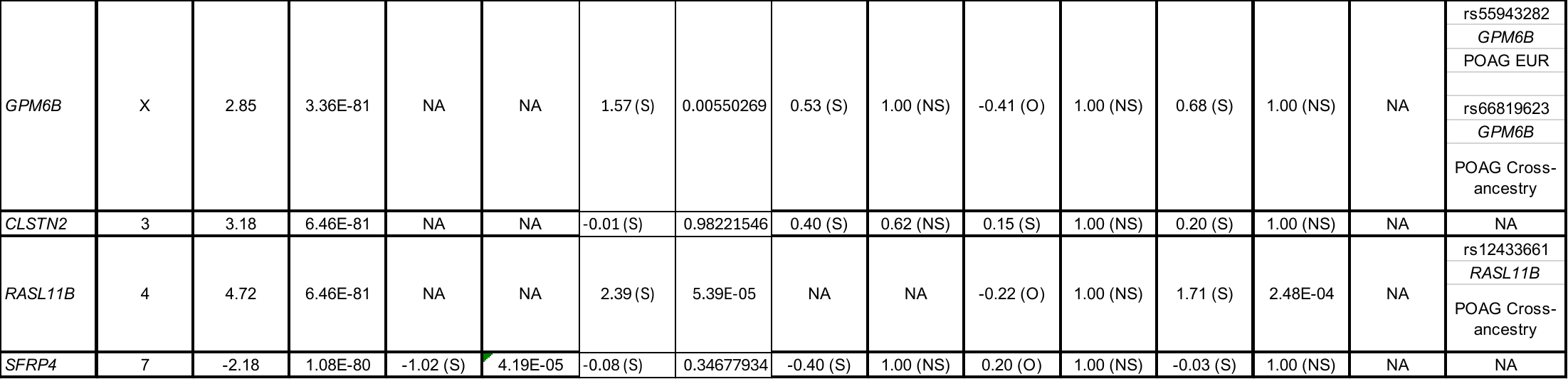

* Notation in parentheses indicates if the direction of log2FC is the same (S) or opposite (O) compared with the direction in the other TM or SCE time point experiments.

** If the differentially expressed gene is a target gene of an eQTL or sQTL previously shown to colocalize with a POAG or IOP genome-wide association study (GWAS) locus (Hamel *et al*. 2024), the lead variant of the IOP or POAG locus and the associated triait are listed.

***These eQTLs were not consistent in direction with the DEX-induced differential gene expression findings.

^†^ Genome-wide significant IOP GWAS lead variants (Khawaja *et al*, 2018) within ±250kB of the differentially expressed gene, annotated with rs ID|DEG in GWAS locus|GWAS name.

^‡^ Genome-wide significant POAG cross-ancestry GWAS lead variants within ±250kB of the differentially expressed gene, annotated with rs ID|DEG in GWAS locus|GWAS name.

Chromosome positions are in GRCh38 build.

Abbreviations: Chr= Chromosome, log2FC= log(base2) Fold-Change, 2d= 2 days dexamethasone exposure, TM= trabecular meshwork, Padj=adjusted P value, SCE= Schlemm’s Canal Endothelial Cells, NS (Non-significant), 6h= 6 hours dexamethasone exposure, 1h= 1 hour dexamethasone exposure, IOP = intraocular pressure, POAG= primary open angle glaucoma, eQTL = expression quantitative trait locus, sQTL = expression quantitative trait locus.

**Supplementary Table 9.** Common differentially expressed genes (DEGs) between trabecular meshwork (TM) and Schlemm’s canal endothelial (SCE) cell strains with 2 days dexamethasone exposure. Please see the tab titled “SuppTable9_TM-SC-2Days_CommDEGs” in the Excel file titled: Supplementary Excel Files.

**Supplementary Table 10.** All significant results for gene set enrichment analyses in Gene Ontology for gprofiler for differential gene expression (DEG) results from 2-day dexamethasone exposure to trabecular meshwork (TM) cells. Significance was determined at adjusted P < 0.05. Please see the tab titled “SuppTable10_TM2Days_GO_profiler” in the Excel file titled: Supplementary Excel Files.

**Supplementary Table 11.** All significant results for gene set enrichment analyses in Gene Ontology for biological processes for differential gene expression (DEG) results from 2-day dexamethasone exposure to trabecular meshwork (TM) cells. Significance was determined at adjusted P < 0.05. Please see the tab titled “SuppTable11_TM2Days_GO_BP” in the Excel file titled: Supplementary Excel Files.

**Supplementary Table 12.** All significant results for gene set enrichment analyses in Gene Ontology for cellular component for differential gene expression (DEG) results from 2-day dexamethasone exposure to trabecular meshwork (TM) cells. Significance was determined at adjusted P < 0.05. Please see the tab titled “SuppTable12_TM2Days_GO_CC” in the Excel file titled: Supplementary Excel Files.

**Supplementary Table 13.** All significant results for gene set enrichment analyses in Gene Ontology for molecular function for differential gene expression (DEG) results from 2-day dexamethasone exposure to trabecular meshwork (TM) cells. Significance was determined at adjusted P < 0.05. Please see the tab titled “SuppTable13_TM2Days_GO_MF” in the Excel file titled: Supplementary Excel Files.

**Supplementary Table 14.** Enriched biological pathways for the differentially expressed genes (DEGs) in trabecular meshwork (TM) cells after 2 days of exposure to dexamethasone. Significance was determined at adjusted P < 0.05. Top 10 statistically most significant enriched pathways from KEGG and REACTOME database are shown. Selective columns are shown.

| **Source** | **Term name** | **Term id** | **Adjusted p-value** |
| --- | --- | --- | --- |
| REAC | Extracellular matrix organization | REAC:R-HSA-1474244 | 1.59E-09 |
| REAC | Signaling by GPCR | REAC:R-HSA-372790 | 5.01E-08 |
| REAC | Signal Transduction | REAC:R-HSA-162582 | 2.61E-06 |
| REAC | GPCR downstream signaling | REAC:R-HSA-388396 | 1.62E-05 |
| REAC | G alpha (i) signaling events | REAC:R-HSA-418594 | 2.35E-05 |
| REAC | GPCR ligand binding | REAC:R-HSA-500792 | 2.35E-05 |
| REAC | Class B/2 (Secretin family receptors) | REAC:R-HSA-373080 | 0.0003711 |
| KEGG | PI3K-Akt signaling pathway | KEGG:04151 | 0.00057936 |
| KEGG | Calcium signaling pathway | KEGG:04020 | 0.00057936 |

GPCR=G protein-coupled receptor

**Supplementary Table 15.** Enriched biological pathways for the differentially expressed genes (DEGs) in trabecular meshwork (TM) cells after 2 days of exposure to dexamethasone. Significance was determined at adjusted P < 0.05. Please see the tab titled “SuppTable15_TM2Days_Pathways” in the Excel file titled: Supplementary Excel Files for the complete table.

**Supplementary Table 16.** Enriched hallmark gene sets for the differentially expressed genes (DEGs) in trabecular meshwork (TM) cells after 2 days of exposure to dexamethasone. Significance was determined at adjusted P < 0.05. Top 10 statistically most significant enriched gene sets are shown. Selective columns are shown.

| **Term** | **Description** | **Log P** | **Log (q-value)** |
| --- | --- | --- | --- |
| M5930 | HALLMARK EPITHELIAL MESENCHYMAL TRANSITION | -19.23314 | -17.534 |
| M5890 | HALLMARK TNFA SIGNALING VIA NFKB | -9.5621731 | -8.164 |
| M5953 | HALLMARK KRAS SIGNALING UP | -7.5672855 | -6.345 |
| M5907 | HALLMARK ESTROGEN RESPONSE LATE | -6.9804317 | -5.980 |
| M5942 | HALLMARK UV RESPONSE DN | -6.9804317 | -5.980 |
| M5891 | HALLMARK HYPOXIA | -6.2753241 | -5.355 |
| M5947 | HALLMARK IL2 STAT5 SIGNALING | -5.0157107 | -4.163 |
| M5909 | HALLMARK MYOGENESIS | -4.9592999 | -4.163 |
| M5915 | HALLMARK APICAL JUNCTION | -3.1807832 | -2.523 |
| M5909 | HALLMARK MYOGENESIS | -4.9592999 | -4.163 |

**Supplementary Table 17.** Enriched hallmark gene sets for the differentially expressed genes (DEGs) in trabecular meshwork (TM) cells after 2 days of exposure to dexamethasone. Significance was determined at adjusted P < 0.05. Please see the tab titled “SuppTable17_TM2Days_Hallmark” in the Excel file titled: Supplementary Excel Files for the complete table.

**Supplementary Table 18.** Enriched transcription factor binding sites for the differentially expressed genes (DEGs) in trabecular meshwork (TM) cells after 2 days of exposure to dexamethasone. Significance was determined at adjusted P < 0.05. Top 10 statistically significant enrichment transcription factors (TF) are shown.

| **Source** | **Term name** | **Term id** | **Adjusted p value** |
| --- | --- | --- | --- |
| TF | Factor: TIEG1; motif: NCCCNSNCCCCGCCCCC | TF:M12351 | 8.94E-21 |
| TF | Factor: MOVO-B; motif: GNGGGGG | TF:M01104 | 8.94E-21 |
| TF | Factor: MAZ; motif: GGGGGAGGGGGNGRGRRRGNRG | TF:M09984 | 4.05E-19 |
| TF | Factor: GKLF; motif: NNNRGGNGNGGSN | TF:M07289 | 4.49E-17 |
| TF | Factor: BTEB3; motif: CCNNSCCNSCCCCKCCCCC | TF:M09826 | 2.41E-16 |
| TF | Factor: ETF; motif: GVGGMGG | TF:M00695 | 3.01E-16 |
| TF | Factor: Churchill; motif: CGGGNN | TF:M00986 | 5.90E-16 |
| TF | Factor: MAZ; motif: GGGMGGGGS | TF:M10432 | 5.76E-15 |
| TF | Factor: AP-2alpha; motif: NGCCYSNNGSN | TF:M01857 | 1.31E-14 |

**Supplementary Table 19.** Enriched transcription factor binding sites for the differentially expressed genes (DEGs) in trabecular meshwork (TM) cells after 2 days of exposure to dexamethasone. Significance was determined at adjusted P < 0.05. Please see the tab titled “SuppTable19_TM2Days_TFBS” in the Excel file titled: Supplementary Excel Files for the complete table.

**Supplementary Table 20.** Enriched human phenotype ontology for the differentially expressed genes (DEGs) in trabecular meshwork (TM) cells after 2 days of exposure to dexamethasone. Significance was determined at adjusted P < 0.05. Only one statistically significant enrichment was observed and shown below.

| **Source** | **Term name** | **Term id** | **Adjusted p value** |
| --- | --- | --- | --- |
| HP | Autosomal dominant inheritance | HP:0000006 | 0.0329851 |

**Supplementary Table 21.** Enriched human phenotype ontology for the differentially expressed genes (DEGs) in trabecular meshwork (TM) cells after 2 days of exposure to dexamethasone. Significance was determined at adjusted P < 0.05. Please see the tab titled “SuppTable21_TM2Days_HPO” in the Excel file titled: Supplementary Excel Files for the complete table.

**Supplementary Table 22.** All significant results for gene set enrichment analyses in Gene Ontology for gprofiler of differential gene expression (DEGs) results from 2-day dexamethasone exposure to Schlemm’s canal endothelial (SCE) cells. Significance was determined at adjusted P < 0.05. Please see the tab titled “SuppTable22_SC2Day_GO_gprofiler” in the Excel file titled: Supplementary Excel Files.

**Supplementary Table 23.** All significant results for gene set enrichment analyses in Gene Ontology of biological processes of differential gene expression (DEGs) results from 2-day dexamethasone exposure to Schlemm’s canal endothelial (SCE) cells. Significance was determined at adjusted P < 0.05. Please see the tab titled “SuppTable23_SC2Days_GO_BP” in the Excel file titled: Supplementary Excel Files.

**Supplementary Table 24.** All significant results for gene set enrichment analyses in Gene Ontology of cellular component of differential gene expression (DEGs) results from 2-day dexamethasone exposure to Schlemm’s canal endothelial (SCE) cells. Significance was determined at adjusted P < 0.05. Please see the tab titled “SuppTable24_SC2Days_GO_CC” in the Excel file titled: Supplementary Excel Files.

**Supplementary Table 25.** All significant results for gene set enrichment analyses in Gene Ontology of molecular function of differential gene expression (DEGs) results from 2-day dexamethasone exposure to Schlemm’s canal endothelial (SCE) cells. Significance was determined at adjusted P < 0.05. Please see the tab titled “SuppTable25_SC2Days_GO_MF” in the Excel file titled: Supplementary Excel Files.

**Supplementary Table 26.** Enriched biological pathways for the differentially expressed genes (DEGs) in Schlemm’s canal endothelial (SCE) cells after 2 days of exposure to dexamethasone. Significance was determined at adjusted P < 0.05. Top 10 statistically most significant enriched pathways from KEGG and REACTOME database are shown. Selective columns are shown.

| **Source** | **Term name** | **Term id** | **Adjusted p value** |
| --- | --- | --- | --- |
| REAC | Extracellular matrix organization | REAC:R-HSA-1474244 | 6.37E-07 |
| REAC | Interleukin-4 and Interleukin-13 signaling | REAC:R-HSA-6785807 | 4.46E-06 |
| REAC | Regulation of Insulin-like Growth Factor transport and uptake by Insulin-like Growth Factor Binding Proteins | REAC:R-HSA-381426 | 9.38E-05 |
| REAC | Signal Transduction | REAC:R-HSA-162582 | 0.0003119 |
| KEGG | Extracellular matrix-receptor interaction | KEGG:04512 | 9.24E-05 |
| KEGG | Pathways in cancer | KEGG:05200 | 0.000505 |
| KEGG | Proteoglycans in cancer | KEGG:05205 | 0.000505 |
| KEGG | PI3K-Akt signaling pathway | KEGG:04151 | 0.00114997 |
| KEGG | Hypertrophic cardiomyopathy | KEGG:05410 | 0.00260922 |

**Supplementary Table 27.** Enriched biological pathways for the differentially expressed genes (DEGs) in Schlemm’s canal endothelial (SCE) cells after 2 days of exposure to dexamethasone. Significance was determined at adjusted P < 0.05. Please see the tab titled “SuppTable27_SC2Days_Pathways” in the Excel file titled: Supplementary Excel Files for the complete table.

**Supplementary Table 28.** Enriched hallmark gene sets for the differentially expressed genes (DEGs) in Schlemm’s canal endothelial (SCE) cells after 2 days of exposure to dexamethasone. Significance was determined at adjusted P < 0.05. Top 10 statistically most significant enriched gene sets are shown. Selective columns are shown.

| **Term** | **Description** | **LogP** | **Log** **(q-value)** |
| --- | --- | --- | --- |
| M5930 | HALLMARK EPITHELIAL MESENCHYMAL TRANSITION | -33.2393 | -31.540 |
| M5890 | HALLMARK TNFA SIGNALING VIA NFKB | -24.7444 | -23.346 |
| M5891 | HALLMARK HYPOXIA | -19.6434 | -18.422 |
| M5906 | HALLMARK ESTROGEN RESPONSE EARLY | -16.5492 | -15.452 |
| M5942 | HALLMARK UV RESPONSE DN | -11.9366 | -10.937 |
| M5902 | HALLMARK APOPTOSIS | -9.73329 | -8.812 |
| M5947 | HALLMARK IL2 STAT5 SIGNALING | -9.23823 | -8.384 |
| M5953 | HALLMARK KRAS SIGNALING UP | -9.15716 | -8.362 |
| M5939 | HALLMARK P53 PATHWAY | -9.10707 | -8.362 |
| M5924 | HALLMARK MTORC1 SIGNALING | -8.09893 | -7.400 |

**Supplementary Table 29.** Enriched biological pathways for the differentially expressed genes (DEGs) in Schlemm’s canal endothelial (SCE) cells after 2 days of exposure to dexamethasone. Significance was determined at adjusted P < 0.05. Please see the tab titled “SuppTable29_SC2Days_Hallmark” in the Excel file titled: Supplementary Excel Files for the complete table.

**Supplementary Table 30.** Enriched transcription factor binding sites for the differentially expressed genes (DEGs) in Schlemm’s canal endothelial (SCE) cells after 2 days of exposure to dexamethasone. Significance was determined at adjusted P < 0.05. Top 10 statistically significant enrichment transcription factors (TF) are shown. Selective columns are shown.

| **Source** | **Term name** | **Term id** | **Adjusted p value** |
| --- | --- | --- | --- |
| TF | Factor: MAZ; motif: GGGGGAGGGGGNGRGRRRGNRG | TF:M09984 | 3.52E-07 |
| TF | Factor: WT1; motif: RGGNGGGGGAGGRGGNGGRG | TF:M10108 | 7.83E-07 |
| TF | Factor: GKLF; motif: NNNRGGNGNGGSN | TF:M07289 | 7.83E-07 |
| TF | Factor: MAZ; motif: GGGGGAGGGGGNGRGRRRGNRG | TF:M09984 | 7.83E-07 |
| TF | Factor: MAZ; motif: GGGMGGGGS | TF:M10432 | 2.9167E-06 |
| TF | Factor: DB1; motif: GGRRRRGRRGGAGGGGGNGRRR | TF:M10107 | 4.0555E-06 |
| TF | Factor: Miz-1; motif: NNRGGWGGGGGAGGGGMRR | TF:M10112 | 5.4733E-06 |
| TF | Factor: GKLF; motif: NNRRGRRNGNSNNN | TF:M07040 | 1.5033E-05 |
| TF | Factor: GKLF; motif: NNRRGRRNGNSNNN | TF:M07040 | 1.765E-05 |

**Supplementary Table 31.** Enriched transcription factor binding sites for the differentially expressed genes (DEGs) in Schlemm’s canal endothelial (SCE) cells after 2 days of exposure to dexamethasone. Significance was determined at adjusted P < 0.05. Please see the tab titled “SuppTable31_SC2Days_TFBS” in the Excel file titled: Supplementary Excel Files for the complete table.

**Supplementary Table 32.** Enriched human phenotype ontology for the differentially expressed genes (DEGs) in Schlemm’s canal endothelial (SCE) cells after 2 days of exposure to dexamethasone. Significance was determined at adjusted P < 0.05. Only one statistically significant enrichment was observed. Selective columns are shown.

| **Source** | **Term name** | **Term id** | **Adjusted p value** |
| --- | --- | --- | --- |
| HP | Ischemic stroke | HP:0002140 | 0.01189018 |
| HP | Abnormal placenta morphology | HP:0100767 | 0.01903043 |
| HP | Autosomal dominant inheritance | HP:0000006 | 0.02967698 |
| HP | Aortic dissection | HP:0002647 | 0.02967698 |
| HP | Peripheral arterial stenosis | HP:0004950 | 0.02967698 |
| HP | Ischemic stroke | HP:0002140 | 0.01189018 |

**Supplementary Table 33.** Enriched human phenotype ontology for the differentially expressed genes (DEGs) in Schlemm’s canal endothelial (SCE) cells after 2 days of exposure to dexamethasone. Significance was determined at adjusted P < 0.05. Please see the tab titled “SuppTable33_SC2Days_HPO” in the Excel file titled: Supplementary Excel Files for the complete table.

**Supplementary Table 34.** Enriched biological pathways for the differentially expressed genes (DEGs) common in trabecular meshwork (TM) and Schlemm’s canal endothelial (SCE) cells after 2 days of exposure to dexamethasone. Significance was determined at adjusted P < 0.05. All statistically most significant enriched pathways from KEGG and REACTOME are shown. Selective columns are shown.

| **Source** | **Term name** | **Term id** | **Adjusted p value** |
| --- | --- | --- | --- |
| KEGG | Basal cell carcinoma | KEGG:05217 | 0.01625848 |
| REAC | Extracellular matrix organization | REAC:R-HSA-1474244 | 0.02070554 |
| REAC | Interleukin-4 and Interleukin-13 signaling | REAC:R-HSA-6785807 | 0.03239383 |

**Supplementary Table 35.** Enriched biological pathways for the differentially expressed genes (DEGs) common in trabecular meshwork (TM) and Schlemm’s canal endothelial (SCE) cells after 2 days of exposure to dexamethasone. Significance was determined at adjusted P < 0.05. Please see the tab titled “SuppTable35_TM-SC-2DaysPathways” in the Excel file titled: Supplementary Excel Files for the complete table.

**Supplementary Table 36.** All significant results of Gene Ontology gene set enrichment analyses for biological processes for 411 differentially expressed genes (DEGs) common to trabecular meshwork (TM) and Schlemm’s canal endothelial (SCE) cells after 2 days of dexamethasone exposure. Significance was determined at adjusted P < 0.05. Please see the tab titled “SuppTable36_TM-SC-2Days_GO_BP” in the Excel file titled: Supplementary Excel Files.

**Supplementary Table 37.** All significant results of Gene Ontology gene set enrichment analyses for cellular component for 411 differentially expressed genes (DEGs) common to trabecular meshwork (TM) and Schlemm’s canal endothelial (SCE) cells after 2 days of dexamethasone exposure. Significance was determined at adjusted P < 0.05. Please see the tabs titled “SuppTable37_TM-SC-2Days_GO_CC” in the Excel file titled: Supplementary Excel Files.

**Supplementary Table 38.** All significant results of Gene Ontology gene set enrichment analyses for molecular function for 411 differentially expressed genes (DEGs) common to trabecular meshwork (TM) and Schlemm’s canal endothelial (SCE) cells after 2 days of dexamethasone exposure. Significance was determined at adjusted P < 0.05. Please see the tabs titled “SuppTable38_TM-SC-2Days_GO_MF” in the Excel file titled: Supplementary Excel Files.

**Supplemental Table 39.** Selective quality control metrics for RNA sequencing of the trabecular meshwork (TM) and Schlemm’s canal endothelial cell (SCE) strains used in the 6-hour exposure experiment. All quality metrics were obtained from RNA sequencing. The sequencing was done for 151 cycles.

| **Sample ID** | **Mapping Rate** | **High Quality Rate** | **High Quality Reads** | **Mapped Reads** |
| --- | --- | --- | --- | --- |
| TM92_DEX | 0.989868 | 0.931436 | 73,179,700 | 78,566,500 |
| TM92_CTRL | 0.988145 | 0.921995 | 61,667,000 | 66,884,300 |
| TM96_DEX | 0.98997 | 0.934843 | 97,902,900 | 104,727,000 |
| TM96_CTRL | 0.989353 | 0.936415 | 88,248,700 | 94,241,100 |
| TM135_DEX | 0.988305 | 0.919565 | 86,286,500 | 93,834,000 |
| TM135_CTRL | 0.990212 | 0.935459 | 74,468,600 | 79,606,500 |
| TM151_DEX | 0.986633 | 0.919932 | 81,215,500 | 88,284,200 |
| TM151_CTRL | 0.986774 | 0.920866 | 62,673,700 | 68,059,500 |
| SC67_DEX | 0.986738 | 0.940881 | 110,693,000 | 117,648,000 |
| SC67_CTRL | 0.988036 | 0.93903 | 135,423,000 | 144,216,000 |
| SC77_DEX | 0.986521 | 0.933164 | 120,399,000 | 129,023,000 |
| SC77_CTRL | 0.99234 | 0.942734 | 128,809,000 | 136,633,000 |
| SC89_DEX | 0.988736 | 0.937027 | 106,757,000 | 113,931,000 |
| SC89_CTRL | 0.991333 | 0.944151 | 104,509,000 | 110,691,000 |
| SC91_DEX | 0.985813 | 0.934479 | 117,157,000 | 125,371,000 |
| SC91_CTRL | 0.990986 | 0.941642 | 134,636,000 | 142,980,000 |

Mapping Rate: The proportion of all reads in the BAM which were mapped, and not secondary alignments or platform/vendor quality control (QC) failing reads ("Mapped Reads").

High Quality Rate: The proportion of properly paired reads with less than 6 mismatched bases and a perfect mapping quality out of all "mapped reads"

High Quality Reads: Mapped reads that passed the following criteria: aligned as proper pairs, mismatches (’NM’ tag) at or below threshold, passed mapping quality (MAPQ) threshold

Mapped Reads: Unique mapping, vendor QC passed reads that were mapped

RNA=ribonucleic acid

**Supplementary Table 40.** High confidence results for 248 differentially expressed genes (DEGs) in trabecular meshwork (TM) cells after 6 hours of exposure to dexamethasone. Please see the tab titled “SuppTable40_TM6Hrs_HighConfDEGs” in the Excel file titled: Supplementary Excel Files for the complete table.

**Supplementary Table 41.** Full results of differential gene expression analysis (n=17,132) in trabecular meshwork (TM) cells after 6 hours of exposure to dexamethasone. Please see the tab titled “SuppTable41_TM6Hrs_All_DEGs” in the Excel file titled: Supplementary Excel Files for the complete table.

**Supplementary Table 42.** All significant Gene Ontology results from gene set enrichment analyses for biological processes of differentially expressed genes (DEGs) in trabecular meshwork (TM) cells after exposure to dexamethasone for 6 hours. Significance was determined at adjusted P < 0.05. Please see the tab titled “SuppTable42_TM6Hrs_GO_BP” in the Excel file titled: Supplementary Excel Files.

**Supplementary Table 43.** All significant Gene Ontology results from gene set enrichment analyses for cellular components of differentially expressed genes (DEGs) in trabecular meshwork (TM) cells after exposure to dexamethasone for 6 hours. Significance was determined at adjusted P < 0.05. Please see the tab titled “SuppTable43_TM6Hrs_GO_CC” in the Excel file titled: Supplementary Excel Files.

**Supplementary Table 44.** All significant Gene Ontology results from gene set enrichment analyses for molecular function of differentially expressed genes (DEGs) in trabecular meshwork (TM) cells after exposure to dexamethasone for 6 hours. Significance was determined at adjusted P < 0.05. Please see the tab titled “SuppTable44_TM6Hrs_GO_MF” in the Excel file titled: Supplementary Excel Files.

**Supplementary Table 45**. Enriched biological pathways for the differentially expressed genes (DEGs) in trabecular meshwork cells (TM) after 6-hours of exposure to dexamethasone. Significance was determined at adjusted P < 0.05. Top 10 statistically most significant enriched pathways from REACTOME and KEGG database are shown. Selective columns are shown.

| **Source** | **Term name** | **Term id** | **Adjusted p value** |
| --- | --- | --- | --- |
| REAC | Signaling by Interleukins | REAC:R-HSA-449147 | 8.2034E-09 |
| REAC | Cytokine Signaling in Immune system | REAC:R-HSA-1280215 | 8.2034E-09 |
| REAC | Signal Transduction | REAC:R-HSA-162582 | 1.158E-07 |
| REAC | Interleukin-4 and Interleukin-13 signaling | REAC:R-HSA-6785807 | 1.3241E-06 |
| REAC | Signaling by Nuclear Receptors | REAC:R-HSA-9006931 | 1.2055E-05 |
| KEGG | AGE-RAGE signaling pathway in diabetic complications | KEGG:04933 | 1.4964E-08 |
| KEGG | Cytokine-cytokine receptor interaction | KEGG:04060 | 7.4629E-08 |
| KEGG | TNF signaling pathway | KEGG:04668 | 1.8098E-06 |
| KEGG | Pathways in cancer | KEGG:05200 | 2.7326E-06 |
| KEGG | MAPK signaling pathway | KEGG:04010 | 2.7326E-06 |

**Supplementary Table 46**. Enriched biological pathways for the differentially expressed genes (DEGs) in trabecular meshwork cells (TM) after 6-hours of exposure to dexamethasone. Significance was determined at adjusted P < 0.05. Please see the tab titled “SuppTable46_TM6Hrs_Pathways” in the Excel file titled: Supplementary Excel Files for the complete table.

**Supplementary Table 47**. Enriched hallmark gene sets for the differentially expressed genes (DEGs) in trabecular meshwork (TM) cells after 6-hours of exposure to dexamethasone. Significance was determined at adjusted P < 0.05. Top 10 statistically most significant enriched gene sets are shown. Selective columns are shown.

| **Term** | **Description** | **Log P** | **Log (q-value)** |
| --- | --- | --- | --- |
| M5890 | HALLMARK TNFA SIGNALING VIA NFKB | -27.105197 | -25.41 |
| M5891 | HALLMARK HYPOXIA | -8.7995399 | -7.40 |
| M5902 | HALLMARK APOPTOSIS | -7.5956905 | -6.37 |
| M5932 | HALLMARK INFLAMMATORY RESPONSE | -7.4344701 | -6.34 |
| M5947 | HALLMARK IL2 STAT5 SIGNALING | -6.9097805 | -5.91 |
| M5930 | HALLMARK EPITHELIAL MESENCHYMAL TRANSITION | -6.5145038 | -5.59 |
| M5897 | HALLMARK IL6 JAK STAT3 SIGNALING | -5.8729943 | -5.02 |
| M5939 | HALLMARK P53 PATHWAY | -4.7110943 | -3.97 |
| M5906 | HALLMARK ESTROGEN RESPONSE EARLY | -4.3094051 | -3.65 |
| M5942 | HALLMARK UV RESPONSE DN | -3.7254624 | -3.11 |

**Supplementary Table 48**. Enriched hallmark gene sets for the differentially expressed genes (DEGs) in trabecular meshwork (TM) cells after 6-hours of exposure to dexamethasone. Significance was determined at adjusted P < 0.05. Please see the tab titled “SuppTable48_TM6Hrs_HallMark” in the Excel file titled: Supplementary Excel Files for the complete table.

**Supplementary Table 49.** Enriched human phenotype ontology for the differentially expressed genes (DEGs) in trabecular meshwork (TM) cells after 6 hours of exposure to dexamethasone. Statistically significant enrichments are shown. Selective columns are shown.

| **Source** | **Term name** | **Term id** | **Adjusted p value** |
| --- | --- | --- | --- |
| HP | Supraventricular tachycardia | HP:0004755 | 0.01637351 |
| HP | Paroxysmal supraventricular tachycardia | HP:0004763 | 0.01637351 |
| HP | Supraventricular arrhythmia | HP:0005115 | 0.01637351 |

**Supplementary Table 50.** Enriched human phenotype ontology for the differentially expressed genes (DEGs) in trabecular meshwork (TM) cells after 6 hours of exposure to dexamethasone. Significance was determined at adjusted P < 0.05. Please see the tab titled “SuppTable50_TM6Hrs_HPO” in the Excel file titled: Supplementary Excel Files for the complete table.

**Supplementary Table 51.** Enriched transcription factor binding sites for the differentially expressed genes (DEGs) in trabecular meshwork (TM) cells after 6-hours of exposure to dexamethasone. Significance was determined at adjusted P < 0.05. Top 10 statistically significant enrichment transcription factors (TF) are shown. Selective columns are shown.

| **Source** | **Term name** | **Term id** | **Adjusted p value** |
| --- | --- | --- | --- |
| TF | Factor: Spi-B; motif: TTCYBC | TF:M03851 | 3.3091E-18 |
| TF | Factor: Kid3; motif: CCACN | TF:M01160 | 1.849E-16 |
| TF | Factor: CPBP; motif: SNCCCNN | TF:M01822 | 1.7197E-13 |
| TF | Factor: Kaiso; motif: GCMGGGRGCRGS | TF:M03876 | 1.163E-09 |
| TF | Factor: ETF; motif: GVGGMGG | TF:M00695 | 7.2826E-09 |
| TF | Factor: ETF; motif: CCCCGCCCCYN | TF:M07039 | 1.5339E-08 |
| TF | Factor: E2F-2; motif: GCGCGCGCGYW | TF:M11531 | 2.4926E-08 |
| TF | Factor: Pax-5; motif: RRNGRNGCAN | TF:M03577 | 2.651E-08 |
| TF | Factor: MAZ; motif: GGGMGGGGSSGGGGGGGGGGGG | TF:M09636 | 3.2203E-08 |
| TF | Factor: TIEG1; motif: NCCCNSNCCCCGCCCCC | TF:M12351 | 3.5352E-08 |

**Supplementary Table 52.** Enriched transcription factor binding sites for the differentially expressed genes (DEGs) in trabecular meshwork (TM) cells after 6-hours of exposure to dexamethasone. Significance was determined at adjusted P < 0.05. Please see the tab titled “SuppTable52_TM6Hrs_TFBS” in the Excel file titled: Supplementary Excel Files for the complete table.

**Supplemental Table 53.** Selective quality control metrics for RNA sequencing of the trabecular meshwork (TM) and Schlemm’s canal (SC) endothelial cell strains used in the 1-hour exposure experiment. All the quality metrics obtained from RNA sequencing. The sequencing was done for 151 cycles.

| **Sample ID** | **Mapping Rate** | **High Quality Rate** | **High Quality Reads** | **Mapped Reads** |
| --- | --- | --- | --- | --- |
| TM96_CTRL | 0.993341 | 0.944487 | 78,216,300 | 82,813,500 |
| TM96_DEX | 0.992836 | 0.944916 | 83,812,500 | 88,698,400 |
| TM123_CTRL | 0.993571 | 0.94747 | 99,532,800 | 105,051,000 |
| TM123-DEX | 0.993461 | 0.947486 | 90,228,400 | 95,229,300 |
| TM151_CTRL | 0.993737 | 0.945982 | 102,764,000 | 108,632,000 |
| TM151_DEX | 0.991409 | 0.926953 | 80,657,700 | 87,013,700 |
| SC86_CTRL | 0.992211 | 0.943079 | 94,506,200 | 100,210,000 |
| SC86_DEX | 0.99286 | 0.945232 | 74,502,900 | 78,819,700 |
| SC89_CTRL | 0.993168 | 0.937114 | 51,184,200 | 54,618,900 |
| SC89_DEX | 0.992031 | 0.940211 | 88,946,900 | 94,603,100 |
| SC91_CTRL | 0.993885 | 0.93997 | 84,512,000 | 89,909,200 |
| SC91_DEX | 0.993256 | 0.942401 | 49,748,600. | 52,789,300 |

Mapping Rate: The proportion of all reads in the BAM which were mapped, and not secondary alignments or platform/vendor quality control (QC) failing reads ("Mapped Reads").

High Quality Rate: The proportion of properly paired reads with less than 6 mismatched bases and a perfect mapping quality out of all "Mapped Reads"

High Quality Reads: Mapped Reads that passed the following criteria: aligned as proper pairs, mismatches (’NM’ tag) at or below threshold, passed mapping quality (MAPQ) threshold

Mapped Reads: Unique mapping, vendor QC passed Reads that were mapped

RNA=ribonucleic acid, DEX=dexamethasone-treated cells, CTRL=vehicle-treated cells (controls

**Supplementary Table 54.** High confidence results for differentially expressed genes (DEGs) after 6-hour exposure to dexamethasone in Schlemm’s Canal Endothelial (SCE) cells. Significance was determined at adjusted P < 0.05. Please see the tab titled “SuppTable54_SCE6Hrs_HighConf_DEGs" in the Excel file titled: Supplementary Excel Files for the complete table.

**Supplementary Table 55.** Full results of differential gene expression analysis (n=17,327) after 6-hour exposure to dexamethasone in Schlemm’s Canal Endothelial (SCE) cells. Please see the tab titled “SuppTable55_SCE6Hrs_AllDEGs" in the Excel file titled: Supplementary Excel Files for the complete table.

**Supplementary Table 56.** All significant Gene Ontology results from gene set enrichment analyses for biological processes of differentially expressed genes (DEGs) in SCE after exposure to dexamethasone for 6 hours. Significance was determined at adjusted P < 0.05. Please see the tab titled “SuppTable56_SCE6Hrs_GO_BP” in the Excel file titled: Supplementary Excel Files.

**Supplementary Table 57.** All significant Gene Ontology results from gene set enrichment analyses for biological processes of differentially expressed genes (DEGs) in SCE after exposure to dexamethasone for 6 hours. Significance was determined at adjusted P < 0.05. Please see the tab titled “SuppTable56_SCE6Hrs_GO_MF” in the Excel file titled: Supplementary Excel Files.

**Supplementary Table 58.** All significant Gene Ontology results from gene set enrichment analyses for biological processes of differentially expressed genes (DEGs) in SCE after exposure to dexamethasone for 6 hours. Significance was determined at adjusted P < 0.05. Please see the tab titled “SuppTable56_SCE6Hrs_GO_CC” in the Excel file titled: Supplementary Excel Files.

**Supplementary Table 59.** Enriched biological pathways for differentially expressed genes (DEGs) in SCE after exposure to dexamethasone for 6 hours. Significance was determined at adjusted P < 0.05. Please see the tab titled “SuppTable59_SCE6Hrs_Pathways” in the Excel file titled: Supplementary Excel Files.

**Supplementary Table 60**. Enriched hallmark gene sets for the differentially expressed genes (DEGs) in SCE cells after 6-hours of exposure to dexamethasone. Significance was determined at adjusted P < 0.05. Please see the tab titled “SuppTable60_SC6Hrs_HallMark” in the Excel file titled: Supplementary Excel Files for the complete table.

**Supplementary Table 61.** Enriched transcription factor binding sites for the differentially expressed genes (DEGs) in SCE cells after 6-hours of exposure to dexamethasone. Significance was determined at adjusted P < 0.05. Please see the tab titled “SuppTable61_SC6Hrs_TFBS” in the Excel file titled: Supplementary Excel Files for the complete table.

**Supplementary Table 62 .** High confidence results for common significant differentially expressed genes (DEGs) after 1 hour exposure to dexamethasone in TM and SCE cells. Significance was determined at adjusted P < 0.05. Please see the tab titled “SuppTable62_Comm_TM_SC_6Hr " in the Excel file titled: Supplementary Excel Files for the complete table.

**Supplementary Table 63.** Significant differentially expressed genes (DEGs) after 1 hour exposure to dexamethasone in trabecular meshwork (TM) cells. Significance was determined at adjusted P < 0.05. Please see the tab titled “SuppTable63_TM1Hr_HighConf_DEGs” in the Excel file titled: Supplementary Excel Files for the complete table.

**Supplementary Table 64.** All significant results for differentially expressed genes (DEGs) after 1 hour exposure to dexamethasone in trabecular meshwork (TM) cells. Significance was determined at adjusted P < 0.05. Please see the tab titled “SuppTable64_TM1Hr_All_DEGs” in the Excel file titled: Supplementary Excel Files for the complete table.

**Supplementary Table 65.** High confidence results for differentially expressed genes (DEGs) after 1 hour exposure to dexamethasone in Schlemm’s canal endothelial (SCE) cells. Significance was determined at adjusted P < 0.05. Please see the tab titled “SuppTable65_SC1Hr_HighConf_DEGs” in the Excel file titled: Supplementary Excel Files for the complete table.

**Supplementary Table 66.** Full results for differentially expressed genes (DEGs) after 1 hour exposure to dexamethasone in Schlemm’s canal endothelial (SCE) cells. Please see the tab titled “SuppTable66_SC1Hr_All_DEGs” in the Excel file titled: Supplementary Excel Files for the complete table.

**Supplementary Table 67.** Common differentially expressed genes (DEGs) between trabecular meshwork (TM) and Schlemm’s canal endothelial (SCE) cell lines with 1 hour of dexamethasone exposure. Significance was determined at adjusted P < 0.05. Please see the tab titled “SuppTable67_Common_TM_SC_1Hr” in the Excel file titled: Supplementary Excel Files

**Supplementary Table 68.** Gene set enrichment analysis Gene Ontology results for biological processes of differentially expressed genes (DEGs) from trabecular meshwork (TM) cells after exposure to dexamethasone for one hour. Significance was determined at adjusted P < 0.05. Please see the tab titled “SuppTable68_TM1Hr_GSEA_GO_BP” in the Excel file titled: Supplementary Excel Files.

**Supplementary Table 69.** Gene set enrichment analysis Gene Ontology results for cellular components of differentially expressed genes (DEGs) from trabecular meshwork (TM) cells after exposure to dexamethasone for one hour. Significance was determined at adjusted P < 0.05. Please see the tab titled “SuppTable69_TM1Hr_GSEA_GO_CC” in the Excel file titled: Supplementary Excel Files.

**Supplementary Table 70.** Gene set enrichment analysis Gene Ontology results for molecular function of differentially expressed genes (DEGs) from trabecular meshwork (TM) cells after exposure to dexamethasone for one hour. Significance was determined at adjusted P < 0.05. Please see the tab titled “SuppTable70_TM1Hr_GSEA_GO_MF” in the Excel file titled: Supplementary Excel Files.

**Supplementary Table 71.** Enriched biological pathways for the differentially expressed genes (DEGs) in trabecular meshwork (TM) cells after 1-hour of exposure to dexamethasone. Significance was determined at adjusted P < 0.05. Top 10 statistically most significant enriched pathways from KEGG and REACTOME database are shown. Selective columns are shown.

| **Source** | **Term name** | **Term id** | **Adjusted p value** |
| --- | --- | --- | --- |
| KEGG | IL-17 signaling pathway | KEGG:04657 | 4.11E-07 |
| KEGG | TNF signaling pathway | KEGG:04668 | 2.1375E-06 |
| KEGG | Kaposi sarcoma-associated herpesvirus infection | KEGG:05167 | 0.00031888 |
| KEGG | Human T-cell leukemia virus 1 infection | KEGG:05166 | 0.00063155 |
| KEGG | Measles | KEGG:05162 | 0.00114698 |
| REAC | Interleukin-10 signaling | REAC:R-HSA-6783783 | 0.00090634 |
| REAC | Interleukin-4 and Interleukin-13 signaling | REAC:R-HSA-6785807 | 0.00090634 |
| REAC | Signal Transduction | REAC:R-HSA-162582 | 0.00090634 |
| REAC | Cytokine Signaling in Immune system | REAC:R-HSA-1280215 | 0.00156978 |

**Supplementary Table 72.** Enriched biological pathways for the differentially expressed genes (DEGs) in trabecular meshwork (TM) cells after 1-hour of exposure to dexamethasone. Significance was determined at adjusted P < 0.05. Please see the tab titled “SuppTable72_TM1Hr_GSEA_Pathways” in the Excel file titled: Supplementary Excel Files for the complete table.

**Supplementary Table 73.** Enriched transcription factor binding sites for the differentially expressed genes (DEGs) in trabecular meshwork (TM) cells after 1-hour of exposure to dexamethasone. Significance was determined at adjusted P < 0.05. Top 10 statistically significant enrichment transcription factor (TF) are shown. Selective columns are shown.

| **Source** | **Term name** | **Term id** | **Adjusted p value** |
| --- | --- | --- | --- |
| TF | Factor: SRF; motif: ATGCCCATATATGGWNNT | TF:M00152 | 0.004022 |
| TF | Factor: ATF; motif: CNSTGACGTNNNYC | TF:M00017 | 0.02189958 |
| TF | Factor: CREB,; motif: NTGACGTNA | TF:M00981 | 0.02189958 |
| TF | Factor: AP-2gamma; motif: GCCYNNGGS | TF:M00470 | 0.02292335 |
| TF | Factor: ATF-1; motif: TNACGTCAN | TF:M01861 | 0.02292335 |
| TF | Factor: CREB; motif: TGACGTMA | TF:M00039 | 0.02292335 |
| TF | Factor: ATF-1; motif: NNNTGACGTNNN | TF:M07034 | 0.02348811 |
| TF | Factor: SP2; motif: GGGCGGGAC | TF:M01783 | 0.02636228 |
| TF | Factor: ATF6; motif: TGACGTGG | TF:M00483 | 0.02636228 |

TF=transcription factor

**Supplementary Table 74.** Enriched transcription factor binding sites for the differentially expressed genes (DEGs) in trabecular meshwork (TM) cells after 1-hour of exposure to dexamethasone. Significance was determined at adjusted P < 0.05. Please see the tab titled “SuppTable74_TM1Hr_GSEA_TFBS” in the Excel file titled: Supplementary Excel Files for the complete table.

**Supplementary Table 75.** Gene set enrichment Gene Ontology results for biological processes of differentially expressed genes (DEGs) from Schlemm’s canal endothelial (SCE) cells after exposure to dexamethasone for one hour. Significance was determined at adjusted P < 0.05. Please see the tab titled “SuppTable75_SC1Hr_GSEA_GO_BP” in the Excel file titled: Supplementary Excel Files.

**Supplementary Table 76.** Gene set enrichment Gene Ontology results for cellular components of differentially expressed genes (DEGs) from Schlemm’s canal endothelial (SCE) cells after exposure to dexamethasone for one hour. Significance was determined at adjusted P < 0.05. Please see the tab titled “SuppTable76_SC1Hr_GSEA_GO_CC” in the Excel file titled: Supplementary Excel Files.

**Supplementary Table 77.** Gene set enrichment Gene Ontology results for molecular function of differentially expressed genes (DEGs) from Schlemm’s canal endothelial (SCE) cells after exposure to dexamethasone for one hour. Significance was determined at adjusted P < 0.05. Please see the tab titled “SuppTable77_SC1Hr_GSEA_GO_MF” in the Excel file titled: Supplementary Excel Files.

**Supplementary Table 78.** Enriched biological pathways for the differentially expressed genes (DEGs) in Schlemm’s canal cells (SCE) after 1-hour of exposure to dexamethasone. Significance was determined at adjusted P < 0.05. Top 10 statistically most significant enriched pathways from KEGG and REACTOME database are shown. Selective columns are shown.

| **Source** | **Term name** | **Term id** | **Adjusted p value** |
| --- | --- | --- | --- |
| REAC | NGF-stimulated transcription | REAC:R-HSA-9031628 | 1.6717E-05 |
| REAC | Metallothioneins bind metals | REAC:R-HSA-5661231 | 9.1579E-05 |
| REAC | Nuclear Events (kinase and transcription factor activation) | REAC:R-HSA-198725 | 9.1579E-05 |
| REAC | Response to metal ions | REAC:R-HSA-5660526 | 0.00016407 |
| REAC | Signaling by NTRK1 (TRKA) | REAC:R-HSA-187037 | 0.00085447 |
| REAC | Signaling by NTRKs | REAC:R-HSA-166520 | 0.00153047 |
| KEGG | TNF signaling pathway | KEGG:04668 | 0.00220034 |
| KEGG | IL-17 signaling pathway | KEGG:04657 | 0.00349101 |
| KEGG | KEGG root term | KEGG:00000 | 0.00366515 |
| KEGG | Osteoclast differentiation | KEGG:04380 | 0.00495675 |

**Supplementary Table 79.** Enriched biological pathways for the differentially expressed genes (DEGs) in Schlemm’s canal cells (SCE) after 1-hour of exposure to dexamethasone. Significance was determined at adjusted P < 0.05. Please see the tab titled “SuppTable79_SC1Hr_GSEA_Pathways” in the Excel file titled: Supplementary Excel Files for the complete table.

**Supplementary Table 80.** Enriched transcription factor binding sites for the differentially expressed genes (DEGs) in trabecular meshwork (TM) cells after 1-hour of exposure to dexamethasone. Significance was determined at adjusted P < 0.05. Top 10 statistically significant enrichment transcription factors (TF) are shown. Selective columns are shown.

| **Source** | **Term name** | **Term id** | **Adjusted p value** |
| --- | --- | --- | --- |
| TF | Factor: CPBP; motif: SNCCCNN | TF:M01822 | 0.00313872 |
| TF | Factor: NF-kappaB; motif: GGGGATYCCC | TF:M00051 | 0.00313872 |
| TF | Factor: E2F-3:TBR2; motif: ANGTGYKANGGCGCSTTNNCRNNT | TF:M08207 | 0.00376536 |
| TF | Factor: Kid3; motif: CCACN | TF:M01160 | 0.00376536 |
| TF | Factor: Spi-B; motif: TTCYBC | TF:M03851 | 0.00376536 |
| TF | Factor: ZNF614; motif: NCYCWGCCYYNNN | TF:M09862 | 0.00376536 |
| TF | Factor: AP-2gamma; motif: GCCYNNGGS | TF:M00470 | 0.00700604 |
| TF | Factor: MTF-1; motif: TBTGCACHCGGCCC | TF:M00650 | 0.01096083 |
| TF | Factor: AP-4; motif: RNCAGCTGC | TF:M00927 | 0.01096083 |
| TF | Factor: E2F-1:HES-7; motif: GGCRCGTGSYNNWNGGCGCSM | TF:M08525 | 0.01096083 |

TF=transcription factor

**Supplementary Table 81.** Enriched transcription factor binding sites for the differentially expressed genes (DEGs) in trabecular meshwork (TM) cells after 1-hour of exposure to dexamethasone. Significance was determined at adjusted P < 0.05. Please see the tab titled “SuppTable81_SC1Hr_GSEA_TFBS” in the Excel file titled: Supplementary Excel Files for the complete table.

**Supplementary Table 82.** Marker gene expression for Trabecular Meshwork cell lines. Please see the tab titled “SuppTable82_TM_MarkerExpression” in the Excel file titled: Supplementary Excel Files for the complete table.

**Supplementary Table 83.** Marker gene expression for Schlemm’s Canal Endothelium cell lines. Please see the tab titled “SuppTable83_SC_MarkerExpression” in the Excel file titled: Supplementary Excel Files for the complete table.

**Supplementary Table 84.** Comparative of quantitative polymerase chain reaction (qPCR) relative quantification (RQ) and RNA-seq fold-change (FC) for 2 days of dexamethasone exposure across nine trabecular meshwork (TM) cell strains. RQ is defined as the ratio of expression between the test and calibrated control samples from qPCR. RNA-seq FC was computed using DESeq2. Strong correlation between log_2_ qPCR RQ and RNA-Seq FC was observed (see Fig. 8; Pearson correlation r=0.94) among all the strains with one exception for *NR3C1* in one sample, TM137.

| **Gene Name** | **TM92** | **TM96** | **TM123** | **TM134** | **TM135** | **TM137** | **TM140** | **TM151** | **TM155** | **RNA-seq FC** | **RNA-seq log_2_(FC)** |
| --- | --- | --- | --- | --- | --- | --- | --- | --- | --- | --- | --- |
| *NR3C1* | 0.80 | 0.68 | 0.76 | 0.80 | 0.61 | 1.97 | 0.59 | 0.88 | 0.98 | 0.78 | -0.36 |
| *KLF9* | 2.03 | 3.58 | 2.29 | 2.03 | 3.87 | 4.90 | 2.18 | 1.43 | 1.92 | 1.89 | 0.92 |
| *PDK4* | 2.28 | 4.82 | 2.45 | 2.28 | 5.23 | 10.47 | 3.30 | 1.77 | 1.90 | 2.00 | 1.00 |
| *LTBP2* | 1.72 | 3.82 | 1.94 | 1.72 | 2.54 | 6.99 | 1.46 | 1.37 | 1.63 | 2.28 | 1.19 |
| *TNFRSF21* | 2.69 | 1.57 | 2.30 | 2.69 | 3.32 | 5.11 | 2.21 | 2.51 | 3.09 | 2.81 | 1.49 |
| *ITGA10* | 2.09 | 4.91 | 2.19 | 2.09 | 12.17 | 27.26 | 3.41 | 3.02 | 5.91 | 2.91 | 1.54 |
| *NEDD9* | 3.03 | 5.19 | 4.82 | 3.03 | 2.15 | 1.32 | 2.94 | 2.15 | 2.95 | 3.39 | 1.76 |
| *PER1* | 2.87 | 4.23 | 4.36 | 2.87 | 3.48 | 4.87 | 2.92 | 3.00 | 2.94 | 3.61 | 1.85 |
| *FAM107A* | 5.73 | 11.88 | 3.13 | 5.73 | 21.49 | 16.96 | 18.53 | 4.24 | 9.13 | 4.06 | 2.02 |
| *FKBP5* | 4.89 | 14.27 | 4.85 | 4.89 | 9.54 | 28.86 | 6.79 | 3.06 | 4.88 | 4.38 | 2.13 |
| *LINC01088* | 14.49 | 56.04 | 15.65 | 14.49 | 52.14 | 32.64 | 23.68 | 19.34 | 11.28 | 12.91 | 3.69 |

**Supplementary Table 85.** Comparative of quantitative polymerase chain (qPCR) reaction relative quantification (RQ) and RNA-seq fold-change (FC) for 2 days of dexamethasone exposure across four Schlemm’s canal endothelial (SCE) cell strains. RQ is defined as the ratio of expression between the test and calibrated control samples from qPCR. RNA-seq FC was computed using DESeq2. Strong correlation between log_2_ qPCR RQ and RNA-seq FC was observed (see Fig. 8; Pearson correlation r=0.98) among all SCE samples.

| **Gene Name** | **SC67** | **SC86** | **SC89** | **SC91** | **RNA-seq FC** | **RNA-seq log_2_(FC)** |
| --- | --- | --- | --- | --- | --- | --- |
| *COL15A1* | 0.10 | 0.25 | 0.18 | 0.13 | 0.13 | -3.00 |
| *NR3C1* | 0.64 | 0.71 | 0.69 | 0.61 | 0.64 | -0.64 |
| *TSC22D3* | 4.17 | 9.04 | 4.23 | 6.30 | 4.5 | 2.17 |
| *GLUL* | 3.68 | 3.47 | 6.73 | 5.91 | 4.89 | 2.29 |
| *ANGPTL1* | 2.62 | 3.01 | 2.50 | 2.41 | 5.74 | 2.52 |
| *APOD* | 4.81 | 8.60 | 4.65 | 10.31 | 6.54 | 2.71 |
| *SAMHD1* | 6.60 | 12.45 | 10.64 | 18.53 | 9.58 | 3.26 |
| *FKBP5* | 9.82 | 20.04 | 21.05 | 11.20 | 13.45 | 3.75 |
| *GPX3* | 14.23 | 24.27 | 14.45 | 62.00 | 17.27 | 4.11 |
| *FAM107A* | 26.59 | 38.73 | 43.37 | 31.90 | 25.11 | 4.65 |

**Supplementary Table 86.** TaqMan probes and the GenBank Accession numbers for each gene expression assay. Please see the tab titled “SuppTable86_TaqManProbes” in the Excel file titled: Supplementary Excel Files for the complete table.
