## Supplementary Materials for "Time-dependent Glucocorticoid-Induced Transcriptomic Changes in Human Trabecular Meshwork and Schlemm’s Canal"

**Supplementary Material**

**Supplementary Methods**

**Cell Culture**

Donors ranged between 28 to 77 years of age. Cell cultures were maintained in Dulbecco’s modified Eagle’s medium supplemented with 10% fetal bovine serum, 100ug/mL penicillin, 100ug/mL streptomycin, and 292 ug/mL L-glutamine at 37ºC in a 5% CO_2_ atmosphere. To differentiate the confluent cultures, TM cells were switched to medium containing 1% fetal bovine serum the week before treatments and remained in medium containing 1% fetal bovine serum during treatments. SCE cells were switched to a medium containing 10% adult bovine serum and remained in that during the drug treatment.

**RNA-Sequencing Methods**

Total RNA was quantified using the Quant-iT™ RiboGreen® RNA Assay Kit and normalized. Quality of extracted RNA was determined by RQS from LabChip GX Touch. An RQS above 6 was considered good quality RNA. External RNA Control Consortium RNA (ERCC) spike-in controls were added to each sample before sequencing, and a k562 control was used as well. The ERCC controls were used to control for variability, including level of cellularity and RNA yield. An aliquot for each sample was taken for library preparation, using an automated variation of the Illumina TruSeq™ Stranded mRNA Sample Preparation Kit. The resultant cDNA then went through dual-indexed library preparation which includes ‘A’ base addition, making it a poly(A) selection library. After enrichment the libraries were quantified using Quant-iT PicoGreen. The sample set was pooled and quantified using the KAPA Library Quantification Kit for Illumina Sequencing Platforms. Strand-specific, 101 bp (for the 2-day time point) or 151 bp (for the 1-hour and 6-hour time points) paired-end RNA-seq was performed on Illumina’s NovaSeq 6000 platform with a minimum of 50 million paired-end reads, as per manufacturer’s protocols.

**Quality control (QC) of trabecular meshwork (TM) and Schlemm’s canal endothelial** **(SCE) RNA-seq samples**

Post alignment, quality control (QC) was performed using RNA-SeQC 2^1^ with particular emphasis on the following metrics: mapping rate, high quality rate, high quality reads, unique mapped reads, read length, mean 3’ and 5’ bias, and read end 1 and end 2 sense strand information. For more information on these parameters please refer to Graubert et al.^1^. Additionally, raw read counts were transformed to normalized TPM values for each gene in every sample using RNA-SeQC 2. RNA-seq samples that passed extensive quality control including high RNA quality, sequencing depth of over 50 million reads, and read mapping rate above 0.99 were used in downstream analyses (**Supplementary Tables 1-2, Supplementary Figures 1-2, 8-11**).

*QC of 2-day GC Exposure RNA-seq samples*

Of the ten TM cell strains used in the 2-day GC exposure experiments, nine strains reached the minimum sequencing read depth of 50 million reads. The strain that did not reach this threshold (TM141) was excluded from all analysis. All five SCE cell strains met the minimum read count. All dexamethasone (DEX)-treated and control TM and SCE samples had high quality RNA with a minimum relative quantification score (RQS) of 8 and average RQS ranging between 8.10 and 9.42 (**Supplementary Table 1**). Selective QC metrics for RNA-seq results of the nine TM and five SCE paired samples are shown in **Supplementary Table 2**. All samples displayed high mapping rate (>0.99) and high sequencing coverage (average high-quality reads of 86.7 million). We additionally inspected the distribution of gene expression for the TM and SCE samples using transcript per million (TPM) values (**Supplementary Figure 1) to identify any outliers**.

For the TM cell strains, similar distributions were observed across the samples with only minor differences in the mean values. The TM92 sample pair showed slightly lower gene expression values compared to the other samples, but they still clustered as expected within the clades (**Supplementary Figure 2A**), so they were included in the analysis. For the SCE cell strains, similar distributions were observed across all samples except for the DEX-treated SC71 sample, which had a significantly lower mean that the other samples (**Supplementary Figure 2B**). The sample underwent re-sequencing but remained an outlier based on its gene expression profile, even after cross-sample normalization, so this sample was removed from further analyses. Therefore, nine TM cells and four SCE samples were used in subsequent analyses.

*QC of 6-hour GC Exposure RNA-seq samples*

Four TM and SCE cell strains from the 2-day exposure experiments were exposed to DEX and no treatment (control) for 6 hours, followed by RNA-sequencing. Quality control metrics for the RNA-seq results of these four TM and SCE paired samples are shown in **Supplementary Table 39**. TPM distributions for the TM and SCE paired samples are shown in **Supplementary Figure 8A and 8B**. Comparable distributions were observed across the samples with only minor differences in the mean values. The hierarchical clustering of gene expression profiles across the samples is shown in **Supplementary Figure 9A and 9B**. Samples from the same donors clustered within the same clade based on their gene expression profiles, as expected.

*QC of 1-hour GC Exposure RNA-seq samples*

Three TM and three SCE cell strains from the 2-day exposure experiments were exposed to DEX for 1 hour, followed by RNA-sequencing. TPM distributions for the TM and SCE paired samples are shown in **Supplementary Figure 10**. For both the TM and SCE cell strains, a comparable distribution was seen across the samples with only minor differences in the mean values. The hierarchical clustering of gene expression profiles across the samples is shown in **Supplementary Figure 11A and 11B**. Samples from the same donors clustered within the same clade based on their gene expression profiles, as expected.

*Gene Expression Analysis Methods*

Gene expression quantification was computed using featureCount from the SubRead package with the following non-default settings: reads must be paired, both read pairs must be mapped, only uniquely mapped reads were used, multi-mapped reads were not counted, chimeric reads were not counted, and strand specificity was turned on. To check for possible sample mismatch, the DESeq2^2^ normalized read counts of all samples were hierarchically clustered using Euclidean distance between the expression profiles, and Spearman’s rank correlation was computed between all pairwise comparisons.

**qPCR validation of RNA-seq differential gene expression**

To validate the RNA-seq results, we performed quantitative qPCR on a subset of the significantly differentially expressed genes. Genes were chosen based on (1) being among the topmost highly differentiated genes in the experiments (lowest adjusted P value), (2) being significantly differentiated at multiple DEX exposure time points, (3) being also previously associated with POAG and/or IOP in previous genome-wide association study (GWAS) meta-analyses for these traits, and (4) being enriched for in gene-set enrichment analysis (GSEA). In total, 11 genes from TM and 10 genes from SCE expression patterns were validated by quantitative PCR. The delta CT (threshold cycle) method was used for relative quantification (RQ), defined as the ratio of expression between the test and calibrated control samples. The RQ values of greater than 1 refer to up-regulation and less than 1 down-regulation. The Pearson correlation coefficient, *r* was used to assess the association of gene expression changes between RNA-Seq and qPCR RQ values.

Isolated total RNA (RIN>7.0) was DNase-treated with ezDNase (Life Technologies Corporation, Carlsbad, CA) to remove any genomic DNA contaminants and quantified. Reverse transcription (RT) was performed using SuperScript IV Vilo (Life Technologies Corporation, Carlsbad, CA) with equal mass amounts of RNA for each case and control sample containing a minimum of 500ng according to the manufacturer’s recommendations for oligo(dT)20 primed cDNA-synthesis, along with a negative control. Candidate normalizer genes, *GAPDH, PPIA and UBC,* were chosen based on their constant expression levels in RNA-seq studies. *GAPDH* was selected as the normalizer gene after performing qPCR on 1:10, 1:20 and 1:100 dilutions of an equal volume of each case and control cDNA and comparing threshold cycle (CT) values directly between case and control samples. Finally, cDNA was diluted to 1:20 prior to use in qPCR.

TaqMan probes were selected for the target genes to bind specifically to human cDNA and all gene expression assays have been validated to have >95% PCR efficiency (Life Technologies Corporation, Carlsbad, CA). TaqMan probes were added to the TaqMan Fast Advanced Master Mix containing reporter fluorescent dye FAM (6-carboxy-fluorescein) and carry the quencher dye TAMRA (6-carboxy-tetramethyl-rhodamine, with passive reference dye ROX and 2uL of diluted cDNA was added to the mix. PCR was performed in a QuantStudio 3Flex Real‑Time PCR System in 96–well fast microamp plates using a final volume of 20 uL. Amplifications were performed starting with a 20 second polymerase activation step at 95C, followed by 40 cycles of denaturation at 95C for 1 second and combined primer annealing/extension at 60C for 20 seconds. Fluorescence increase of FAM was measured during PCR. All case and control samples were amplified in triplicate. Ct values of <40 were excluded from further mathematical calculations. The CT is defined as the number of cycles needed for the fluorescence to reach a specific threshold level of detection and is inversely correlated with the amount of template nucleic acid present in the reaction (Walker et al, 2002).

*Gene set enrichment analysis (GSEA) of DEX-responsive genes – Hallmark gene sets*

Metascape [v3.5] was used for GSEA of Hallmark gene sets with the exception of the 6-hour TM analysis, where g:Profiler was used for enrichment using the same Hallmark gene sets that was used by Metascape.

*Mapping of DEX-responsive genes to IOP and glaucoma GWAS loci*

The identified high confidence DEGs were examined for association with primary open angle glaucoma (POAG) and IOP genetic loci taken from the largest available GWAS meta-analyses of these traits, including the UK Biobank GWAS. We inspected 133 independent variants in 112 loci reported for IOP in a GWAS meta-analysis of 139,555 primarily European individuals ^5^, 127 POAG loci identified in a cross-ancestry meta-analysis of 34,179 cases and 349,321 controls ^6^, 68 (partially overlapping) POAG loci from a European subset of 16,677 POAG cases and 199,580 controls ^6^, 312 loci from a cross-ancestry multi-trait analysis of GWAS of POAG, IOP and vertical cup-to-disc ratio (n=697,345 participants) and 263 loci from the European subset of the POAG MTAG (n=644,750 participants) ^7^. Since the majority of genetic associations with POAG and IOP lie in noncoding regions, we assumed that the underlying causal mechanisms of GWAS loci are regulatory, and thus, that target genes of expression quantitative trait loci (eQTLs) or splicing QTLs (sQTLs) that colocalize (i.e., share a causal variant) with a GWAS locus may be the implicated causal gene. To determine if any of the DEX DEGs may be associated with POAG risk or IOP variation, we used a previous mapping that we conducted of putative disease-causing genes to POAG and IOP loci based on colocalization analysis of eQTLs and sQTLs from 49 GTEx tissues ^8^, many of which glaucoma relevant cell types (e.g., artery, fibroblasts, neurons), and retina tissue^9^. with each of the GWAS loci ^10^. Colocalization analysis was performed by applying two Bayesian methods, eCAVIAR ^11^ and enloc^12^ to all eQTLs and sQTLs, which overlapped each of the IOP or POAG loci stated above. Each DEG from TM and SCE for all the time points was examined for being a target gene of an e/sQTL that colocalized with a GWAS locus. Since the e/sQTLs did not capture a causal mechanism for all POAG and IOP loci, we also annotated the DEGs based on whether they fell within ± 250kb of the lead GWAS variant of each of the IOP or POAG loci above. The chromosome positions of the GWAS variants and colocalizing e/sQTL variants were lifted over from GRCh38 to GRCh37 for the annotation of the DEX-responsive genes.

**Supplementary Results**

*Two-day GC Exposure Results*

*GSEA – TM DEGs*

Apart from extracellular matrix-receptor interaction, the other most enriched pathways in TM were

followed by cell adhesion (GO, adjP=1.49E-17), signaling by GPCR (Reactome adjP=5.01E-08), PI3K-Akt signaling pathway (KEGG adjP=5.79E-04), and Wnt signaling pathway (KEGG adjP=0.001) (**Figure 3A**). Additionally, from the GO database, regulation of multicellular organismal process (BP, adjP=2.81E-16) and cell migration (BP, adjP=2.64E-12) were other top ranking functional groups (**Supplementary Table 8**). A readme (GOFigure_GO_README) tab explaining the GOFigure data is provided in the Supplementary Table Excel File.

*GSEA – SCE DEGs*

Apart from signal receptor binding and extracellular matrix-receptor interaction, the other most enriched pathways in SCE were Interleukin-4 and Interleukin-13 signaling (adjP=4.46E-06) and regulation of insulin-like growth factor (IGF) transport and uptake by IGFBPs (adjP=9.38E-05). Additionally, from the GO database, similar to TM enrichment, regulation of multicellular organismal process (BP, adjP=5.08E-31), cell migration (BP, adjP=7.64E-27), and cell adhesion (BP, adjP=3.81E-26) were other top ranking functional groups for SCE cells (**Figure 3B**). Some of the pathways only found significant in SCE and not in TM after 2day DEX exposure included vasculature development (GO BP adjP=3.18E-21), vascular smooth muscle contraction (KEGG adjP=0.0048), and response to hypoxia (GO BP adjP=3E-07) (Blue outlines in (**Figure 3B**). **Figure 3C** shows the significantly enriched pathways in both TM and SCE after 2-day DEX exposure. Importantly, as may be expected, the DEGs in both TM and SCE cells were enriched in the response to glucocorticoid (GO BP adjP<4.4E-04) and cellular response to corticosteroid stimulus (GO BP adjP<2E-04). In TM and SCE, Spi-B (Spi-B Transcription Factor, adjP=2.51E-08) and MAZ (MYC Associated Zinc Finger Protein, adjP=3.52E-07) were the top-ranking transcription factors, respectively, whose transcription factor binding site (TFBS) were enriched among the DEX DEGs.

*Inspecting association of DEX-responsive genes with IOP or POAG loci via eQTLs and sQTLs*

To further narrow down a high confidence set of DEX-responsive genes whose gene expression changes may be associated with IOP or POAG, we tested whether the 2-day TM exposure DEGs are target genes of eQTLs or sQTLs from GTEx or retina tissues that colocalized with an IOP or POAG GWAS locus based on our previous analysis.^10^ Twenty-five (2.9%) and 52 (2.5%) of the TM and SCE DEGs, respectively, were proposed to be associated with IOP and/or POAG based on eQTLs and/or sQTLs.

Among the DEGs in TM, *VCAM1* was the most significant DEG that also showed evidence of colocalization between its eQTL and an IOP or POAG GWAS locus and whose direction of expression change in response to DEX was consistent with POAG risk, as predicted by the eQTL. An eQTL acting on *VCAM1* colocalized with POAG association with lead variant rs12466440^6^ (**Supplementary** **Table 3**). The colocalizing eQTL in four GTEx tissues suggests that increased expression of *VCAM1* in TM cells after the 2-day exposure to DEX may decrease POAG risk (**Supplementary Table 3**).

Among the DEGs in SCE, *AFAP1* was the most significant DEG whose eQTL also colocalized with an IOP or POAG GWAS locus and is in a consistent direction of expression change to the DEX-responsive findings in SCE (**Supplementary Table 5**). An eQTL acting on *AFAP1* colocalized with rs28649910 and rs12507127 loci for IOP,^5^ and the rs938604 locus for POAG ^6^ (**Supplementary Table 5**). There was decreased expression of *AFAP1* in SCE cells after 2-day exposure to DEX. The rs28649910 and rs12507127 alleles that decrease expression of *AFAP1* in two GTEX tissues based on the colocalizing eQTL were found to increase IOP levels. The rs938604 allele that increases POAG risk was suggested to decrease expression of *AFAP1* in four GTEX tissues based on the colocalizing eQTL.

*Six-hour GC Exposure Results*

To examine the short-term effects of DEX on gene expression, four TM and SCE cell strains from the 2-day exposure experiments were exposed to DEX for 6 hours and RNA-seq was performed. All samples passed QC (**Supplementary Figures 8-9, Supplementary Table 39**).

*6-hr DEX-Induced Differential Gene Expression in TM cells*

The genes that were differentially expressed in TM cells after 6 hours of DEX exposure, accounting for the first SV are shown in **Supplementary Figure 12A**. In the TM, a total of 248 genes were differentially expressed (FDR<0.05) with 186 being up-regulated and 62 being down-regulated (**Supplementary Table 40**). Full results for all the DEGs are shown in **Supplementary Table 41**.

*GSEA of 6-hr DEX-responsive DEGs in TM*

The GSEA for the 248 6-hour DEX-exposed DEGs are shown in **Supplementary Figure 13** and **Supplementary Tables 42 - 52.** From non-redundant GO terms, response to stimulus (BP, 1.97E-27), cell communication (BP, 2.08E-24), cell periphery (CC, adjP=2.40E-18), and binding (MF, adjP=4.00E-23) were top ranked pathways. Enrichment for biological pathways identified signaling by interleukins (adjP= 8.20E-09) and cytokine signaling in immune system from Reactome (adj=8.20E-09) as the top ranked pathways. Transcription factor Spi-B (adjP= 3.3E-18) was identified as the top-ranking transcription factor whose TFBS were enriched among the 6-hours DEX DEGs, as with the 2-day DEX DEGs.

*6 Hour DEX-Induced Differential Gene Expression SCE cells*

The genes that were differentially expressed in SCE cells after 6 hours of DEX exposure with one SV are shown in **Supplementary Figure 12B.** A total of 3,208 genes were differentially expressed with 1482 being up-regulated and 1726 being down-regulated (**Supplementary Table 54**); full results for all the DEGs are shown in **Supplementary Table 55**.

*GSEA of 6-hr DEX-responsive DEGs in SCE*

The GSEA for the 3,208 6-hour DEX-exposed DEGs are shown in **Supplementary Figure 14** and **Supplementary Tables 54 - 61.** From non-redundant GO terms, cellular response to chemical stimulus (BP, adjP=3.81E-37), signaling (BP, adjP=3.03E-34), cell periphery (CC, adjP=1.30E-23), plasma membrane (CC, adjP=1.67E-22) and cell receptor binding/activity (MF, adjP=5.56E-07;adjP=1.81E-06) were top-ranked pathways. Enrichment for biological pathways identified signal transduction (adjP= 3.32E-11) and Interleukin-4 and Interleukin-13 signaling from Reactome (adj=2.76E-09) as the top-ranked pathways. Transcription factor TIEG1 (adjP= 8.94E-21) and MOVO-B (adjP= 8.94E-21) were identified as the top-ranking transcription factors whose TFBS were enriched among the 6-hours DEX DEGs. From the Hallmark GSEA, TNFA_SIGNALING_VIA_NFKB (adjP=1.42E-12) was the top-ranking pathway.

*Overlap between DEGs in TM and SCE cells after 6-hour DEX exposure*

Two hundred and two genes were differentially expressed in both SCE and TM cells, and the distributions of the fold-changes in both cell lines were not statistically different (Wilcoxon rank sum; p-value= 0.1046, **Supplementary Figure 13**). These significant genes are annotated in **Supplementary Table 62.**

*One-hour GC Exposure Results*

To examine the immediate effects of DEX on gene expression, three TM and three SCE cell strains from the 2-day exposure experiments were exposed to DEX for 1 hour followed by RNA-seq. All samples passed QC (**Supplementary Table 53, Supplementary Figure 16**).

*1 Hour DEX-Induced Differential Gene Expression in TM and SCE cells*

The genes that were differentially expressed in TM cells after 1-hour of DEX exposure accounting for the first two SVs are shown in **Supplementary Figure 16A.** A total of 33 genes were differentially expressed with 23 being up-regulated and 10 being down-regulated (**Supplementary Table 63**); full results for all the DEGs are shown in **Supplementary Table 64**. The genes that were differentially expressed in SCE 1 hour of DEX exposure, accounting for the first SV are shown in **Supplementary Figure 16B.** A total of 55 genes were differentially expressed with 34 being up-regulated and 21 being down-regulated (**Supplementary Table 65**), and full results for all the DEGs are shown in **Supplementary Table 66.**

*Overlap between DEGs in TM and SCE cells after 1 hour DEX exposure*

Eighteen genes were differentially expressed in both SCE and TM cells, and the distributions of the fold-changes in both cell lines were not statistically different (Wilcoxon rank sum; p-value= 0.1687, **Supplementary Figure 17**). These genes are annotated in **Supplementary Table 67.**

*GSEA of 1 Hour DEX-responsive DEGs in TM and SCE*

*TM Cells after 1 Hour DEX Exposure*

The GSEA for the 33 DEGs after 1 hour DEX exposure in TM cells are shown in **Supplementary Figure 16**A and **Supplementary Tables 68 - 71**. Enrichment from KEGG database identified IL-17 signaling pathway (adjP=4.11E-07) and TNF signaling pathway (adjP=2.14E-06) as the top ranked biological pathways. From non-redundant GO terms, regulation of multicellular organismal development (BP, adjP=1.59E-09), nucleus (CC, adjP=0.0007315) and RNA polymerase II cis-regulatory region sequence-specific DNA binding, DNA-binding transcription activator activity, and RNA polymerase II-specific and kinase binding (MF, adjP=1.05E-03) were top ranked processes. Transcription factor SRF (adjP=0.004) was identified as the top-ranking transcription factor whose TFBS were enriched among the 1-hour DEX DEGs in TM.

*SCE Cells after 1 Hour DEX Exposure*

The GSEA results for the 55 DEGs are shown in **Supplementary Figure 16B** and **Supplementary Tables 75 - 81.** Enrichment from Reactome database identified NGF-stimulated transcription (adjP=1.67E-05) and response to metal ions (adjP=1.6E-04) as the top ranked biological pathways. From non-redundant GO terms, multicellular organismal process, response to stimulus, negative regulation of biological process, and vasculature development (BP, adjP<9E-07) were top ranked processes. Nucleus, cellular anatomical entity and cytoplasm (CC, adjP, 3.16E-05) and protein binding (MF, adjP, 0.01) were among the top ranked cellular components and molecular functions, respectively. Enrichment of TFBS identified CPBP and NF-kappaB (adjP=0.003) among the top-ranking transcription factor whose TFBS were enriched among the 1-hour DEX DEGs in SCE.

*Quantitative Polymerase Chain Reaction (qPCR) Confirmation of RNA-seq Results*

We chose eleven and ten genes from the TM and SCE RNA-seq experiments, respectively, to validate the DEG results with qPCR (**Supplementary Table 84-85**). The TaqMan probes and the GenBank Accession numbers for each gene expression assay are shown in **Supplementary Table 86.** For all genes tested the direction of differential expression change was consistent between qPCR and RNA-seq across all TM and SCE samples except for one instance of an opposite expression pattern for sample TM137 and the gene *NR3C1*. The disparity of *NR3C1* expression between qPCR and RNA-seq for sample TM137 seems to be biological in nature, as TM137 does not appear to be an outlier based on correlation and clustering analysis of all TM biological replicates using transcript per million (TPM) across all genes (**Supplementary Figures 1 and 2**) or only the qPCR tested genes (**Supplementary Figure 20**). The correlation between the RNA-seq fold-change from DESeq2 and relative quantitative (RQ) expression values from qPCR is shown in **Figure 8** for the 2-day DEX exposure TM and SCE samples. There was high correlation between the average differential expression measured by qPCR and RNA-seq for the tested genes, with a Pearson correlation coefficient (r) of 0.94 and 0.98 for TM and SCE, respectively (**Figure 8**).
